# Targeted anticancer pre-vinylsulfone covalent inhibitors of carbonic anhydrase IX

**DOI:** 10.1101/2024.05.20.594908

**Authors:** Aivaras Vaškevičius, Denis Baronas, Janis Leitans, Agnė Kvietkauskaitė, Audronė Rukšėnaitė, Elena Manakova, Zigmantas Toleikis, Algirdas Kaupinis, Andris Kazaks, Marius Gedgaudas, Aurelija Mickevičiūtė, Vaida Juozapaitienė, Helgi B Schiöth, Kristaps Jaudzems, Mindaugas Valius, Kaspars Tars, Saulius Gražulis, Franz-Josef Meyer-Almes, Jurgita Matulienė, Asta Zubrienė, Virginija Dudutienė, Daumantas Matulis

## Abstract

We designed novel pre-drug compounds that transform into an active form that covalently modifies particular His residue in the active site, a difficult task to achieve, and applied to carbonic anhydrase (CAIX), a transmembrane protein, highly overexpressed in hypoxic solid tumors, important for cancer cell survival and proliferation because it acidifies tumor microenvironment helping invasion and metastases processes. The designed compounds have several functionalities: 1) primary sulfonamide group recognizing carbonic anhydrases (CA), 2) high-affinity moieties specifically recognizing CAIX among all CA isozymes, and 3) forming a covalent bond with the His64 residue. Such targeted covalent compounds possess both high initial affinity and selectivity for the disease target protein followed by complete irreversible inactivation of the protein via covalent modification. Our designed prodrug candidates bearing moderately active pre-vinyl sulfone esters or weakly active carbamates optimized for mild covalent modification activity to avoid toxic non-specific modifications and selectively target CAIX. The lead inhibitors reached 2 pM affinity, highest among known CAIX inhibitors. The strategy could be used for any disease drug target protein bearing a His residue in the vicinity of the active site.

## INTRODUCTION

Covalently binding compounds performing targeted covalent inhibition (TCI) (1–4) bear special functional groups called “warheads”, usually Michael acceptors. Initially, these compounds bind to the target protein reversibly via their specific structural features that recognize the target protein (5). In a subsequent step, an irreversible covalent bond is formed between the “warhead” fragment and the targeted amino acid, mostly a nucleophilic one like cysteine (5). This irreversible mode of binding provides a prolonged mechanism of action, full and reversible or irreversible target inactivation, the need for lower drug dosages, the opportunity for higher selectivity toward the target, and in some cases – effective inhibition of drug-resistant enzyme mutants (5–7). Among successfully applied targeted covalent inhibition examples are ibrutinib (BTK inhibitor), afatinib (EFGR T790M mutant inhibitor), osimertinib (improved EFGR T790M mutant inhibitor), and sotorasib (KRAS G12C mutant inhibitor), which are already approved by the Food and Drug Administration (8, 9).

Here we introduce previnylsulfone warhead for TCI by the formation of covalent bond with the protein histidine residue. In contrast to cysteine, the intrinsic nucleophilicity of histidine is weaker and there are few reports about histidine labeling in proteins by small molecules, most of them being highly reactive and therefore not usable as warhead in drug development due to non-specific reactions (10–13). Known histidine-targeting compounds mostly carry a highly reactive warhead, such as sulfonylfluorides, which provide some degree of selectivity due to suitable substituents, which interact selectively with their target protein (14, 15), but it is almost impossible to prevent non-specific labeling. Chemoselective modification of histidine is very difficult to achieve. A light-promoted and radical-mediated selective C-H-alkylation of histidine for peptide synthesis has been suggested (16), but is not applicable to proteins. Another method for chemoselective histidine bioconjugation uses thiophosphorodichloridate reagents, which mimic naturally occurring histidine phosphorylation (10). A light-driven selective approach for labeling histidine residues in native biological systems was developed with thioacetal as thionium precursor (17). However, the light-driven approach requires the presence of high concentrations of Rose Bengal as catalyst, not suitable for therapeutic applications.

We applied the TCI strategy to human carbonic anhydrases (CA), metalloenzymes that catalyze reversible CO_2_ hydration by producing acid proton and bicarbonate anion. There are 12 catalytically active CA isozymes in humans. The CAI, II, III, VII, and XIII are cytosolic, CAIV is membrane-bound, while CAVA and VB are found in mitochondria. CAVI is the only secreted isozyme found in saliva and milk, whereas CAIX, XII, and XIV are transmembrane proteins bearing extracellular catalytic domains (18). Catalytic domains of CA isozymes are highly homologous and bear structurally very similar beta-fold. However, the isozymes differ in their enzymatic activity, tissue distribution, and cellular localization.

CAIX is a hypoxia-inducible protein, that participates in cancer cell proliferation and metastasis (19, 20). As recently demonstrated by proteomic analysis, CAIX interacts with amino acid and bicarbonate transporters to control cancer cell adhesion, a critical process involved in migration and invasion. The CAIX also plays an important role in the migration of cancer cells by interaction with collagen-, laminin-binding integrins, and MMP-14 (21, 22). It is proposed that targeting CAIX catalytic activity and/or interrupting the interactions with metabolic transport proteins and cell adhesion/migration/invasion proteins will have therapeutic benefits by involving pH regulation, metabolism, invasion and metastasis(22, 23).

Sulfonamides stand out as the most extensively researched class of carbonic anhydrase (CA) inhibitors (24, 25). They exhibit a high binding affinity in their deprotonated state to the zinc-bound water form of CA (26–28). Drugs used in the clinic as CA inhibitors, such as acetazolamide, methazolamide, dichlorophenamide, dorzolamide, and brinzolamide, have various side effects (29). The development of isozyme-selective CA inhibitors is a major goal of drug discovery. Any such drugs will be more beneficial than the currently available mostly non-selective CA inhibitors, as the reduction of side effects will improve the effectiveness of the therapy.

Covalent inhibitors have been previously designed for CAI and CAII, namely, bromoacetazolamide and N-bromoacetylacetazolamide (30–32). Although N-bromoacetylacetazolamide formed a covalent bond with CAI His67 and bromoacetazolamide with CAII His64 amino acids, further research of these compounds was discontinued (32). Recently, studies involving covalent modification of CA isozymes (mainly CAII) have been published. However, the synthesized molecules were designed not to act as enzyme inhibitor, but as a model protein to investigate benzenesulfonamide-bearing fluorescent label and a warhead able to bind the enzyme covalently (29). One of their new probes bearing an epoxide reactive group was not only able to form a covalent bond with the protein, but it did it selectively for His64 (33). Similarly, analogous sulfonamides without fluorescent groups were synthesized and formed the covalent bonds with His3 or His4 of CAII (34, 35). Different warheads were applied to react with His64 and His3 (36, 37) bearing S(IV) fluoride to present a new way for the expansion of the liganded proteome (36, 37).

In this work, we investigated fluorinated benzenesulfonamide compounds bearing sulfonylethyl ester and sulfonylethyl carbamate moieties as possible covalent CA inhibitors. The 3-substituted-2-((2,5,6-tetrafluoro-4-sulfamoylphenyl)sulfonyl)ethyl acetate exhibited surprisingly high binding affinity for CAIX, which was more than ten fold higher than our previous synthesized lead compound VD11-4-2 (*K*_d(CAIX)_= 32 pM) (38). The MS and X-ray crystallography data confirmed the covalent binding of new compounds to the proton shuttle His64 residue. We showed that sulfonylethyl ester/carbamate behaves as a prodrug by reacting with His64 in the active site of CAs through the elimination mechanism to release ester or carbamate moieties, thus forming a reactive vinylsulfone group. The newly discovered mechanism of inhibition of CA through the formation of a covalent bond between the compound and the protein has great potential for the development of high-affinity compounds for a particular CA isozyme, and such compounds could become precursors of new-generation drugs.

## RESULTS

### Design and mechanism of covalently-binding CA inhibitors

In search of high-affinity and high-selectivity inhibitors of carbonic anhydrases (CA), a series of fluoro-benzenesulfonamide-based compounds were synthesized (Figure 1, schemes S1-S3). The benzenesulfonamide group is important for making a coordination bond with the Zn(II), while the fluorine atoms were included to withdraw electrons from the sulfonamide group and diminish its p*K*_a_ in order to strengthen interaction. Surprisingly, part of the compounds exhibited extremely strong binding affinity. These compounds contained the -SO_2_CH_2_CH_2_OCOR group that was forming an unexpected covalent bond with the protein.

**Figure 1.**
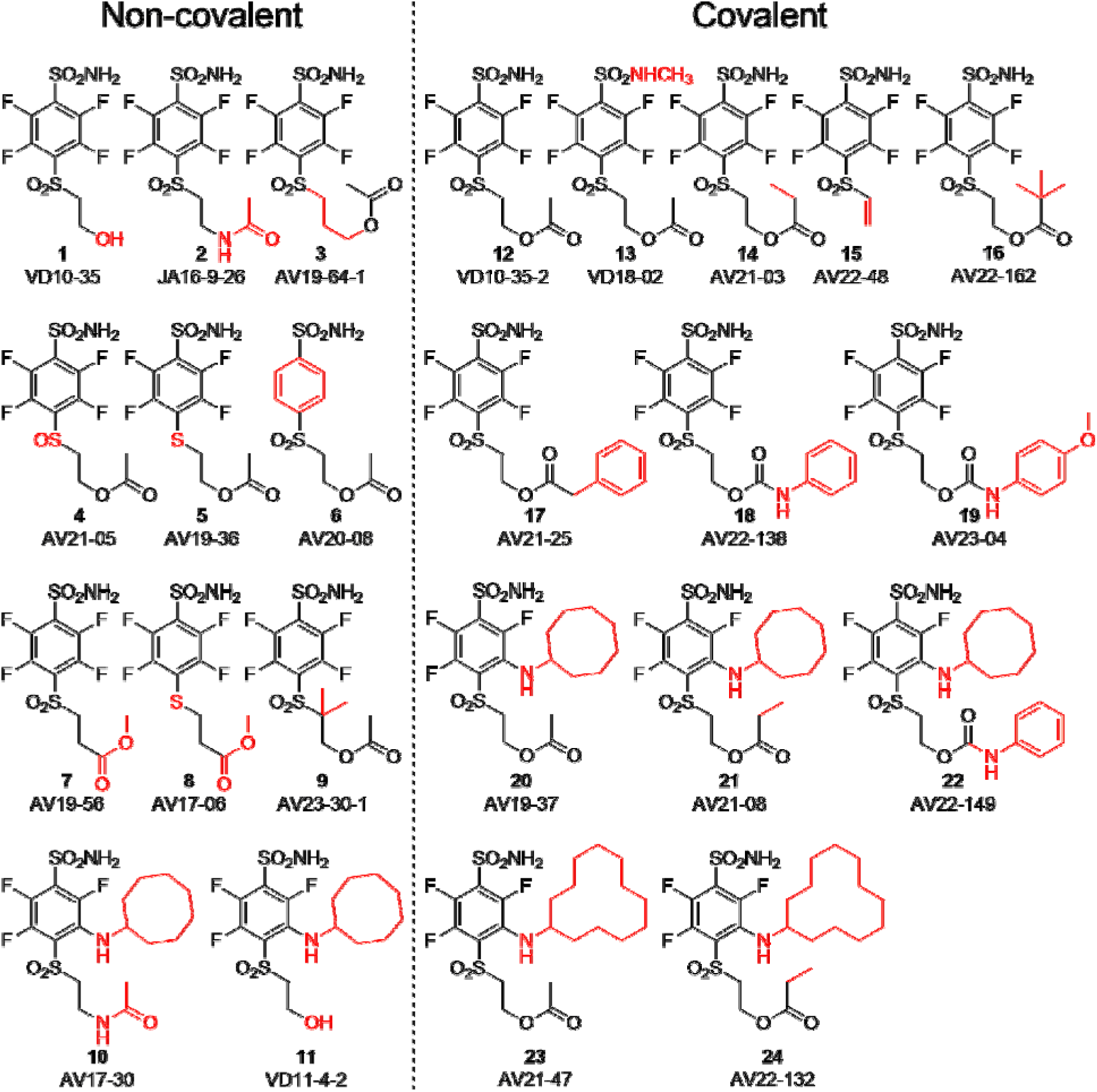
Chemical structures of compounds used in this study and designed to investigate the covalent bindin capability to CA proteins. Compounds on the left of the vertical dashed line do not form the covalent bond, while the ones on the right form the covalent bond with the protein molecule. Moieties shown in red are important for structural comparison to visualize the chemical groups that are responsible for covalent interaction, high affinity, or high selectivity for CAIX.

It has been known in organic chemistry that compounds bearing this fragment in the presence of bases can rearrange to vinyl-sulfonyl moiety which has been reported as a covalently-modifying “warhead” (39–41). To the best of our knowledge, this kind of rearrangement/elimination has not been applied for enzyme covalent inhibitors. We designed benzenesulfonamides with the SO2CH2CH2OCOR group that can form highly reactive electrophilic species without adding additional base by adding multiple fluorine atoms to the benzene ring. Furthermore, the rearrangement occurred only in the carbonic anhydrase enzyme active site. The active vinyl-sulfonyl group formed via elimination reaction, which occurred easily due to strong electron withdrawing effect of fluorines on benzene ring (Figure 2).

**Figure 2.**
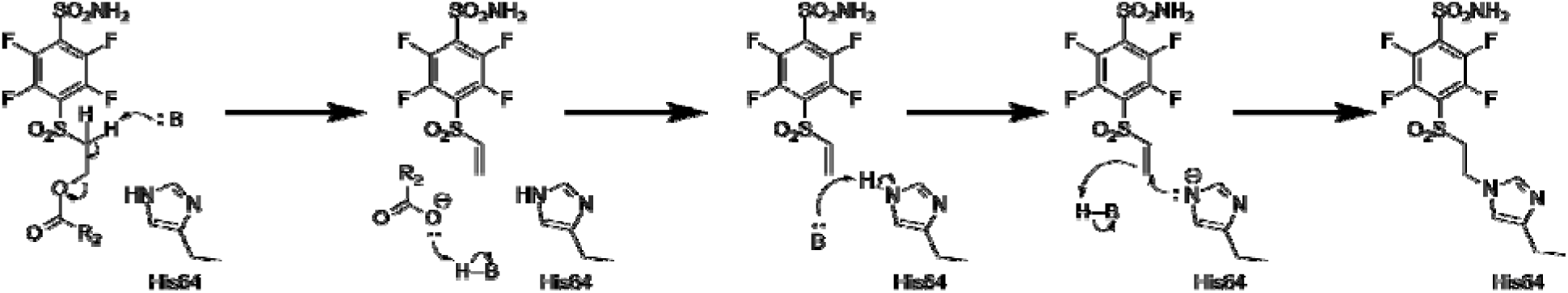
Proposed rearrangement mechanism (beta-elimination) of compounds bearing the -SO_2_CH_2_CH_2_OCOR fragment to vinyl-sulfone and the formation of covalent bond with the histidine (His64) amino acid side chain of the CA protein.

The SO_2_ group at the *para* position relative to the sulfonamide group was also necessary for a covalent bond because compounds **4**, **5** bearing the SO or S groups, respectively, did not form the covalent bond with CA isozymes. Moreover, the change of the -O-CO-R group to -NH-CO-R or -CO-O-R also prevented the formation of the covalent bond as illustrated by compounds **2**, **10**, **7**, and **8** that did not form covalent bond with the protein molecule.

We propose the mechanism where the covalent modification occurs via the elimination mechanism shown in Figure 2, where a basic amino acid residue removes the proton and formed the vinyl sulfone moiety, which only then forms a covalent bond with the nitrogen atom of the histidine residue. To check this mechanism, two control compounds, **3** and **9**, were synthesized. Both these compounds did not form a covalent bond with the protein (Figure S13 and S14). In compound **9**, two methyl groups located at the crucial α carbon replaced the proton that needed to be removed. In compound **3,** the third methylene group prevented the β-elimination reaction. Furthermore, the concept of formation of vinyl sulfone fragment was demonstrated by synthesizing compound **15** bearing the vinyl sulfone itself. This compound readily formed the covalent bond with proteins (Figure S15).

Since it appeared that compounds bearing ester groups (e.g. compounds **17** and **20**) may have too high rate of covalent bond formation leading to non-desired modification of non-targeted proteins, we have designed and synthesized compounds bearing the carbamate functional group (e.g. compounds **18** and **22**). The carbamate group was more stable than the ester and thus the covalent modification reaction of the protein showed milder rate than the ester.

The cyclooctyl or cyclododecyl rings present in covalent compounds **20**-**24** and in non-covalent compounds **10**-**11** have been previously designed by our group to specifically bind to CAIX isozyme and intended not to bind to the other eleven human catalytically active CA isozymes. The ring could fit to the pocket in CAIX, but not so readily to other isozymes (38, 42). The three-way recognition model is shown in Figure 3.

**Figure 3.**
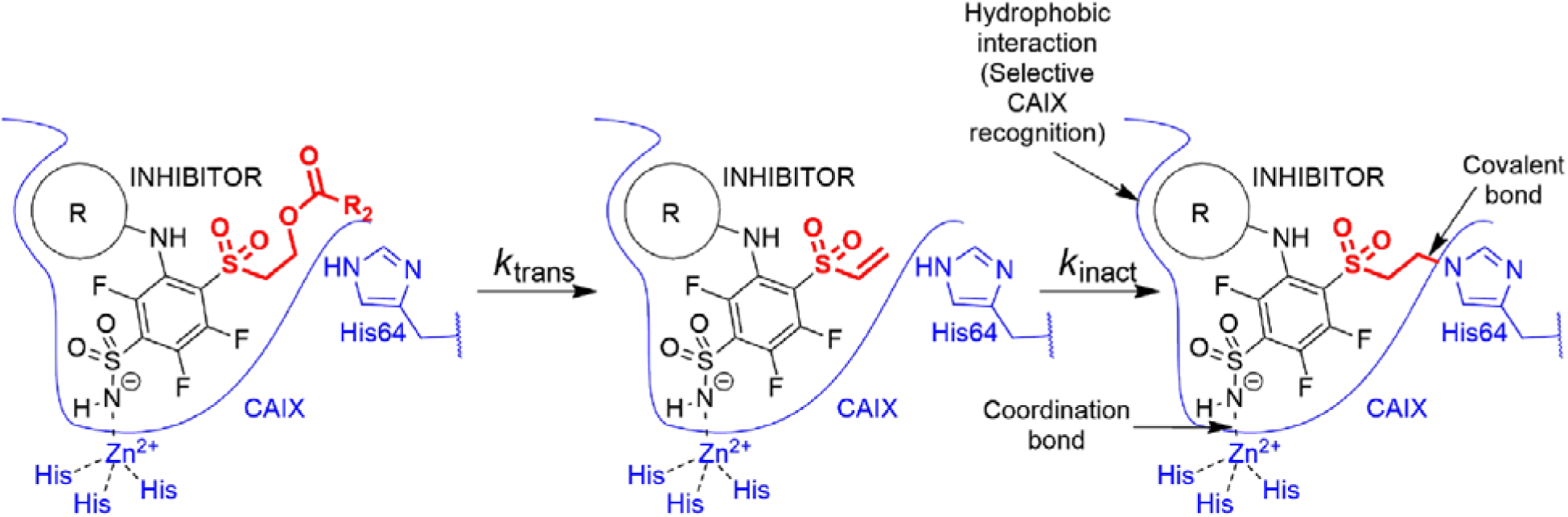
Transformation of pre-drug to the active vinyl sulfone and a three-way recognition of CA isozymes. First, the negatively charged sulfonamide forms coordination bond with the Zn(II). Second, hydrophobic cyclooctyl rin fits to the hydrophobic pocket of CAIX isozyme and provides substantial selectivity over other CA isozymes. Third, covalent bond forms with the histidine providing irreversible inhibition of CAIX enzymatic activity.

### Covalent interaction between inhibitors and CA isozymes byX-ray crystallography

The covalent bond between the compounds and protein was demonstrated by X-ray crystallography (Figure 4). The crystal structures of compounds **21**, **20**, and **23** with CAI, CAII, and CAIX, respectively, showed covalent bond formation between the ligands and the histidine residue of the enzymes. In CAI, the ligand forms a covalent bond with His 67, while in CAII and CAIX, the bond is made with His 64 – a residue responsible for proton shuttle function in CA isoforms (43). The distance between the nitrogen atom of histidine and the vinyl carbon atom of the compounds was around 1.5 Å, consistent with the length of the covalent bond and was clearly visible with strong electron density in all three crystal structures (Figure 4A, 4B, and 4C).

**Figure 4.**
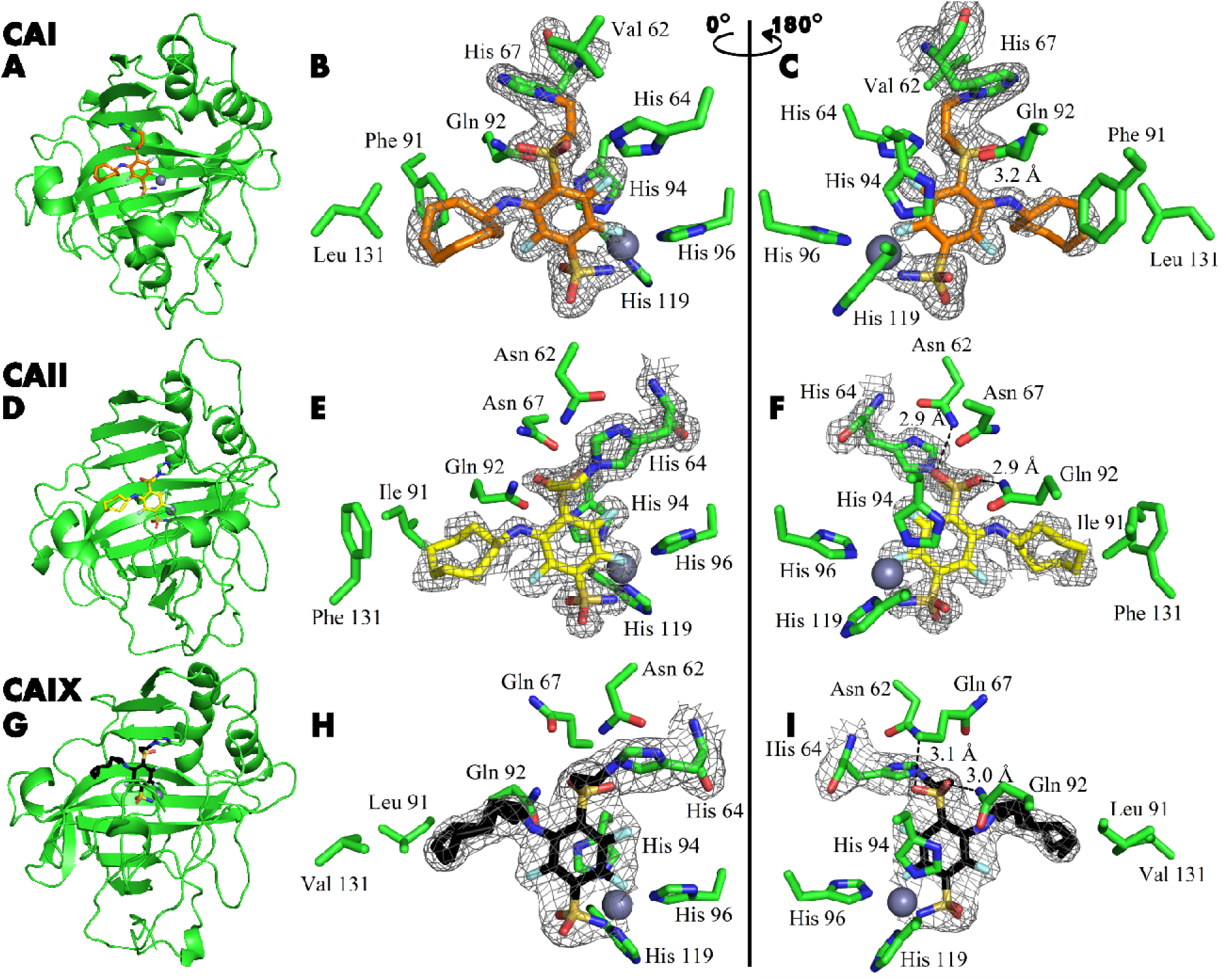
X-ray crystal structures of CAI (**A**-**C**), CAII (**D**-**F**), and CAIX (**G**-**I**) covalently bound with inhibitors **21**, **20**, and **23**, respectively. Left panels show cartoon models of the entire protein molecule with the covalently bound compound and the His64 residue shown as stick model, while the middle and right panels show close-up views of the inhibitor, shown as sticks, displayed with a 180° rotation between the images. The 2Fo-Fc map is shown only for the ligand and the histidine residue with which it forms a covalent bond, contoured at 1σ.

The tail moiety of the ligands is also coordinated by two hydrogen bonds – with Asn 62 and Gln 92 (CAII numbering) in the cases of CAII and CAIX, while in CAI, a possible hydrogen bond is formed with Gln 92 and His 64. Electron density was overall good for the whole ligand in all three crystal structures, with slightly weaker electron density observed in the hydrophobic tail region indicating higher flexibility. The hydrophobic tail part is oriented towards the active site region, which varies most between CA isozymes (the so-called “hot spot” for isozyme (isoform)-selective inhibitor design (44)).

There are two protein subunits in CAI structure, position of the modeled **21** is very similar in both. The electron density of the ligand is better in protein chain B. Sulfonamide group and modification of His67 with para-linker are clearly visible. The fluorine atoms of benzene ring also could be located clearly, but the electron density of benzene ring is partially lost. This could be explained by partial occupancy of the modeled conformation of the inhibitor and by rotation of ligand fraction, since the ligand is probably not fixed by the covalent bond with His67. Cyclooctyl group is visible in both subunits only partially due to flexibility of the ring resulting in multiple conformations. This group is oriented towards the hydrophobic part of the active site, defined by Leu131, Ala135, Leu141, and Leu198.

### Covalent interaction by mass spectrometry and enzymatic activity

The mass spectra of CA isoforms incubated with compound **12** showed a 319 Da shift compared to pure CA isoforms (except CAIII), which is equal to compound **12** molar mass without ester group (Table S2, figure S1-S12). Moreover, after a 3-minute incubation of 1:1 molar ratio of compound **12** with CAXIII protein (29575 Da), nearly half of the protein was already covalently modified in this time as shown by additional 29894 Da peak (319 Da shift, Figure 5A)). After 2-hour incubation, essentially entire protein fraction was covalently modified by the compound. The presence of the minor peak at 30213 Da (638 Da shift) in Figure 5A indicates that there is a second modification site on CAXIII protein. The peptide mapping by digestion of the CAXIII with thrombin detected the compound **12** covalently bound exclusively to the peptide containing His64 residue (Figure S21 and S22). However, in the ^1^H-^15^N-HSQC 2D NMR spectrum, upon incubation of CAII isozyme at 1:1 molar ratio for one hour with covalent compound **12**, in addition to the His64 signal change, we observed a decrease of peak intensity in the N-terminal part of CAII indicating an additional minor fraction of enzyme with covalently bound **12** outside CA active site (Figure S23 and S24). Thus, ester compound **12** may be too reactive for fully specific inhibition. The non-covalent compound **6** was incubated with CAII at 10-fold surplus of the compound, but no covalent modification was detected (Figure S16).

**Figure 5.**
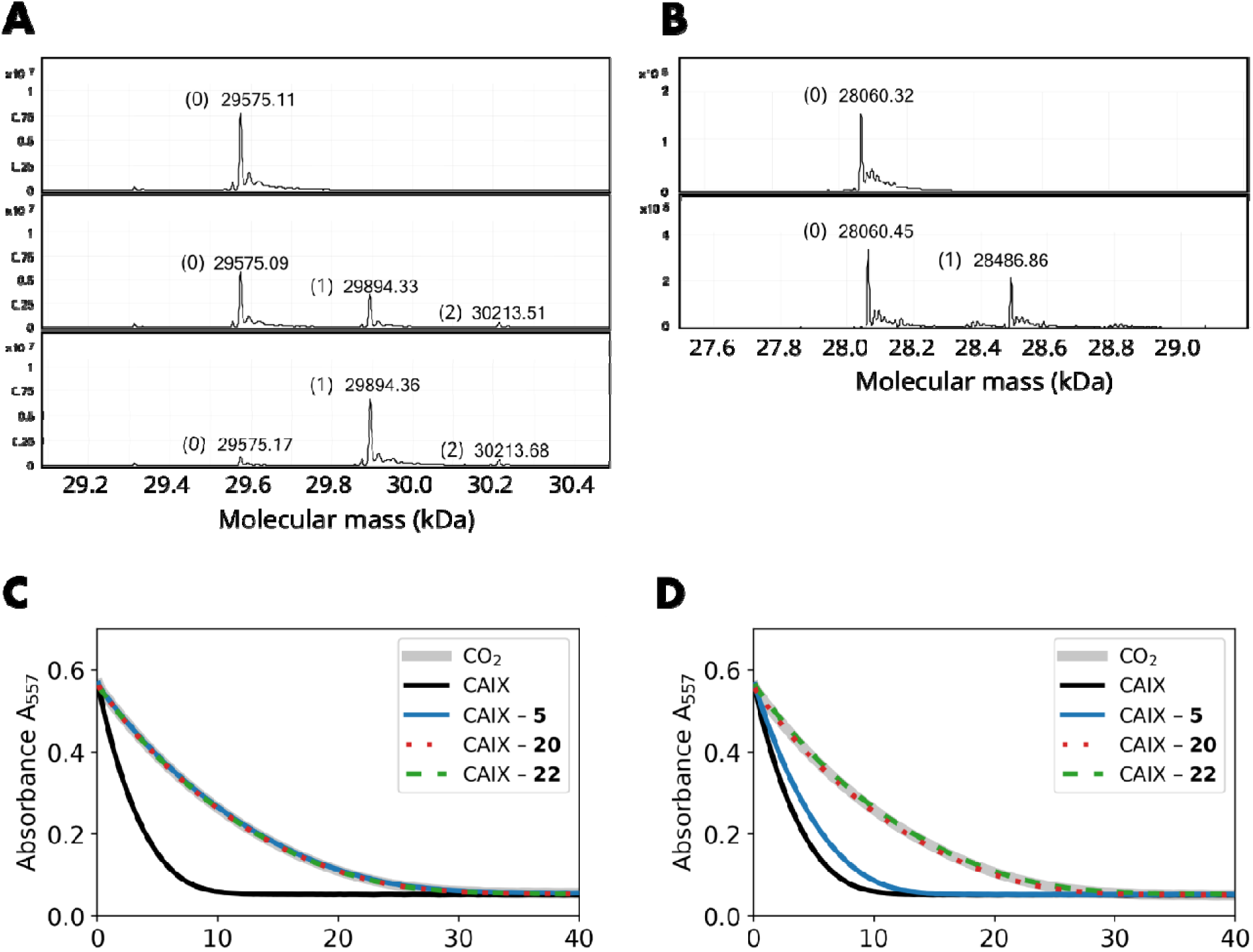
Covalent interaction shown by HRMS, and enzymatic activity recovery assay. **A**. MS spectra of CAXIII in th absence of compound (top panel), the presence of 1:1 molar ratio of compound **12** after 3-minute incubation (middle panel) and 2-hour incubation (bottom panel). **B**. MS spectra of CAIX in the absence of compound (top panel) and after the incubation of 1:1 molar ratio of compound **22** (carbamate) for 4 hours. **C**. Enzymatic activity of CAIX before dialysis, while **D** – after dialysis (32 hours, 4-times buffer change, black solid line – fully active CAIX, grey solid line – spontaneous CO_2_ hydration reaction (coincides with fully inhibited CAIX), blue solid line – CAIX with non-covalent **5**, dotted red line – CAIX with covalent **20**, and dashed green line – CAIX with covalent carbamate **22**. The recombinant CAIX recovered almost full activity after dialyzing out the non-covalent compound, while th activity remained fully inhibited with the covalent compounds.

Therefore, a series of compounds bearing the carbamate leaving group were designed (**18**, **19**, and **22**) to reduce chemical reactivity and reduce undesired reactions with non-intended groups. Compound **22** exhibited covalent modification of CAIX, but after 4-hour incubation there was still significant part of non-modified protein present. Thus the reaction was significantly slower than with the ester group. The MW of CAIX as calculated from the sequence was 28061.68 Da and matched closely the measured mass of 28060.32 (Figure 5B top panel and Figure S20) or 28060.45 (lower panel). The calculated MW of compound **22** was 563.1372 Da and was measured to be 564.1444 Da. The calculated mass of compound **22** without the carbamate leaving group was 426.0895 Da, while the measured difference in Figure 5B was 426.41 Da, a perfect match. Thus compound **22** exhibited both highly specific and relatively slow modification of CAIX, with a good perspective toward drug design.

The covalent irreversible and non-covalent reversible interaction was also demonstrated by comparing the inhibition of enzymatic activity of CAIX by non-covalent compound **5** and covalent compound **20** and their possibility to be dialyzed out. Both compounds fully inhibited the enzymatic activity of CAIX at 1:1 molar ratio in the same dose-dependent manner. The resultant protein-ligand complex was then subjected to 32-hour dialysis. The CAIX complex with the non-covalent **5** regained 73% of the original enzymatic activity, while the CAIX with covalent **20** did not regain any detectable enzymatic activity (Figure 5C and 5D). This indicates that the covalent modification irreversibly inhibited the enzymatic activity of CAIX.

To determine the contribution of the primary sulfonamide group on the capability of making a covalent bond with CA, we synthesized a secondary sulfonamide **13** and compared with the analogous covalent compound **12**. After 2 hour incubation at 10:1 molar surplus of secondary sulfonamide **13**, the free CAII protein still dominated, indicating that only a minor fraction of the protein was covalently modified (Figure S17B). In comparison, using the same conditions, compound **12** bearing the primary sulfonamide group completely modified CAII (Figure S17C).

This shows the significant effect of the primary sulfonamide group in guiding the compound into the CA active site and consequent covalent bond formation with His64 amino acid. In the absence of the guiding sulfonamide group, such as in **13**, a relatively slow modification most likely occurred on nucleophilic residues different than the His64 in the protein active site. This unintended covalent modification by the secondary-sulfonamide **13** was also observed with isozymes other than CAII. Although this compound should have low affinity to all CA isozymes, it still modified CAXIII at a 10:1 compound surplus molar ratio after 2-hour incubation (Figure S18). Despite that, it is important to note that the modification most likely occurred at a different nucleophilic amino acid, not the His64 in the active site.

The presence of non-specific unintended covalent modifications prompted us to synthesize different covalently modifying groups that would be less reactive and more suitable for drug design, such as carbamate compounds **18** and **19**, which showed significantly slower (at least by two orders of magnitude) covalent-modification activity compared to the ester compounds (Figure S19). Even using less reactive carbamates, CA isozymes were still able to make covalent bonds with more than one inhibitor molecule albeit in much lower quantity.

### Specific binding of covalent compounds to CAIX expressed on live cell surface

The HeLa cell culture was grown under hypoxia and shown to express CAIX on the cell surface reaching the concentration of 2-10 nM, determined by saturating with fluorescein-labeled compound GZ19-32 as previously described (45). Covalent compounds were added to the cell culture at various concentrations together with 10 nM of GZ19-32 that strongly and specifically binds CAIX. The tested covalent compounds competed for the binding to the CAIX active site in a dose-dependent manner (Figure 6). At high concentrations (e.g. 100 µM, 10,000-fold surplus over CAIX and GZ19-32), the compounds completely outcompeted the CAIX-specific GZ19-32. However, at low concentrations, around 10 nM, the compounds competed with GZ19-32 depending on the compound’s chemical nature. Thus, the covalent compounds were available for binding to CAIX and, most likely, did not bind to other proteins that are expected to be present in abundant quantities on the cell surface. These other proteins certainly have His residues that would have been modified if the non-specific binding occurred.

**Figure 6.**
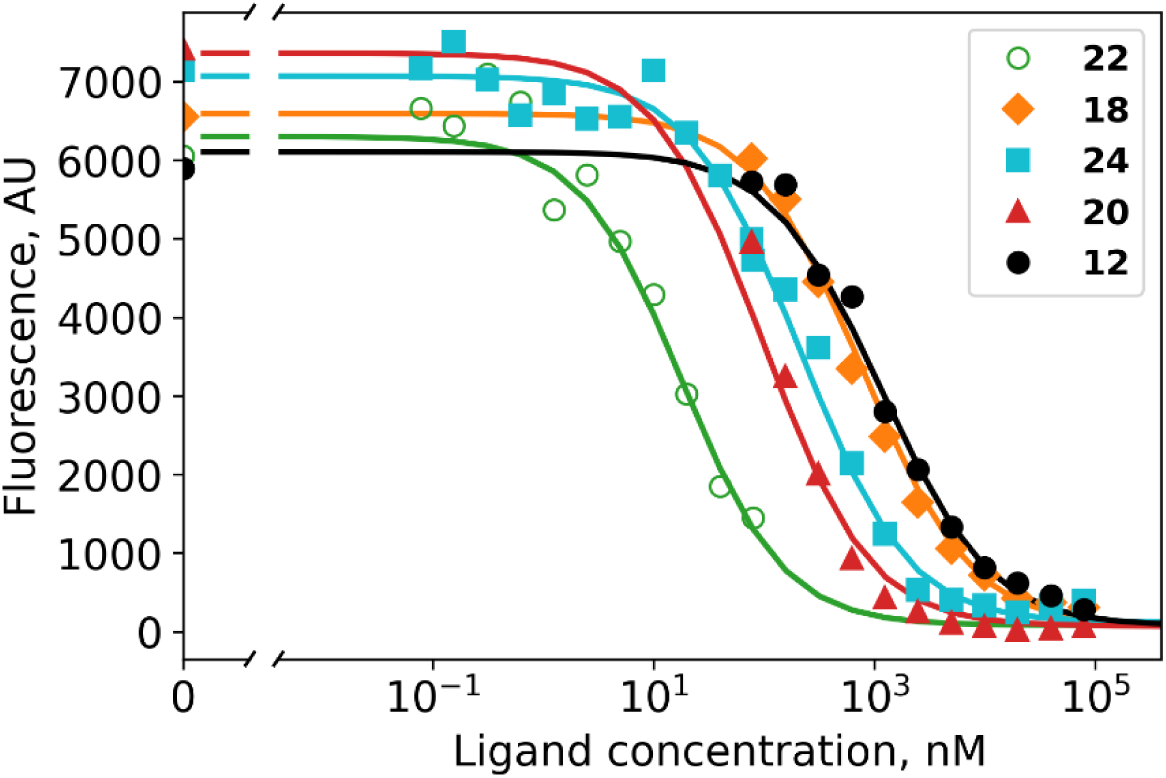
Dosing curves of covalent compounds applied to hypoxic live cell culture expressing CAIX: **22** – green; **18** - orange; **24** – cyan; **20** – red and **12** – black. The compounds competed with the fluorescein-labeled GZ19-32, added at 10 nM concentration to all samples. A competitive binding model applied to obtain the affinities of teste compounds for cell-surface CAIX (Table 1).

**Table 1.** Affinities (apparent dissociation constants K_d,app_) of covalent compound binding to cell-expressed CAIX determined by applying a competitive binding model to data in Figure 6 as previously described (45). Parameters used in the competitive model were: the CAIX protein concentration was 5 nM (P_t_ = 5 nM), the dissociatio constant of GZ19-32 was 150 pM (K_d_B_ = 150 pM), and the concentration of GZ19-32 was 10 nM (L_t_B_ = 10 nM).

| Compound | $K_{d,app\_A}$, nM |
| --- | --- |
| **12** | 22 |
| **18** | 15 |
| **24** | 4.0 |
| **20** | 1.8 |
| **22** | 0.30 |

Several covalent compounds were chosen to demonstrate the importance of compound structural features for CAIX recognition in cell cultures. Two compounds **12** and **18** were para-substituted benzenesulfonamides, non-selective for CAIX. Compound **24** had a cyclododecyl amino substitution, selectively recognizing CAIX, but slightly too-large for optimal binding and solubility. The **20** contains cyclooctylamine substitution at the *meta* position, exhibiting a high affinity for purified CAIX. Finally, **22** beared the cyclooctylamine substitution and the carbamate leaving group optimized for lower covalent modification activity compared to ester.

All tested covalent compounds competed with the fluorescein-labeled GZ19-32 for the binding to cell surface CAIX in a dose-dependent manner. Their apparent dissociation constants, a determined by the competition with GZ19-32, are listed in Table 1. The *para*-substituted compounds that are non-selective for CAIX, bound weaker to cell-surface CAIX than the *meta* substitute-bearing CAIX-selective compounds. The ester compounds bearing both *para* and *meta* substitutions designed for CAIX recognition exhibited single-digit nanomolar affinities (4.0 nM for **24** and 1.8 nM for **20**). However, the carbamate compound **22** exhibited the strongest affinity (300 pM) for cell-surface CAIX among all tested compounds.

The carbamate compound **22** showed the highest affinity for cell-expressed CAIX and irreversibly covalently modified the protein in the active site, thus permanently inhibiting its enzymatic activity. Therefore, this compound is a leader among tested compounds to serve as an anticancer inhibitor of CAIX, highly expressed in hypoxic solid tumors.

### Covalent compound binding apparent affinities to purified CA isozymes

Covalent compounds formed an irreversible covalent bond with the protein molecule. This inhibition mode may occur in two stages. In the first stage, the inhibitor interacts with the enzyme due to its affinity to the targeted enzyme. Here, the affinity is determined by the primary sulfonamide group and the hydrophobic substituent in the *meta* position. The compound is still able to reversibly dissociate and its non-covalent binding affinity is quantified by the dissociation constant *K*_d,_ defined as the ratio of dissociation and association rate constants *k*_off_/*k*_on_:

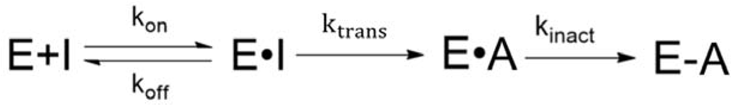

In the second stage, the pre-vinylsulfone compound is chemically transformed into the reactive vinylsulfone electrophile by a basic amino acid of the enzyme at a rate of k_trans_. In the final reaction step, the vinylsulfone may form a covalent bond with the nucleophilic residue with a specific inactivation rate constant *k*_inact_. Since the vinylsulfone is highly reactive, k_trans_ must be rate-limiting and accounting for the apparent inactivation rate, much slower than for vinyl sulfone **15**.

It is obviously incorrect to state covalent compound affinities in terms of a conventional dissociation constant *K*_d_. Therefore, the apparent dissociation constant is valid only to limited extent because if there is an irreversible chemical modification, then eventually all of the protein will be modified independent of the affinity. In our case, we can assume a rapid pre-equilibrium followed by a slow covalent modification. Therefore relative affinity measurements are valid both by competition assay described above and the fluorescence-based thermal shift assay, described below. However, due to the interplay of kinetic and thermodynamic equilibrium contributions, the affinity measurements should still be considered with caution.

We applied the fluorescence-based thermal shift assay to determine the apparent dissociation constants *K*_d,app_ of covalent compounds to arrange them in the order of their apparent affinities (association rate constants) (Figure S25-S33). For example, the CAIX-specific covalent compound **22** bound with an extremely tight affinity, the apparent dissociation constant was determined to be 7.8 pM, the highest affinity among known CAIX-binding compounds. The thermal shift was over 18 °C and exhibited a typical dosing curve often observed for covalent compounds (Figure 7). There was a strong shift of the protein melting temperature caused by the compound but no further shift as observed in reversible non-covalent interactions. Using compound **18**, we determined that the obtained apparent affinity constants were time-independent, meaning lower errors due to compound-protein incubation time and preparation for FTSA (Figure S34-S39).

**Figure 7.**
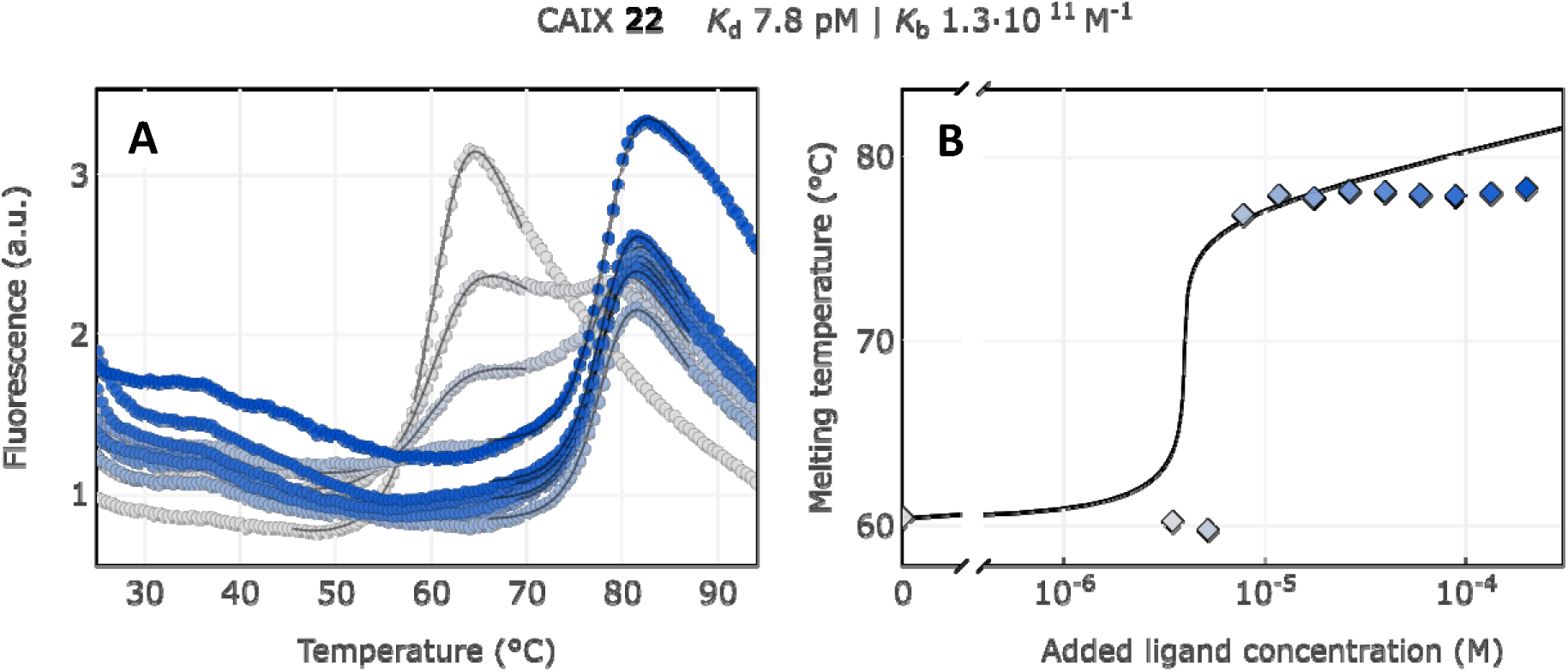
Apparent affinity determination of compound **22** by the thermal shift assay. **A**. Raw FTSA data of compound **22** binding to CAIX (pH 7.0 for 37 °C). **B**. Enzyme melting temperature dependence on compound **2** concentration. Datapoints saturated due to the covalent nature of interaction and therefore did not fully fit to th model line.

The observed *K*_d,app_ values of all covalent compound binding to CA isozymes were higher than those of its noncovalent analogs. For example, CAIX binds to the para-substituted esters **12** and **14**, forming a covalent bond with the protein, up to 1000 times stronger than the noncovalent para-substituted compounds **1** and **3**. Covalent compounds with meta-substituents **20**-**24** bind CAIX up to 10 times more strongly than their noncovalent analogs **10** and **11**. The apparent dissociation constants of covalent compound binding to all 12 human catalytically active CA isozymes as determined by the fluorescence-based thermal shift assay are listed in Table 2. The apparent affinities of covalent compounds can be compared with the non-covalent reversibly-binding analog compounds and estimate the energetic contribution of the covalent bond.

**Table 2.** The apparent dissociation constants K_d,app_ (in nM units) for compound interaction with human recombinant CA isozymes as determined by fluorescence-based thermal shift assay (FTSA) at pH 7.0 for 37 °C. The values ar logarithmic averages of several independent FTSA experiments.

| | Compound | $K_{d,app}$ nM | | | | | | | | | | | |
| --- | --- | --- | --- | --- | --- | --- | --- | --- | --- | --- | --- | --- | --- |
|  |  | CAI | CAII | CAIII | CAIV | CAVA | CAVB | CAVI | CAVII | CAIX | CAXII | CAXIII | CAXIV |
|  | **Non-covalent** |  |  |  |  |  |  |  |  |  |  |  |  |
| **1** | VD10-35 | 0.2 | 20 | 20000 | 500 | 300 | 20 | 70 | 7 | 40 | 300 | 30 | 30 |
| **2** | JA16-9-26 | 0.7 | 50 | 60000 | 2000 | 600 | 5 | 400 | 20 | 5 | 500 | 100 | 20 |
| **3** | AV19-64-1 | 0.6 | 20 | 20000 | 700 | 200 | 2 | 200 | 1 | 50 | 600 | 2 | 8 |
| 4 | AV21-05 | 1 | 10 | ≥200000 | 700 | 70 | 8 | 300 | 3 | 20 | 200 | 10 | 10 |
| 5 | AV19-36 | 0.2 | 10 | 50000 | 700 | 200 | 20 | 700 | 5 | 20 | 200 | 4 | 4 |
| 6 | AV20-08 | 60 | 70 | 2000 | 20 | 10 | 3000 | 1000 | 40 | 20 | 700 | 800 | 40 |
| 7 | AV19-56 | 0.3 | 20 | 20000 | 400 | 300 | 20 | 200 | 20 | 20 | 100 | 20 | 9 |
| 8 | AV17-06 | 0.2 | 10 | 40000 | 800 | 300 | 8 | 300 | 3 | 10 | 200 | 2 | 5 |
| 9 | AV23-30-1 | 0.03 | 2 | 7000 | 100 | nd | nd | 1000 | nd | 3 | 40 | 3 | 5 |
| 10 | AV17-30 | 4000 | 100 | ≥200000 | 100 | 4000 | 20 | 400 | 30 | 0.1 | 10 | 20 | 40 |
| 11 | VD11-4-2 | 800 | 60 | 30000 | 60 | 3000 | 20 | 70 | 9 | 0.03 | 3 | 4 | 4 |
| **Covalent** |  |  |  |  |  |  |  |  |  |  |  |  |  |
| 12 | VD10-35-2 | 0.003 | 0.2 | 30000 | 10 | 2 | 100 | 0.1 | 0.007 | 0.03 | 0.03 | 0.01 | 0.007 |
| 13 | VD18-02 | 50000 | 300000 | ≥1000000 | ≥1000000 | ≥1000000 | ≥1000000 | ≥1000000 | ≥1000000 | 300000 | 300000 | ≥1000000 | 30000 |
| 14 | AV21-03 | 0.003 | 0.07 | 40000 | 2 | 3 | 0.01 | 0.05 | 0.002 | 0.02 | 0.02 | 0.003 | 0.1 |
| 15 | AV22-48 | 0.007 | 0.3 | 30000 | 10 | 1 | 0.01 | 0.3 | 0.006 | 0.07 | 0.03 | 0.06 | 0.06 |
| 16 | AV22-162 | 0.008 | 0.4 | 20000 | 60 | nd | nd | 0.3 | 0.8 | 0.09 | 0.3 | 0.05 | 1 |
| 17 | AV21-25 | 0.002 | 0.1 | 20000 | 50 | 3 | 0.02 | 0.4 | 0.8 | 0.04 | 0.07 | 0.01 | 0.3 |
| 18 | AV22-138 | 0.008 | 0.4 | 4000 | 20 | 1 | 0.005 | 0.1 | 0.4 | 0.04 | 0.1 | 0.02 | 0.4 |
| 19 | AV23-04 | 0.003 | 0.3 | nd | 20 | nd | nd | 0.06 | 0.4 | 0.04 | 0.06 | 0.02 | 0.5 |
| 20 | AV19-37 | 0.05* | 0.8 | ≥200000 | 1 | 500 | 4 | 10 | 0.04 | 0.004 | 0.02 | 0.02 | 0.2 |
| 21 | AV21-08 | 0.005* | 0.7 | ≥200000 | 2 | 1000 | 0.02 | 3 | 0.2 | 0.002 | 0.01 | 0.005 | 0.3 |
| 22 | AV22-149 | 0.01* | 1 | 100000 | 2 | nd | 0.1 | 40 | 80 | 0.008 | 0.01 | 0.02 | 2 |
| 23 | AV21-47 | 10*<br>or<br>0.02 | 1 | ≥200000 | 0.2 | 5000 | 0.3 | 30 | nd | 0.003 | 0.1 | 0.02 | 0.5 |
| 24 | AV22-132 | 20* or<br>0.02 | 2 | ≥200000 | 4 | 3000 | 0.2 | 2000 | nd | 0.009 | 0.2 | 0.01 | 0.9 |

\* 2 different fluorescence shifts were observed of which the dominating one was used for $K_{d,app}$ determination.

### Design of dual CAIX and CAXII-recognizing covalent compounds

A shown above, compounds with a carbamate leaving group are better than ester groups. The carbamate compounds seemed to have a good balance to enhance interaction with CAIX via a covalent bond and, at the same time, should have sufficiently low reactivity to react with any unintended proteins. It has been demonstrated that in some cancers, CAXII isozyme is overexpressed instead of CAIX and sometimes both of these isozymes are expressed. Therefore, dual attack on CAIX and CAXII could be beneficial over a single CAIX interaction. At the same time, inhibition of remaining 10 CA isozymes is expected to cause more harm than benefit.

As seen on the arrows going from two left non-covalent compounds to the adjacent covalent compounds (Figure 8), there is a significant gain in affinity due to the covalent bond, at least several hundred-fold stronger binding. Second, the presence of the cyclooctyl or cyclododecyl ring at the meta-position relative to sulfonamide increased the affinity for CAIX, but – what i even more significant – greatly reduced compound affinity for non-target CAI and CAII. The covalent CAIX-targeting compounds reached the affinity of single-digit picomolar, an incredibly high value, never reached by any CAIX-binding compounds and rare among any interactions. Despite apparent selectivity of compound **23** to CAIX compared to CAI, compound **22** is more promising as drug candidate due to its slower covalent bond formation rate and lower associated off-target toxic effects.

**Figure 8.**
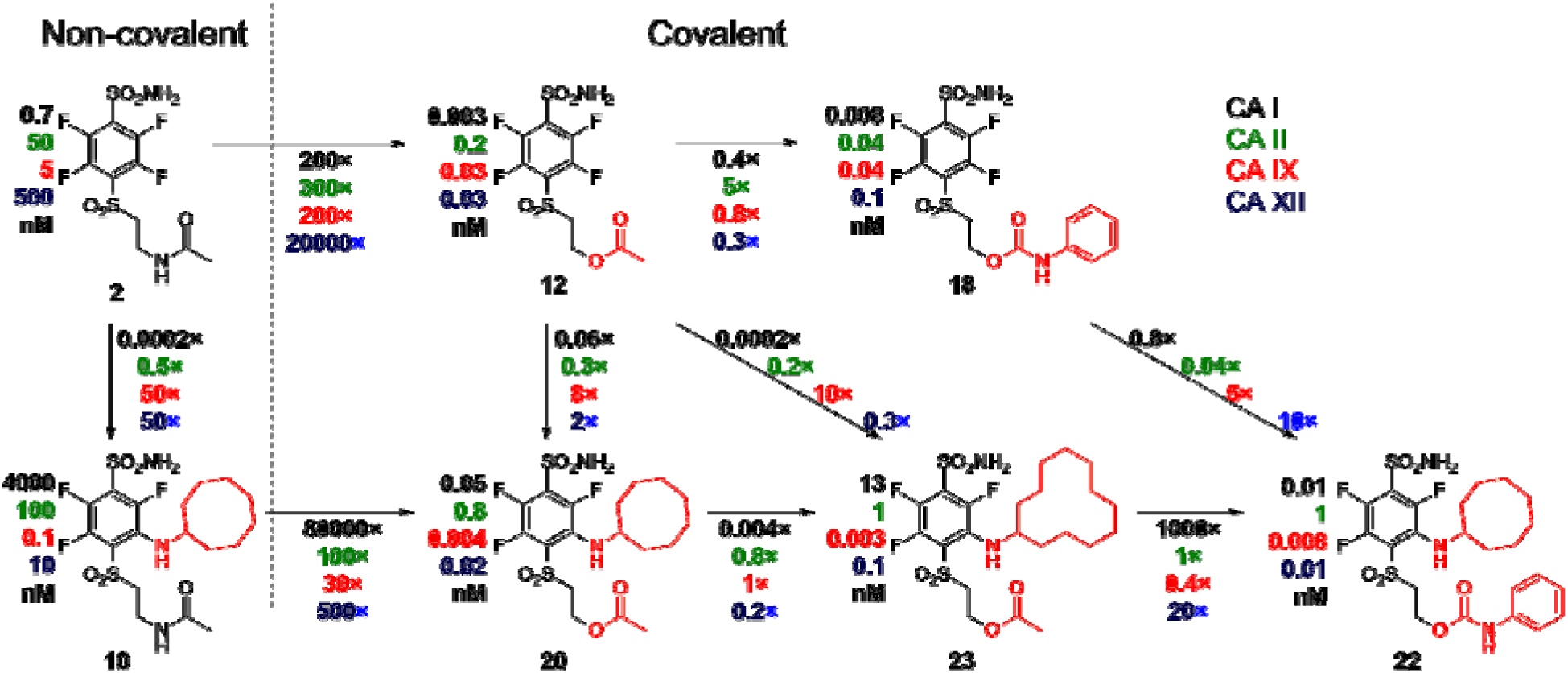
A correlation map between chemical structures and binding affinities showing apparent dissociation constants of compounds for CAI, CAII, CAIX, and CAXII, in nM units. Apparent affinities are listed next to compound structures and the ratios of K_d,app_ – above or below the arrows connecting compounds that are compared. The two left compounds, both upper and lower, located to the left of the vertical dashed line do not form covalent bonds with the proteins, while the rest of the compounds form the covalent bond.

## DISCUSSION

In this work we are introducing a novel previnylsulfone warhead for targeted covalent modification of proteins. The designed group of compounds bound to CAIX, an anticancer target protein, via triple binding model: 1) sulfonamide group formed a coordination bond with the Zn(II) in the active site, 2) hydrophobic ring selectively recognized CAIX over other CA isozymes, and 3) covalent bond formed between the compound and histidine residue of the protein. To reduce reactivity, compounds with the carbamate leaving group were designed. A large series of synthesized compounds distinguish the chemical structures necessary for covalent modification from non-covalent reversible interaction with the protein.

In recent years, covalent inhibitors gained much attention due to their advantages such as complete target protein inhibition, low dosage and effectiveness against mutated targets (unless mutation happens in the targeted nucleophilic residue) (46–48). Although there is still a lot of concern for covalent compound off-target toxic effects, success stories like ibrutinib or, almost a century ago discovered penicillin and aspirin prove that carefully designed covalent compounds can be safely used (49, 50). To this day, the most used strategy in the design of covalent inhibitors is by attaching an optimized “warhead” to a known lead compound. Such inhibitors exhibit efficient and full target inactivation. There are quite a few known electrophilic groups acting as efficient covalently-modifying warheads. However, it is necessary to choose electrophiles with balanced reactivity to avoid off-target toxicity while maintaining steady covalent bond formation with targeted amino acid residue. Thus, among many discovered warheads only six are FDA approved (3). To the best of our knowledge, the fragment SO_2_CH_2_CH_2_OCOR has never been previously described in the literature as a precursor of warhead, a pro-drug capable to rearrange to vinyl-sulfonyl moiety and form covalent bond with the target enzyme.

Nevertheless, it is known that compounds bearing SO_2_CH_2_CH_2_OCOR fragment can rearrange to vinyl-sulfonyl moiety which has been known as a covalently-modifying warhead by reacting with lysine/cysteine residues (3). In most cases it was demonstrated by chemical reaction in basic environments (39–41). However, here we demonstrate that the vinyl-sulfonyl group can react with the His residue. Formation of the covalent bond was shown by X-ray crystallography and 2D NMR. Inability to regain enzymatic activity by dialysis of the compound also confirmed the covalent bond formation.

Several methods exist for comparing the affinity of covalent inhibitors for a target protein, such as comparing compound *K*_d_, IC_50_ or *K*_i_ (47, 51). However, due to time-dependent nature of covalent inhibitor action, the conventional comparisons become challenging or even impractical using these parameters (47, 51). In the case of IC_50_, this value represents the compound concentration inhibiting half of the target enzyme molecules. However, in the case of covalent inhibititors that react irreversibly but slowly, given enough time, covalent inhibitors should give IC_50_ values equal to half of the target concentration as a result of disrupted binding equilibrium (52). The same principle applies to *K*_i_, where if covalent bond formation outpaces compound dissociation, leading to near zero *k*_off_, the observed *K*_i_ values should also approach zero over time. Thus, the *K*_i_ alone is insufficient, since it does not take into account the second stage involving covalent bond formation. In a covalent inhibition model, where initial non-covalent binding precedes covalent bond formation, the most accepted way of describing covalent inhibitor binding commonly involves using k_inact_/*K*_i_ or % covalent occupancy derived from covalent kinetics and pharmacokinetics (52). This approach, however, has limitations especially with compounds exhibiting extremely high picomolar apparent affinities. We propose and have demonstrated that the thermal shift assay could be employed for precise determination of such affinities.

Considering the already complicated covalent inhibitor evaluation due to two-step mechanism, it is even harder to assess compounds bearing SO_2_CH_2_CH_2_OCOR fragments because they act as prodrugs. We must consider an additional step – the elimination reaction, during which an active compound is formed, capable of binding to the protein covalently. Without the elimination step, the active compound is not formed, and the formation of a covalent bond with CA is impossible. If the elimination reaction rate is higher than the covalent bond formation rate, we can ignore it and consider it as part of *k*_inact_ because the limiting step will be *k*_inact_. However, it is challenging to determine separate and correct K_i_, elimination rate constant and *k*_inact_ values for CA isozymes because of exceptionally high compound affinity. Nevertheless, using fluorescent thermal shift assay (FTSA), we could determine the *K*_d,app_ for all twelve catalytically active CA isozymes and could perform an affinity correlation between different CA isozymes. Prior applications of the thermal shift were limited to test the change in the melting temperature of the protein upon covalent modification (53) (54) (55) (56) (57). Therefore, we conclude, to the best of our knowledge, that the apparent affinity determination by FTSA of covalent compound binding to proteins is being demonstrated here for the first time.

There are only few examples of covalent inhibitors of CA – bromoacetazolamide and N-bromoacetylacetazolamide and several compounds designed for enzyme tagging which have been tested on CAII as a model enzyme (33, 58, 59). In bromoacetazolamide case, even with 20 molar surplus, bovine CAII was not fully modified even after 24 hours (58). The experiments with bromoacetazolamide were performed in basic environment (pH 8.2 and 8.7), which was favorable condition for covalent bond formation and thus it is hard to compare with other compounds (60). The covalent tag bearing vinyl-sulfonyl warhead showed better results compared to bromoacetazolamide (>90% of compound covalently bound to bovine CAII after 10 hours at pH 7.4). The compounds **20** and **12** irreversibly bound and inhibited most of the human CA isoforms in less than 2 hours (at pH 7.0) and possibly are the first covalent CA inhibitors tested against CA isozymes, except the above described inhibitors of CAII and CAI, with vinyl-sulfonyl reacting with the histidine residue (33).

To assess the potential of covalent compounds for drug development, it is crucial to investigate whether they can indiscriminately react with different histidine and nucleophilic groups in proteins. Our testing involved examining the binding of these compounds to live cancer cells, thereby interacting with all proteins exposed on the cancer cell surface. The results demonstrated that these compounds exhibited specific and exclusive binding to the target protein CAIX, expressed in hypoxia-grown cancer cells. Notably, these compounds displayed the highest affinities among numerous CAIX inhibitors described in the literature, and their covalent binding led to irreversible inactivation of CAIX expressed in live cancer cells. This compelling evidence suggests significant potential for the development of these compounds as anticancer drugs.

## MATERIALS AND METHODS

### Enzyme purification

All recombinant human CA isozymes were produced as described previously by using either bacterial or mammalian expression system (61, 62). For isozymes possessing transmembrane parts, only catalytic domains were produced. The production of the catalytic domain of CAIX in methylotrophic yeast *Pichia pastoris* was performed as described in (63). Protein purity was confirmed by SDS-PAGE, and MW was confirmed by mass spectrometry.

### Fluorescent thermal shift assay (FTSA)

Thermal unfolding experiments of the purified CA isozymes were carried out by a real-time PCR instrument, Rotor-Gene Q, containing six channels. Twofold serial dilutions of the 10 mM compound stock in DMSO were made by adding 10 μL of DMSO to 10 μL of each compound solution. Overall, 8 different compound concentration solutions were prepared, including the 10 mM compound stock concentration and a sample containing only DMSO, without a ligand. To prepare 12 different concentrations of the ligand, 1.5-fold serial dilutions of 10 mM compound stock were performed by adding 10 μL of DMSO to 20 μL of each compound solution (the last sample contained no ligand). Each prepared compound solution was diluted 12.5 times with the assay buffer (50 mM sodium phosphate (pH 7.0), 100 mM NaCl. The 10 μM CA isozyme solution (or 20 μM of CAIV) was prepared in the same assay buffer, which contained a reporter dye (200 μM ANS or 200x diluted Glomelt). 5 μL of prepared CA solution with dye was added to the 100 μl of PCR tubes. Subsequently, 5 μL of the compound solution is added. The tubes are placed into the real-time thermocycler, and protein unfolding is measured by increasing the temperature from 25 °C to 90 °C, at the rate of 1 °C per minute and measuring the fluorescence of the dye. The raw data were analyzed to determine the *T*_m_ of the proteins. *T*_m_ values were plotted as a function of ligand concentration and the model was fitted to the dosing curves to obtain the binding affinities using Thermott software (64).

### CAIX activity measurement by the stopped-flow assay

CAIX activity was measured in the absence and presence of the compound before and after dialysis. 1.5 µM CAIX was incubated with 15 μM **20**, 15 μM **22** or 100 µM **5** for two hours. The solution of CAIX without a compound and the CAIX-compound complexes were then dialyzed in 25 mM Tris buffer solution (pH = 7) containing 50 mM NaCl (by changing buffer four times every 8/16 hours). CO_2_ hydration velocities were measured by recording the absorbance of phenol red (final concentration 50 µM) at 557 nm using an Applied Photophysics SX.18MV-R stopped-flow spectrometer. Experiments were performed at 25 °C using 25 mM Hepes containing 50 mM NaCl, pH 7.5.

### Mass spectrometry

Mass spectrometry experiments were performed with an electrospray ionization time-of-flight mass spectrometer (Q-TOF). The 0.1 mg/mL CA isozyme solution was prepared in the absence or presence of compounds (1:2; 1:5 or 1:10 CA isozyme : compound molar ratio). The solution was incubated for one hour at room temperature before analysis. The final DMSO concentration was 1% (v/v).

### 2D NMR

All NMR spectra were recorded using a 600 MHz Bruker Avance Neo spectrometer equipped with a cryogenic probe. 2D ^15^N–^1^H HSQC spectra of ^15^N labeled CA2 solution (270 µM 15N CA2, 20 mM sodium phosphate buffer, 50 mM NaCl, 5% D2O, pH 6.8) were recorded at 25 °C using 256 increments in the indirect dimension and 8 scans. The spectra were recorded when the protein solution contained different concentrations of covalent ligand: 0.27 mM, 0.53 mM, 1.0 mM and 1.5 mM (the final DMSO concentration was 7.5%) or non-covalent ligand: 0.35 mM, 0.70 mM, 1.0 mM, 2.0 mM (the final DMSO concentration was 3%). The spectra were analyzed using Topspin 4.1.3 and CcpNMR 2.4.1 analysis software (65).

### HPLC analysis

The HPLC separation was done as previously described (66). In brief, the samples were separated and analysed using Shimadzu UFLC system with a CMB-20A communication module, two LC20AD quaternary and isocratic pumps, a SIL-20AC autosampler, a CTO-20A column compartment and an SPD-M20A DAD detector (Shimadzu Corp., Japan). For the detection of the eluting molecules, the DAD spectra recording was set from 190 to 750 nm with a data rate of 6.25 Hz. The ACE C18-PFP HPLC separation column (100 x 4.6, 3 µm, Avantor) was used as a stationary phase. The HPLC grade MeCN (Fisher Scientific) and Milli-Q water (18.2 MΩ cm^−1^, Milli-Q Plus system, Millipore Bedford, MA, USA) were used for the RP-HPLC separation.

The samples were separated using a trinary gradient consisting of ultrapure water (eluent A), MeCN (eluent B) and 1% TFA in ultrapure water (eluent C). A constant 10% flow of eluent C was used to maintain 0.1% TFA concentration in the column throughout the separation experiment. The gradient between eluents A and B was 36% (0 min), 63% (20 min), and 63% (21 min). Before each analytical run, the column equilibration (10 column volume) was performed. The column thermostat was set to 40 °C and the flow rate to 1 mL min^-1^.

### Protein-compound complex sample preparation for Data-dependent analysis (DDA)

The 0.1 mg/ml CAXIII solution was prepared with compound **12** (molar ratio 1:2 CAXIII : **12**) and filter aided sample preparation (FASP) (67) was used for protein digestion prior to mass spectrometry analyses.

### Data-dependent analysis

Data-dependent analysis (DDA) was performed with the nanoAcquity coupled to a Synapt G2 HDMS mass spectrometer (Waters). For DDA, the instrument performed a 0.7 s MS scan (350-1350 scan range) followed by MS/MS acquisition on the top 5 ions with charge states 2+, 3+ and 4+. MS/MS scan range was 50-2000 Da, 0.6 s scan duration with exclusion after 2 MS/MS scans were acquired, and dynamic exclusion of ions within 100 mDa of the selected precursor m/z was set to 100 s.

The Progenesis QI for proteomics software (Nonlinear Dynamics), in combination with the Mascot server (2.2.07) was employed to identify peptides. The acquired raw files were imported into the Progenesis QI for proteomics software and MS2 spectra were exported directly from Progenesis in mgf format and searched using the MASCOT algorithm.

### Crystallization and structure solution

Human CAI in buffer containing 20 mM HEPES pH 7.6 and 50 mM NaCl was mixed with the **21** in DMSO in the ratio 1:1,1 and incubated 2.5 hrs at room temperature. The protein-**21** complex was concentrated to 30 mg/ml, and crystallization in sitting drops was started. Crystallization solution contained 0.1 M TrisHCl pH8.5, 0.2 M NaCl and 28% (w/v) PEG3350. Before cryo-cooling crystal was shortly incubated in cryo-protection buffer containing 0.1 M TrisHCl pH8.5, 15 % (w/v) PEG8000 and 20% (v/v) Ethylene glycol. The synchrotron data was collected at beamline P13 operated by EMBL Hamburg at the PETRA III storage ring (DESY, Hamburg, Germany).

Diffraction data were integrated by XDS (68), scaled using AIMLESS 0.7.4 and other CCP4 tools v. 7.1.002 (69). The structure was solved by molecular replacement using MOLREP v.11.7.02 (70) and 2CAB as an initial model. The model was refined by REFMAC v. 5.8.0258 (71) and rebuilt in COOT v.0.9 (72). Inhibitor model was created and minimized using AVOGADRO v. 1.2.0 (73).

CAII protein was concentrated to 10 mg/ml in a 20 mM Tris-HCl buffer. It was then mixed with compound **20** (5 mM final concentration) and incubated overnight at 4°C. Previously known crystallization conditions for CAII did not yield any crystals when co-crystallizing with compound **20**. Crystallization condition screening was performed using the Morpheus screen from Molecular Dimensions. After slight optimization, crystals grown using sitting drop technique in 0.06 M magnesium chloride hexahydrate; 0.06 M calcium chloride dihydrate; 0.1 M Tris pH 8.5; 20% v/v PEG 500 MME; 10% w/v PEG 20000 conditions diffracted at 1.4 Å resolution. The alteration in crystallization conditions was likely due to the formation of a covalent bond between the ligand and the enzyme. The dataset of the CAII-ligand complex was collected at BESSY II beamline 14.1 and processed using MOSFLM(74) and SCALA (75). Molecular replacement was performed using MOLREP (70) with 5AMD (76) as the initial model.

CAIX protein was concentrated to 10 mg/ml in a 20 mM Tris-HCl buffer. It was then mixed with compound **23** (5 mM final concentration) and incubated overnight at 4°C. Crystals grew using similar co-crystallization conditions as described before (63).

The dataset of the CAIX-ligand complex was collected at the Diamond Light Source beamline I03 and processed using XDS (68) and AIMLESS (69). Molecular replacement was performed using MOLREP (77) with 8Q18 (78) as the initial model.

Model refinement was performed with REFMAC (71), and the structures were visualized using COOT (72). Ligand parameter files were generated using LIBCHECK (79), and the ligand was manually fitted to the electron density map in COOT (72). The coordinates and structure factors have been deposited in the PDB. The PDB access code, along with the data collection and refinement statistics, are provided in Table ST1.

### Determination of compound K_d_ values for cellular CAIX

Human cervical adenocarcinoma cells (HeLa) were cultured in Dulbecco’s Modified Eagle’s Medium (DMEM) with GlutaMAX™ (Gibco, ThermoFisher) supplemented with 10% fetal bovine serum (ThermoFisher) in a humidified atmosphere at 37 °C and 5% CO_2_.

A covalent compound competition experiment with fluorescein-labeled compound GZ19-32 was conducted as described previously (Matuliene et. al., 2022). In brief, HeLa cells were cultivated in DMEM in 12-well plates under hypoxic conditions (1% O_2_) for 72 hours. The 10 serial two-fold dilutions of covalent compounds were prepared in FluoroBrite DMEM (ThermoFisher) starting with 80 nM (1^st^ tube). No compound was added to the last 12^th^ tube (it contained FluoroBrite only). Subsequently, the same volume of 20 nM GZ19-32 was added to each of the 12 tubes and mixed. Cell culture medium was removed from all 12 wells with HeLa cells and 200 µl of prepared compound mixtures were added, followed by 20 min incubation at 37 ℃ under normoxic conditions (21% O_2_). Post-incubation, the compound solutions were aspirated and the cells were washed 3 times with 400 µl of PBS. Finally, the cells were detached from the well plate surface by TrypLE (ThermoFisher), resuspended by pipetting in 200 µl of FluoroBrite DMEM, and 150 µl of the suspension was transferred to black Thermo Scientific™ Nunc MicroWell 96-Well Optical-Bottom Plates for fluorescence and absorbance measurements.

## ABBREVIATIONS

CA: carbonic anhydrase
CAIX: carbonic anhydrase IX
FTSA: fluorescent (fluorescence-based) thermal shift assay
GZ19-32: fluorescein-labeled compound
PBS: phosphate buffer saline
TCI: targeted covalent inhibitors
TSA: thermal shift assay

## ACKNOWLEDGEMENTS

This research was funded by the grant from Research Council of Lithuania No S-MIP-22-35. Access to the EMBL beamline P13 at PETRA III (DESY) and has been supported by iNEXT-Discovery, project number 871037, funded by the Horizon 2020 program of the European Commission. This research was also s614892v1upported within the framework of the European Union’s Recovery and Resilience Mechanism project No.5.2.1.1.614892v1i.0/2/24/I/CFLA/001 "Consolidation of the Latvian Institute of Organic Synthesis and the Latvian Biomedical Research and Study Centre".

## AUTHOR CONTRIBUTIONS

AV, DB, AZ, VD, and DM designed the study, AV performed organic synthesis, AV and DB performed FTSA and inhibition measurements, JL and KT determined X-ray structures of inhibitor bound to CAII and CAIX, AKv and JM determined inhibitor binding to live cancer cells, AR performed MS experiments of proteins and protein-compound complexes, EM and SG determined X-ray structures of covalent inhibitors bound to CAI, ZT and KJ performed the NMR experiments, AKau and MV performed the MS analysis of peptide digest, MG, AM, VJ, and AKaz produced recombinant proteins, HS and FM conceptualized and analyzed.

