## Supplementary Materials for "Targeted anticancer pre-vinylsulfone covalent inhibitors of carbonic anhydrase IX"

**Supplementary material**

Table of Contents

**Crystallography**	2

**Mass spectrometry data**	3

**2D NMR data**	17

**Fluorescent thermal shift assay**	18

**Chemistry**	25

**References** .................................................................................................................................................................... 47

**Crystallography**

Supplementary Table S1. X-ray crystal structure refinement statistics of CA II-**20**, CA I-**21** and CA IX-**23** complexes.

| Structure | CA II – **20** | CA I – **21** | CA IX – **23** |
| --- | --- | --- | --- |
| Space group | P21 | P 21 21 21 | H3 |
| Cell dimensions | | | |
| *a* (Å) | 42.45 | 62.42 | 152.11 |
| *b* (Å) | 41.57 | 73.27 | 152.11 |
| *c* (Å) | 72.79 | 120.57 | 172.51 |
| *β* (^o^) | 104.4 | 90 | 90 |
| Resolution (Å) | 70.50-1.40 | 120.57-1.39 | 52.35-2.20 |
| Highest resolution shell (Å) | 1.40-1.47 | 1.39-1.41 | 2.20-2.25 |
| No. of reflections (unique) | 47352 | 110929 | 75528 |
| No. of reflections in test set | 2410 | 10962 | 3874 |
| Completeness (%) | 96.9 (98.0^a^) | 98.3 (67.6^a^) | 100.0 (100.0^a^) |
| R _merge_ | 0.14 (0.48^a^) | 0.06 (2.7^a^) | 0.07 (1.16^a^) |
| 〈 *I/σI* 〉 | 6.5 (3.0^a^) | 20.4 (0.5^a^) | 17.2 (1.9^a^) |
| Average multiplicity | 5.0 (5.0^a^) | 13.0 (4.7^a^) | 10.7 (11.0^a^) |
| R-factor | 0.18 (0.32^a^) | 0.21 (0.47^a^) | 0.17 (0.26^a^) |
| R _free_ | 0.22 (0.34^a^) | 0.23 (0.48^a^) | 0.20 (0.32^a^) |
| Average B factor (Å^2^) | 11.0 | 26.6 | 52.4 |
| Average B factor for inhibitor (Å^2^) | 18.7 | 33.1 | 73.5 |
| 〈B〉 from Wilson plot (Å^2^) | 8.9 | 22.1 | 49.5 |
| No. of protein atoms | 2045 | 4032 | 7413 |
| No. of inhibitor atoms | 27 | 54 | 124 |
| No. of solvent molecules | 338 | 539 | 421 |
| RMS deviations from ideal values | | | |
| Bond lengths (Å) | 0.02 | 0.01 | 0.01 |
| Bond angles (^o^) | 1.67 | 1.85 | 1.79 |
| Outliers in Ramachandran plot (%) | 0.39 | 0 | 0.21 |
| PDB code | 8OO8 | 8S4F | 9FLF |

a Values in parenthesis are for the high-resolution bin.

**Mass spectrometry data**

Supplementary Table S2. Carbonic anhydrase isozyme masses in the absence of compound and incubated with covalently-modifying compound **12**. All CA isozymes except CAIII were covalently modified by the compound **12** to variable extent.

| Enzyme | Plasmid Number | Theoretical MW | Obtained mass | Protein mass with compound **12**, most intense peak m/z | Difference |
| --- | --- | --- | --- | --- | --- |
| CA I | pL0067 | 31204.7 | 31074.31 (w/o Met) | 31393.63 | 319.32 |
| CA II | pL0059 | 29246 | 29115.54 (w/o Met) | 29434.81 | 319.27 |
| CA III | pL0066 | 31648.9 | 31518.29 (w/o Met), 31696.34 (w/o Met and glycosylated) | 31518.29 (w/o Met), 31696.34 | - |
| CA IV | pL0307 | 30454.6 | 30320.33 (w/o Met and S-S bridge) | 30639.53 | 319.20 |
| CA VA | pL0245 | 31285.3 | 31154.77 (w/o Met) | 31474.07 | 319.30 |
| CA VB | pL0173 | 34193.6 | 34063.56 (w/o Met) | 34382.29 | 318.73 |
| CA VI | pL0339 | 35367 | 35956.25 (glycosylated)  36208.06(glycosylated)  36412.74(glycosylated)  37643.18(glycosylated)  37934.87(glycosylated) | 36275.15  36528.43  36731.48  37962.27  38254.18 | 318.90  320.37  318.74  319.09  319.31 |
| CA VII | pL0137 | 31821.7 | 31689.57 (w/o Met) | 32008.73  32327.99 | 319.03  638.42 |
| CA IX | * | 28061.7 | 28060.32 | 28379.58 | 319.26 |
| CA XII | pL0119 | 29886.3 | 29754.54 (w/o Met) | 30072.81 | 318.27 |
| CA XIII | pL0058 | 29574.3 | 29574.69 | 29893.97 | 319.28 |
| CA XIV | pL0318 | 32129.7 | 31997.12 (w/o Met)  32175.25(w/o Met and glycosylated) | 32316.37  32494.53 | 319.25  319.28 |

* - CA IX mutant C174S, N346Q, prepared in yeast as described in (1).

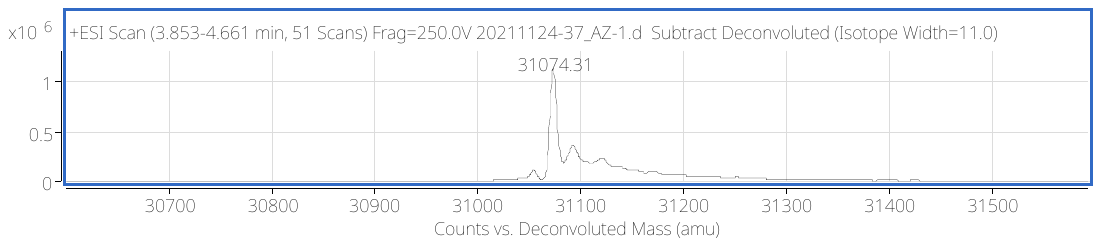

**A**

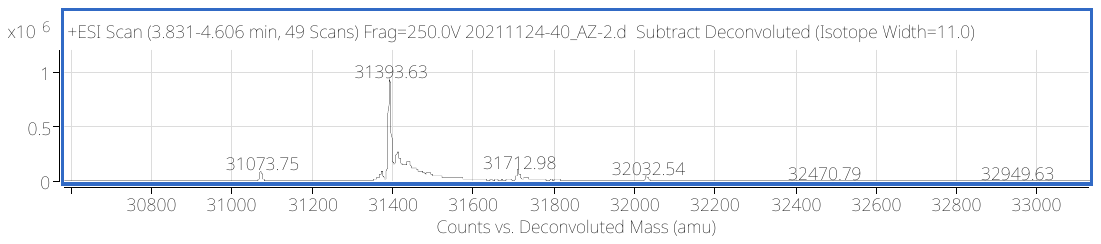

**B**

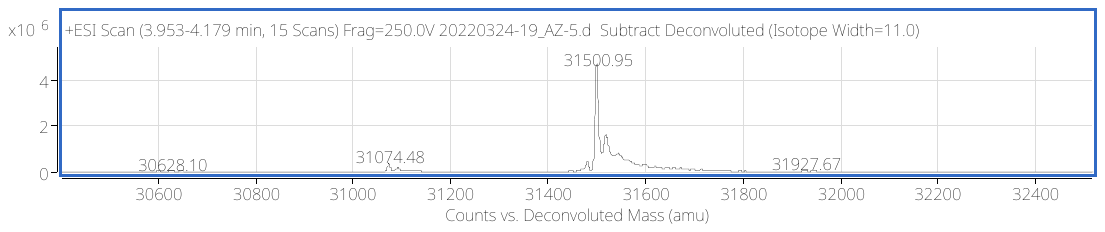

**C**
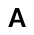

Figure S1. A) CA I mass spectra. B) CA I mass spectra after incubation with **12** C) CA I mass spectra after incubation with **20**.

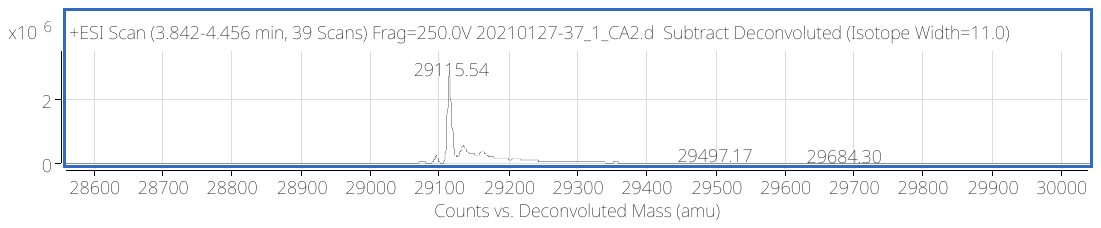

**A**
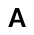

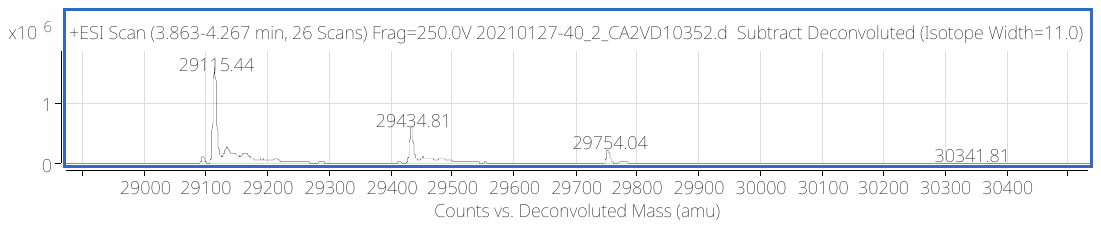

**B**
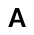

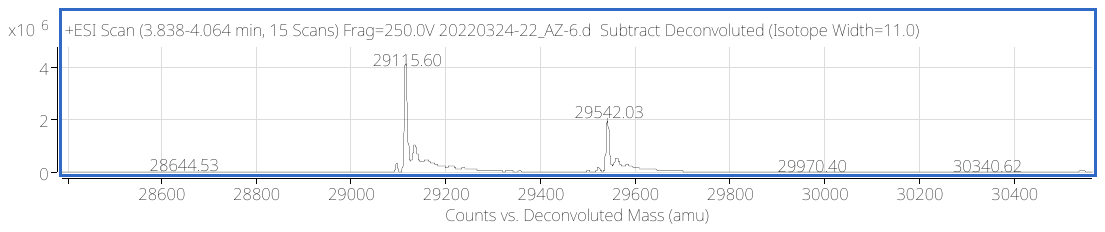

**C**
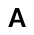

Figure S2. A) CA II mass spectra. B) CA II mass spectra after incubation with **12** C) CA II mass spectra after incubation with **20**.

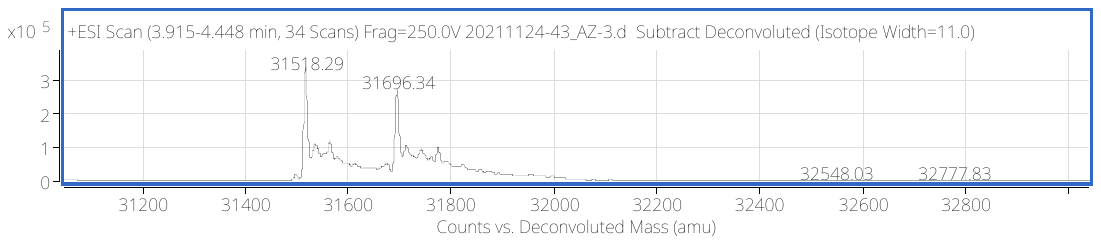

**A**
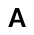

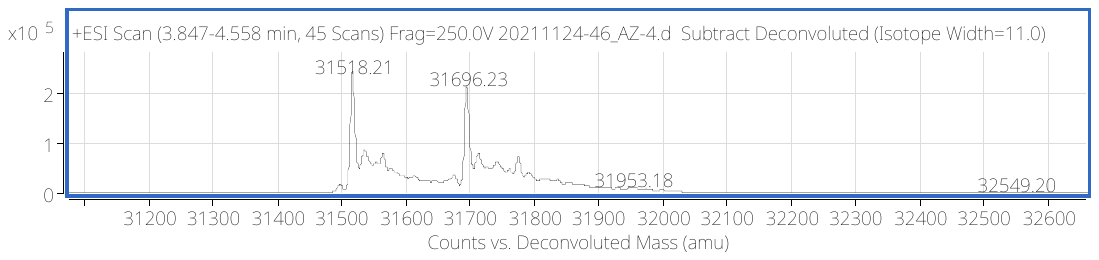

**B**
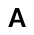

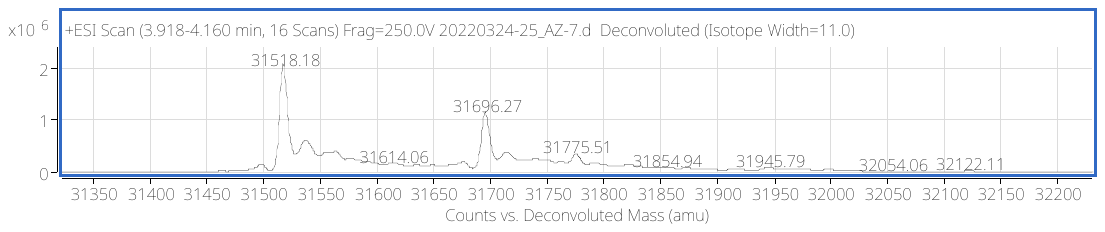

**C**
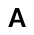

Figure S3. A) CA III mass spectra. B) CA III mass spectra after incubation with **12** C) CA III mass spectra after incubation with **20**.

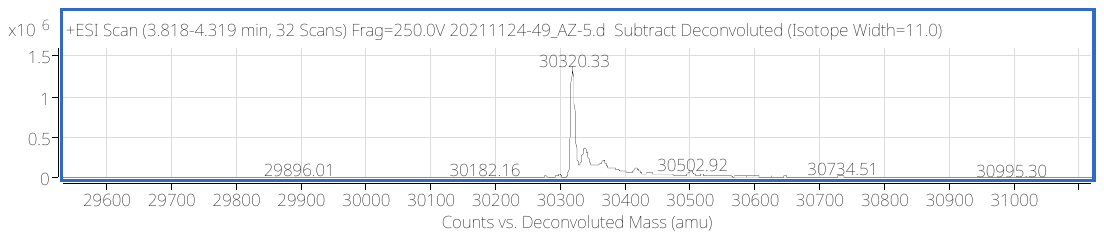

**A**
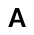

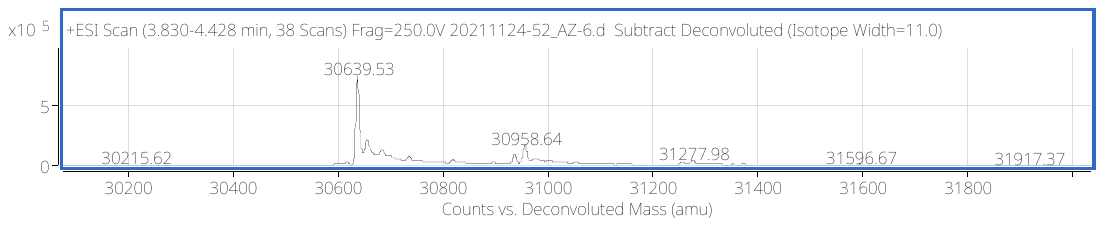

**B**
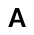

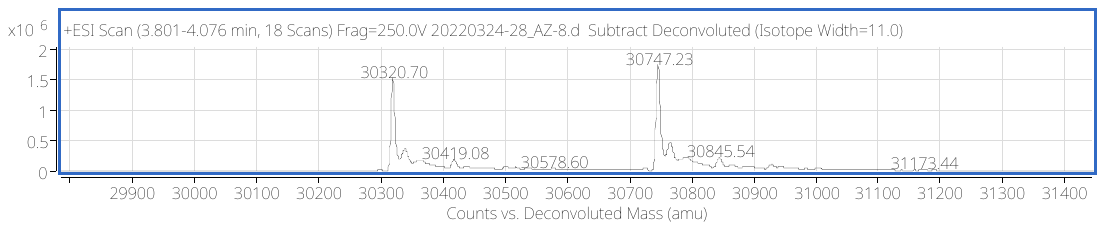

**C**
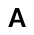

Figure S4. A) CA IV mass spectra. B) CA IV mass spectra after incubation with **12** C) CA IV mass spectra after incubation with **20**.

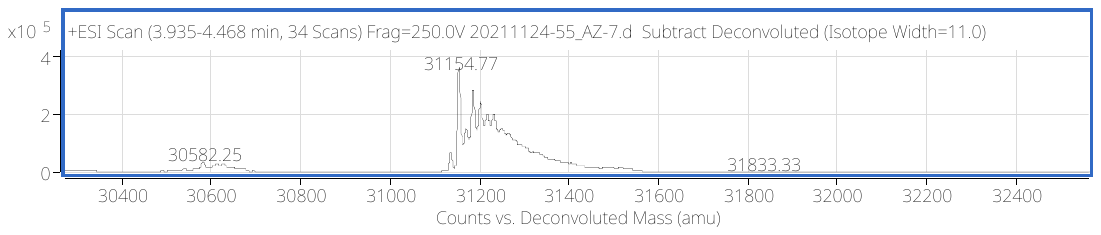

**A**
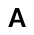

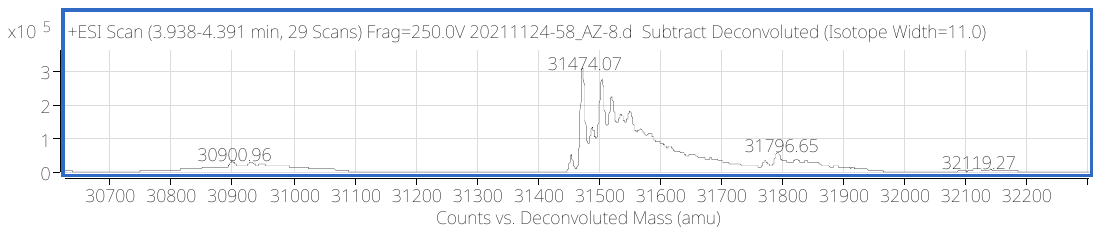

**B**
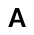

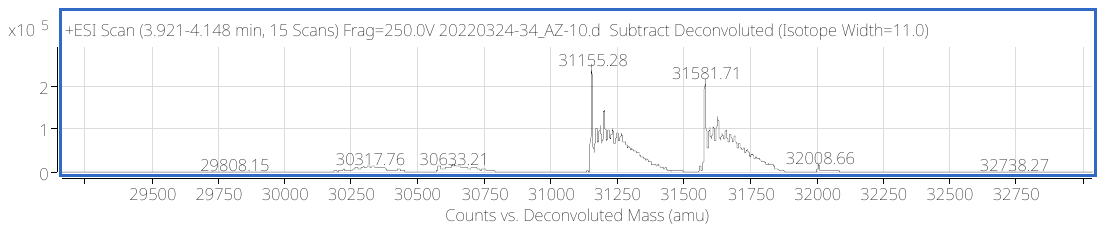

**C**
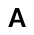

Figure S5. A) CA VA mass spectra. B) CA VA mass spectra after incubation with **12** C) CA VA mass spectra after incubation with **20**.

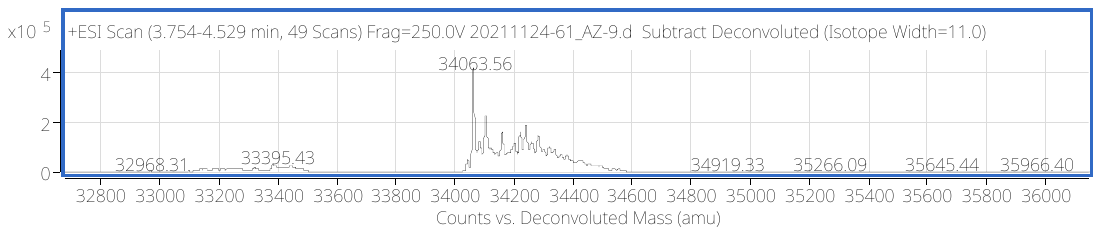

**A**
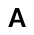

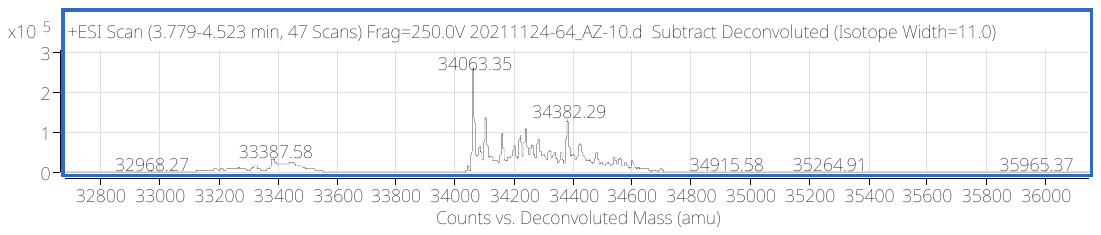

**B**
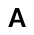

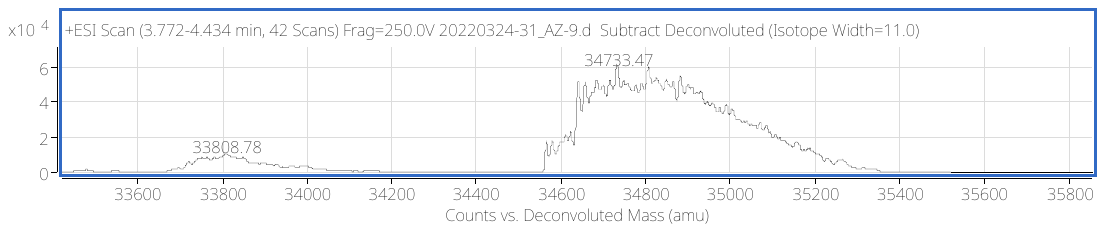

**C**
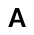

Figure S6. A) CA VB mass spectra. B) CA VB mass spectra after incubation with **12** C) CA VB mass spectra after incubation with **20**.

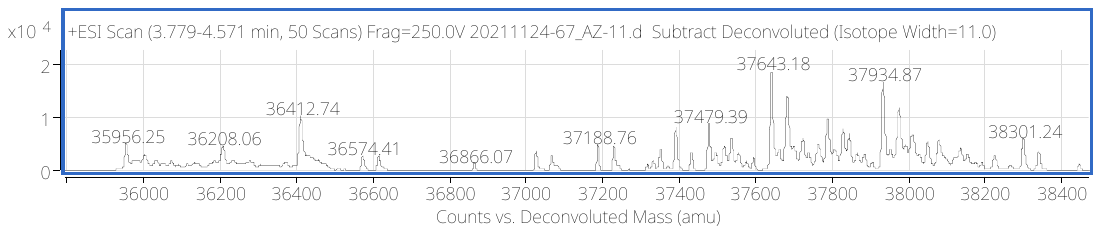

**A**
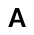

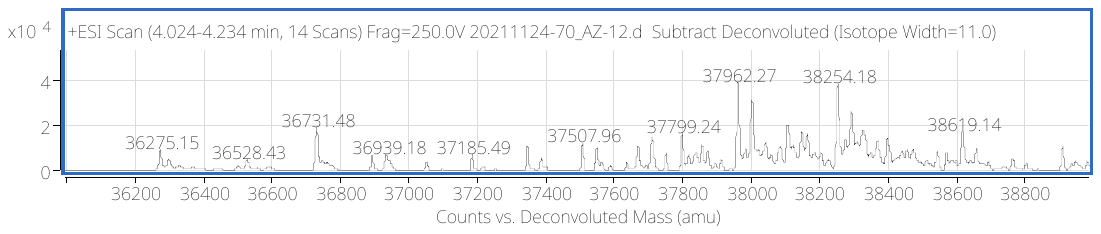

**B**
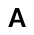

Figure S7. A) CA VI mass spectra. B) CA VI mass spectra after incubation with **12**.

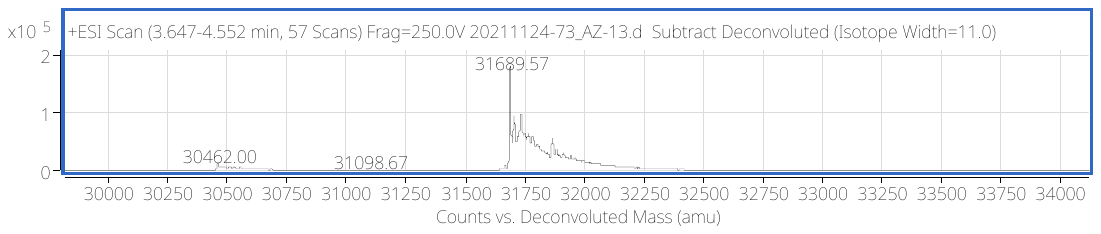

**A**
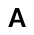

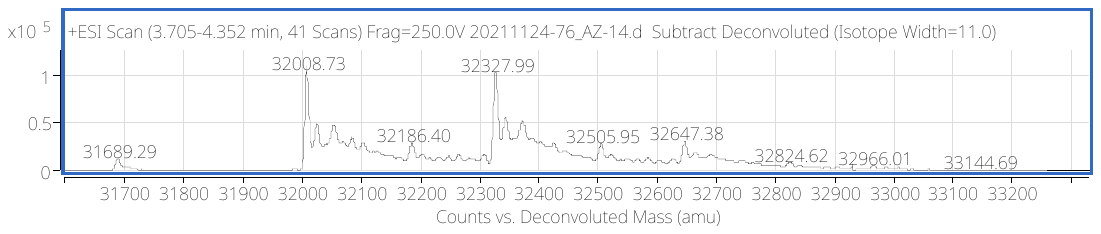

**B**
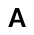

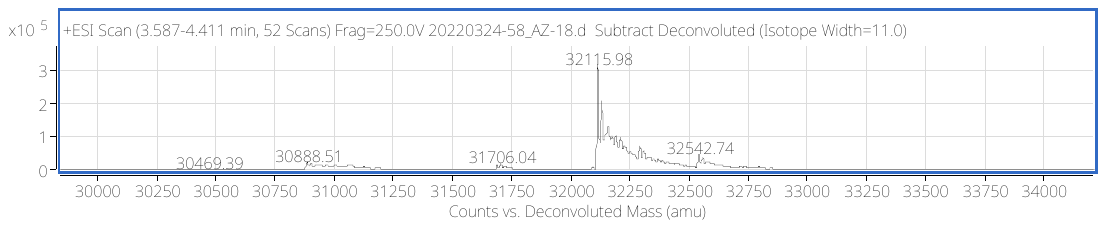

**C**
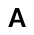

Figure S8. A) CA VII mass spectra. B) CA VII mass spectra after incubation with **12**. C) CA VII mass spectra after incubation with **20**.

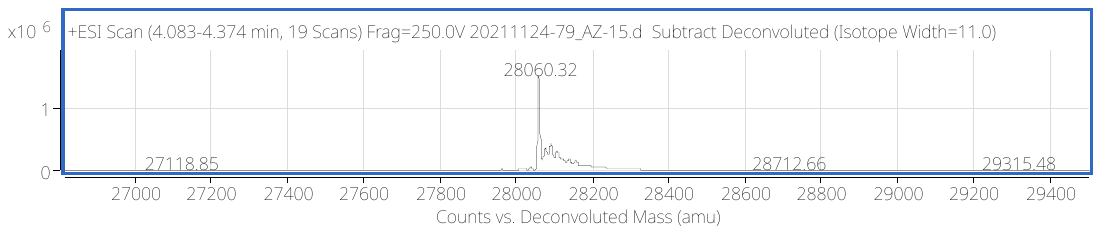

**A**
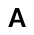

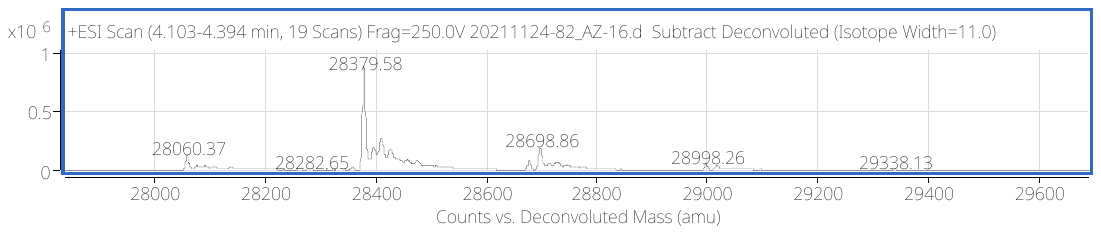

**B**
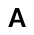

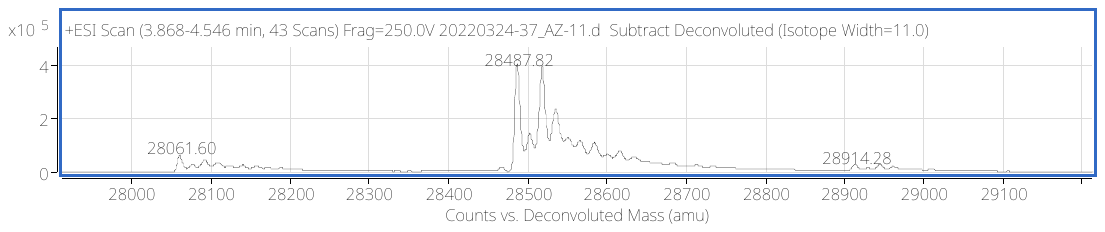

**C**
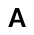

Figure S9. A) CA IX mass spectra. B) CA IX mass spectra after incubation with **12** C) CA IX mass spectra after incubation with **20**.

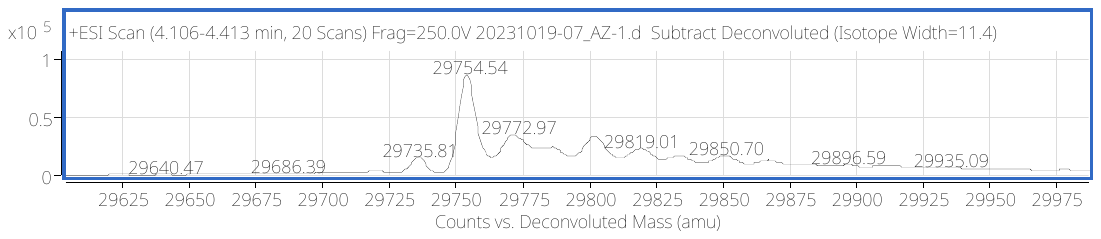

**A**
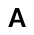

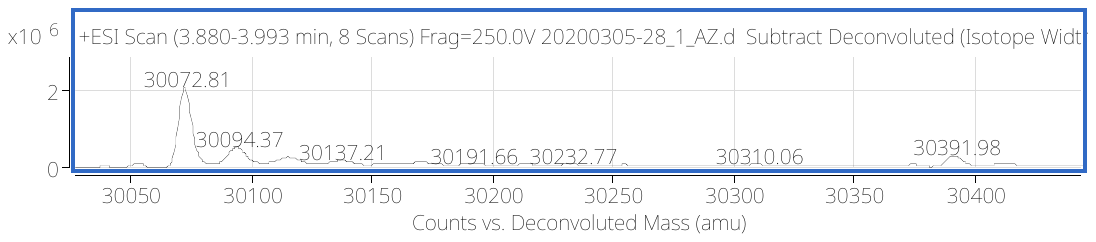

**B**
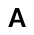

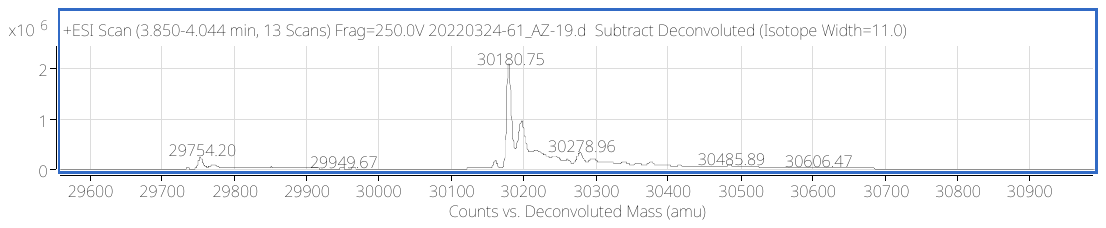

**C**
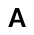

Figure S10. A) CA XII mass spectra. B) CA XII mass spectra after incubation with **12** C) CA XII mass spectra after incubation with **20**.

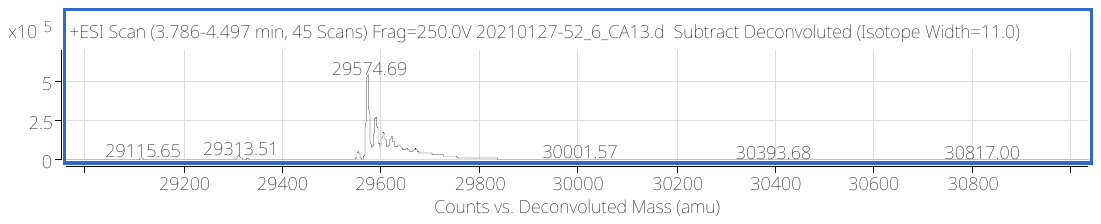

**A**
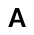

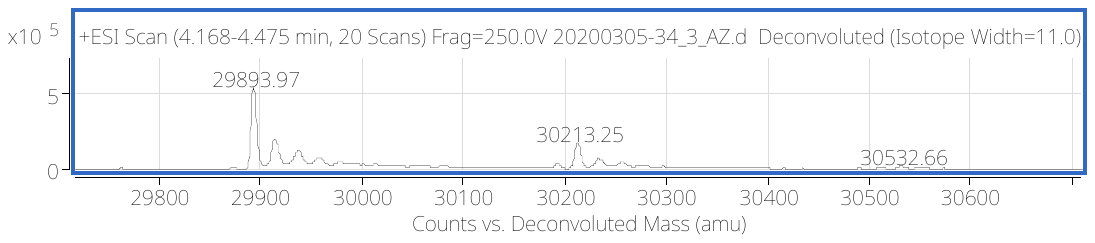

**B**
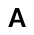

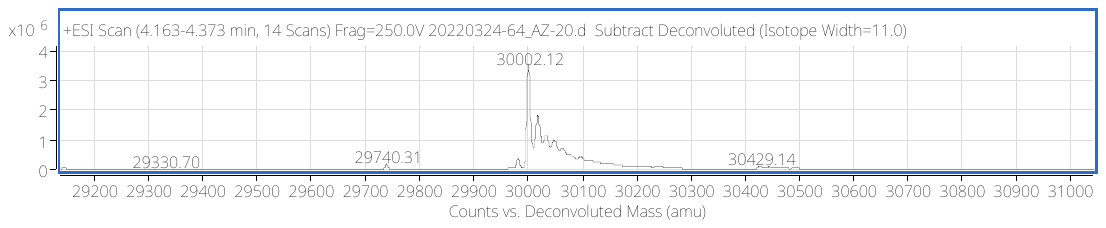

**C**
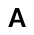

Figure S11. A) CA XIII mass spectra. B) CA XIII mass spectra after incubation with **12** C) CA XIII mass spectra after incubation with **20**.

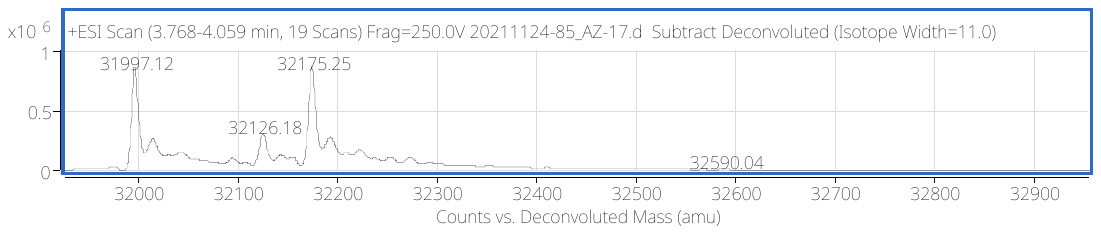

**A**
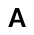

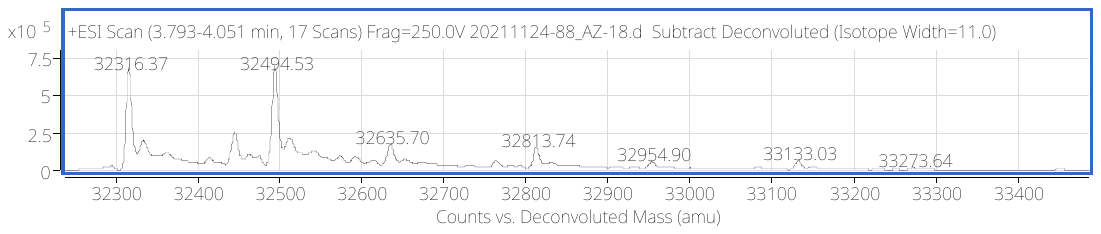

**B**
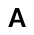

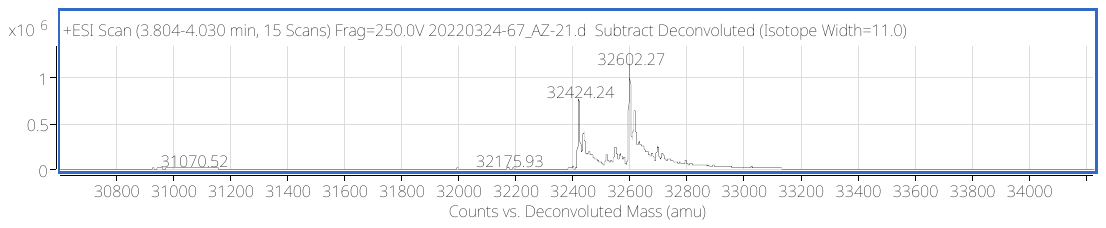

**C**
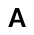

Figure S12. A) CA XIV mass spectra. B) CA XIV mass spectra after incubation with **12** C) CA XIV mass spectra after incubation with **20**.

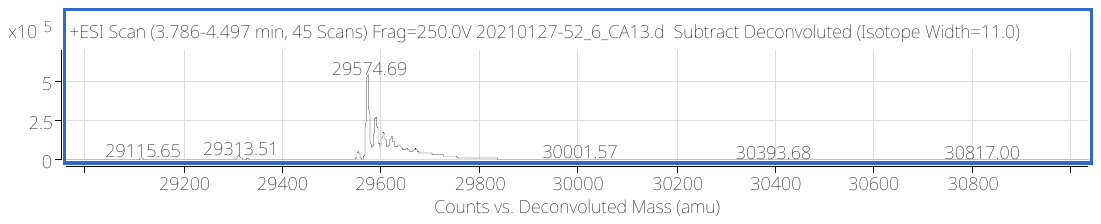

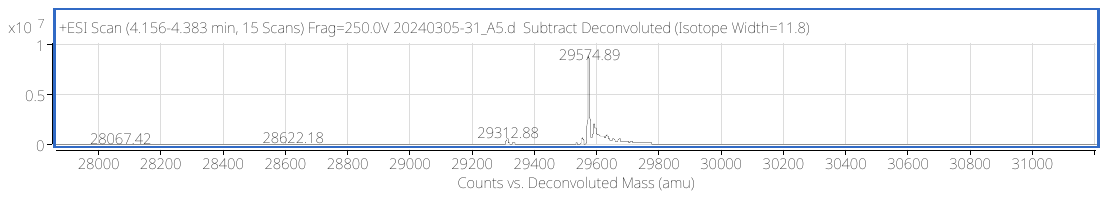

**B**
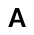

**A**
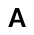

Figure S13. A) CA XIII mass spectra. B) CA XIII mass spectra after incubation with 5 eq. of compound **3**.

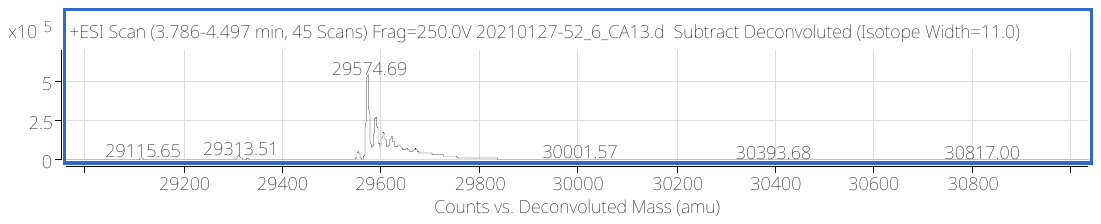

**A**
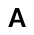

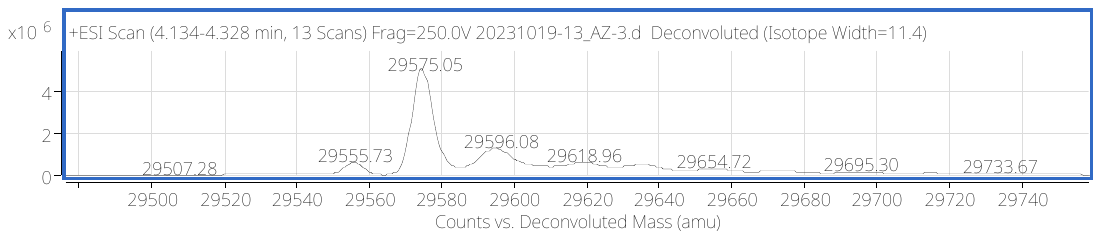

**A**
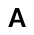

**B**
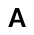

Figure S14. A) CA XIII mass spectra. B) CA XIII mass spectra after incubation with 5 eq. of compound **9**.

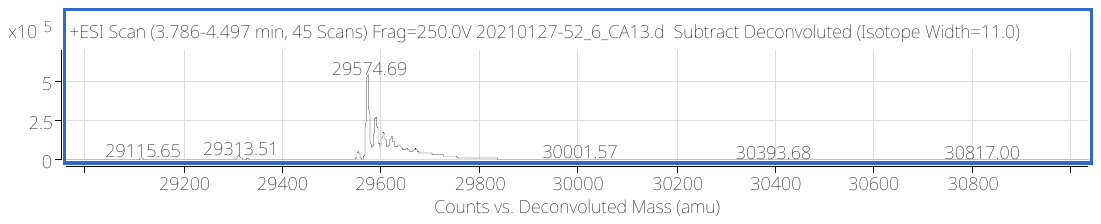

**A**
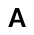

**A**
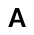

**A**
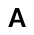

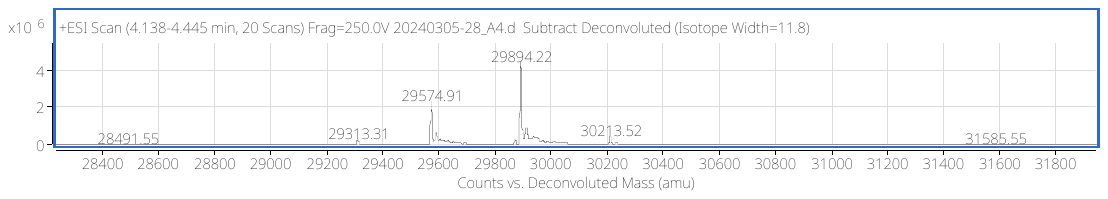

**B**B
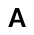

Figure S15. A) CA XIII mass spectra. B) CA XIII mass spectra after incubation with 2 eq. of compound **15**.

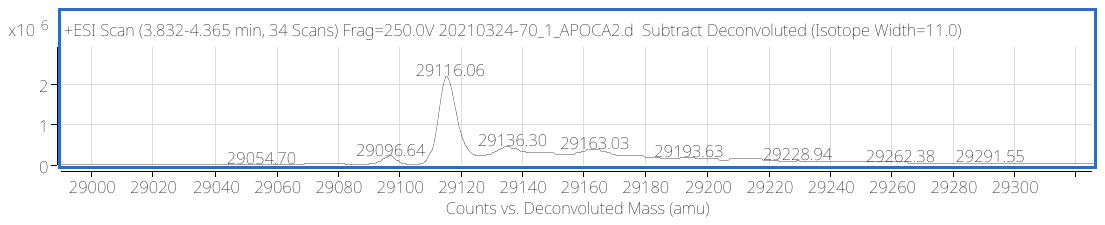

**A**
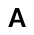

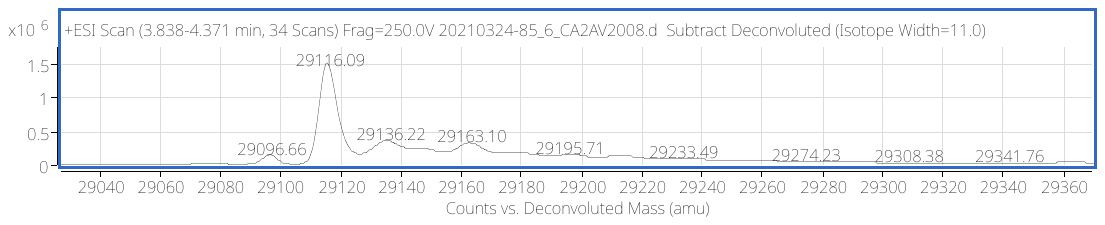

**B**
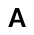

Figure S16. A) CA II mass spectra. B) CA II mass spectra after incubation with 10 eq. of compound **6**.

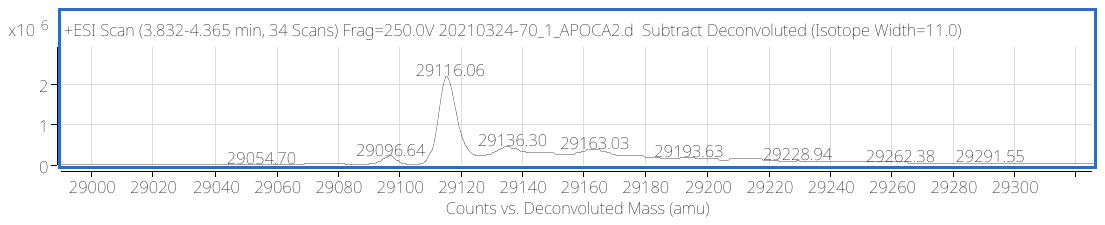

**A**
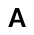

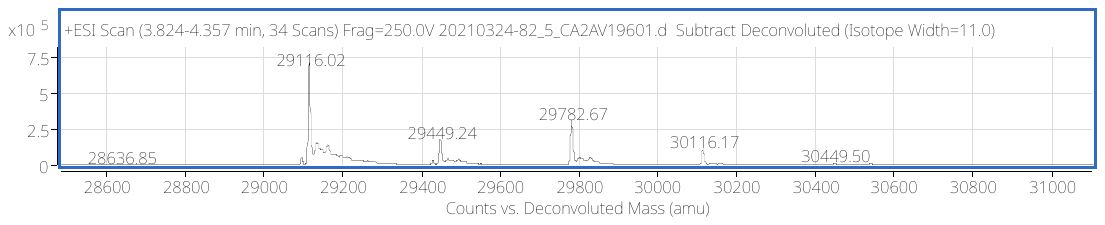

**B**
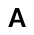

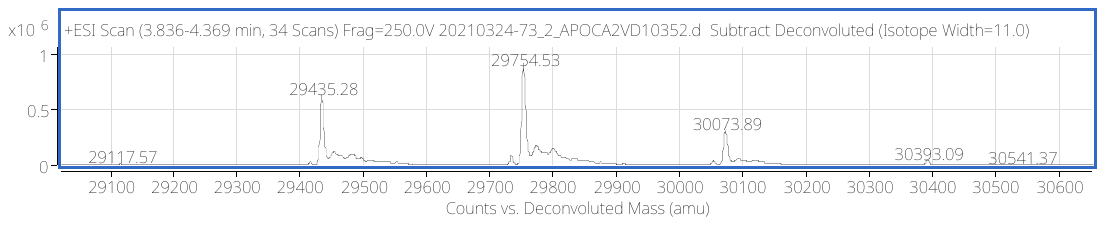

**C**
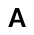

Figure S17. A) CA II mass spectra. B) CA II mass spectra after incubation with 10 eq. of compound **13**. C) CA II mass spectra after incubation with 10 eq. of compound **12**.

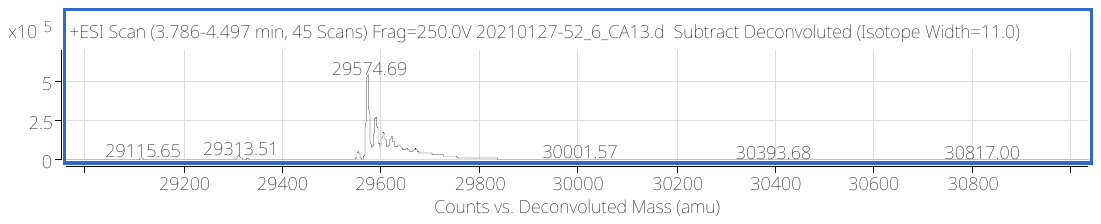

**A**
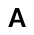

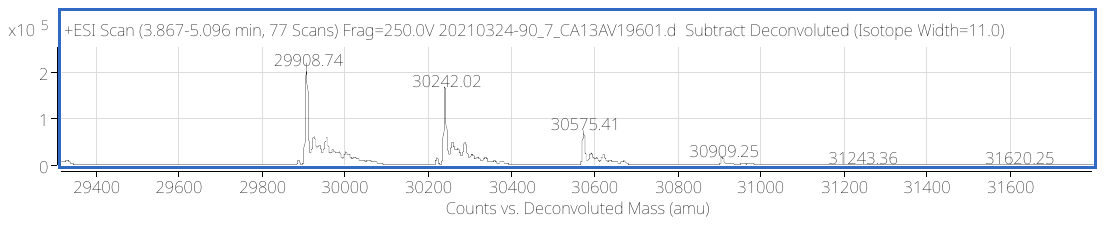

**B**
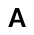

Figure S18. A) CA XIII mass spectra. B) CA XIII mass spectra after incubation with 10 eq. of compound **13**.

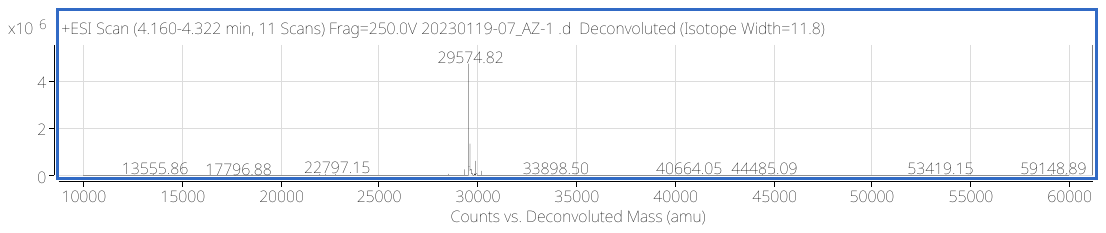

**A**
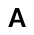

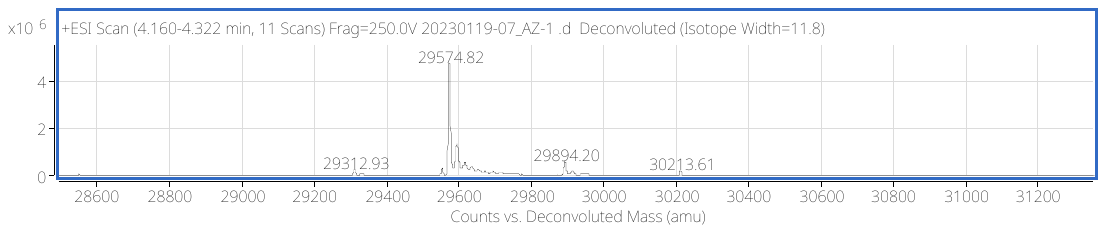

**B**

**C**

**E**

**D**

Figure S19. A) CA XIII Mass spectra; B) mass spectra ~3 minutes after addition of 1 eq. of **18**; C) mass spectra 40 minutes after addition of 1 eq. of **18**; D) mass spectra 2 hours after addition of 1 eq. of **18**; E) mass spectra 80 minutes after addition of 10 eq. of **18**.

**A**

**B**

Figure S20. A) CA IX mass spectra. B) CA IX mass spectra after incubation with 2 eq. of compound **22**.

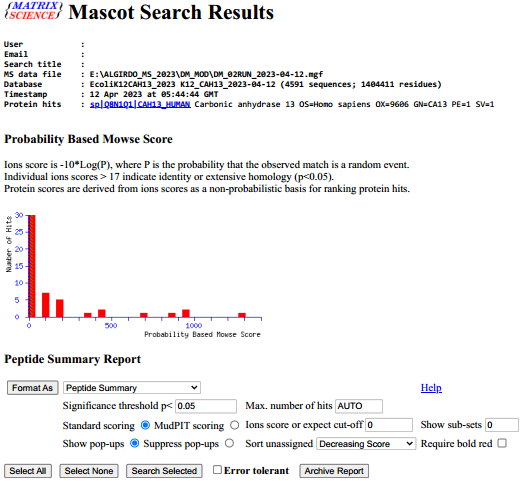

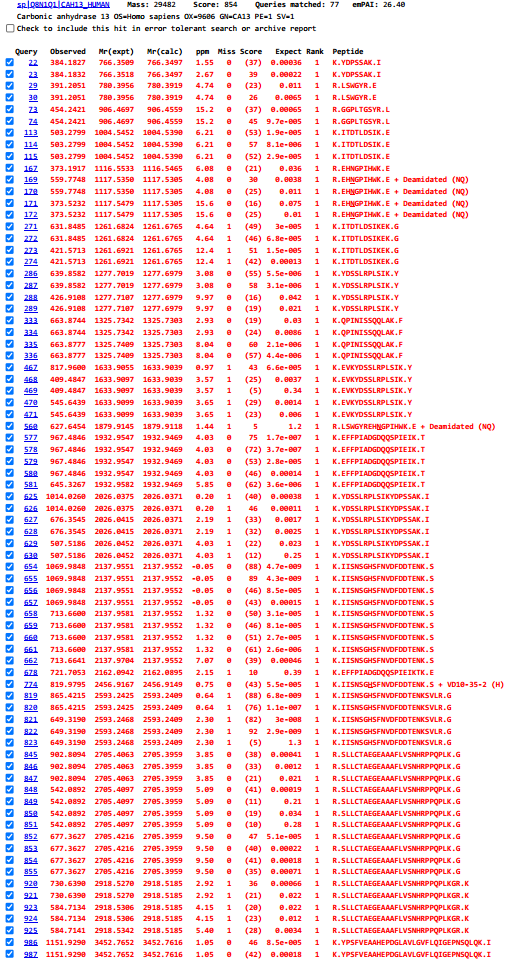

Figure S21. Identified peptides after trypsinisation of CAXIII protein in the presense of compound 12 (VD10-35-2). Proteomic data show only one binding site of compound 12 (VD10-35-2) in the peptide K.IISNSGHSFNVDFDDTENK.S+VD10-35-2 (H).

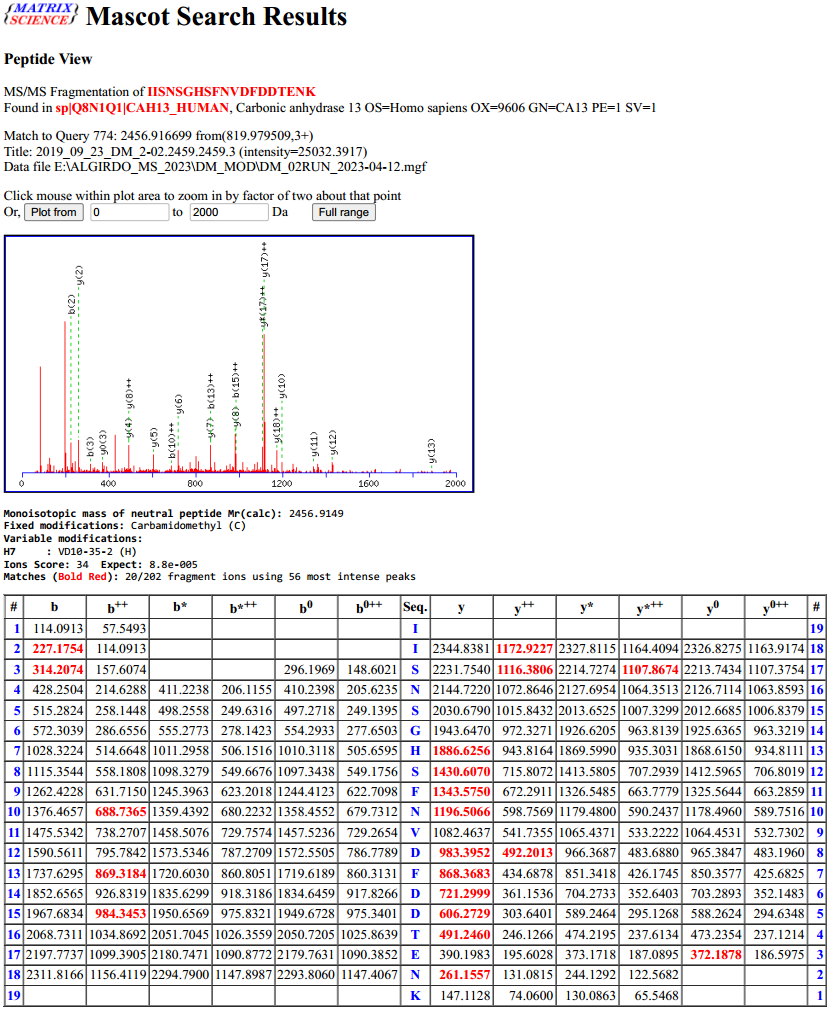

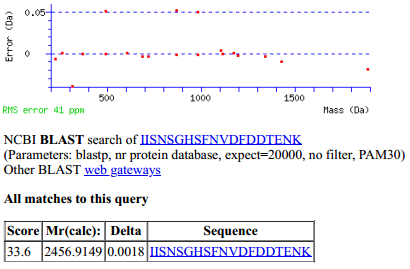

Figure S22. The peptide fragments that appear after fragmentation in the spectrometer. The masses of the actually detected peptide fragments are marked in red.

**2D NMR data**

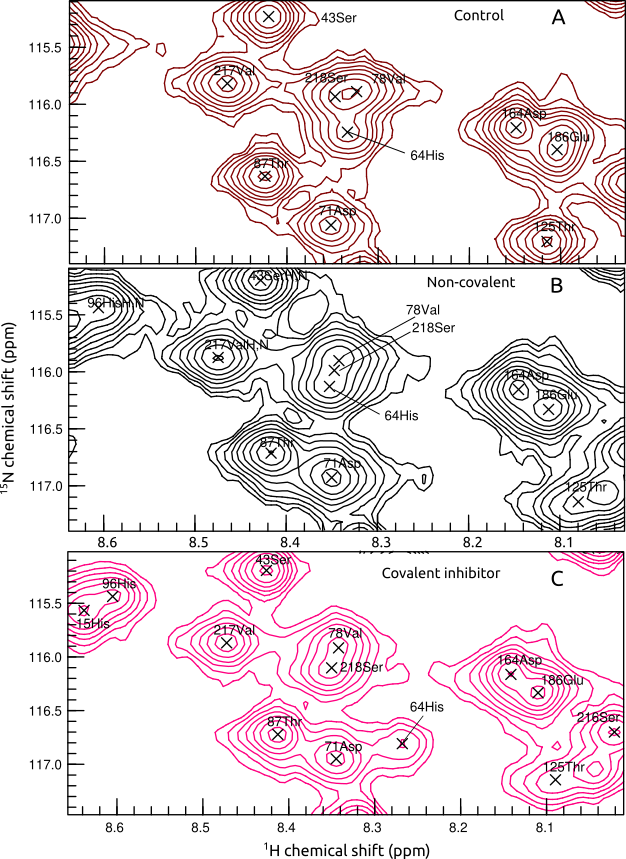

Figure S23. The change of His64 position of CAII in 1H-15N-HSQC spectrum: A – the control; B – after addition of non-covalent sulfonamide **1**; C – after addition of covalent (bottom panel) inhibitor **12**.

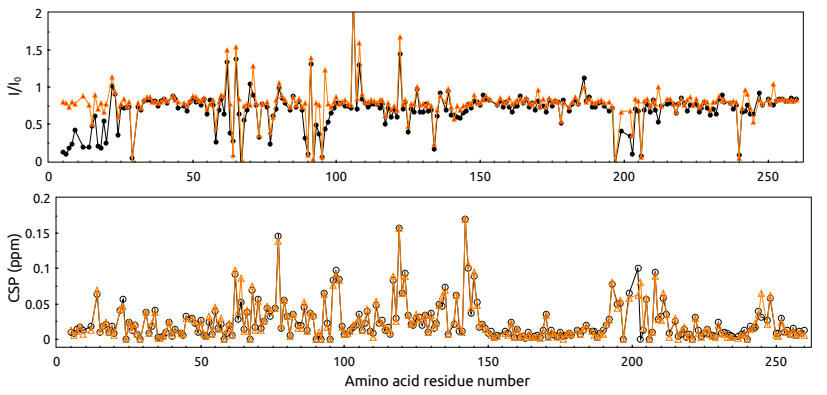

Figure S24. Intensity change and chemical shift perturbation (CSP) of recombinant human carbonic anhydrase peaks in the 1H-15N-HSQC spectrum after addition of non-covalent (orange) or covalent (black) inhibitors

**Fluorescent thermal shift assay (FTSA) data**

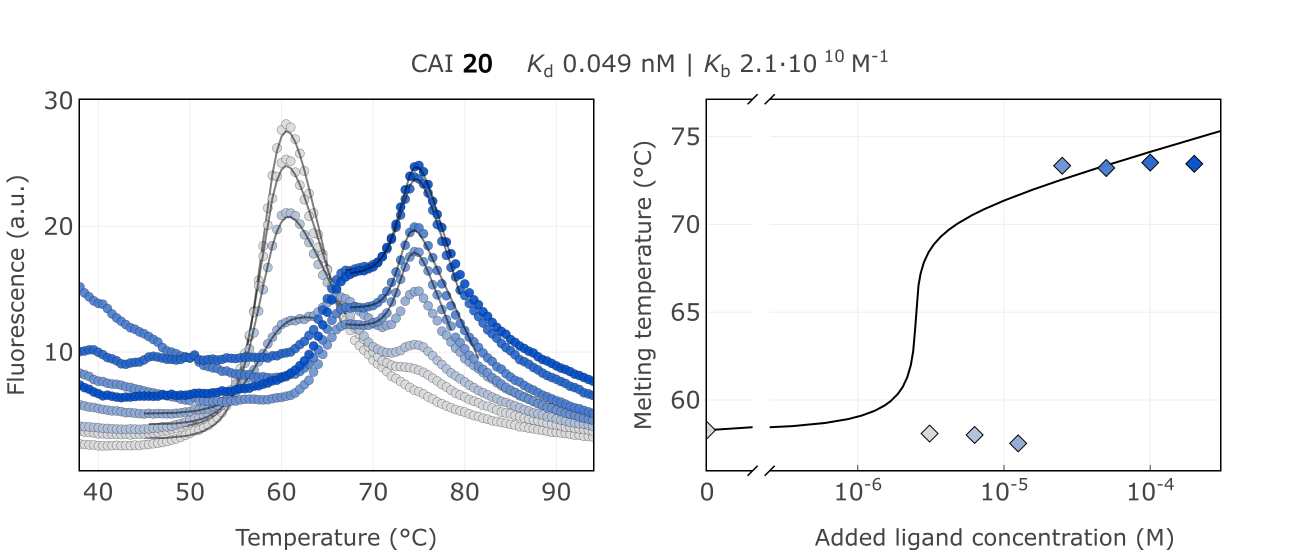

**B**

**A**

Figure S25. A) Raw FTSA data of compound **20** binding to CA I B) Enzyme melting temperature dependence on sulfonamide **20** concentration.

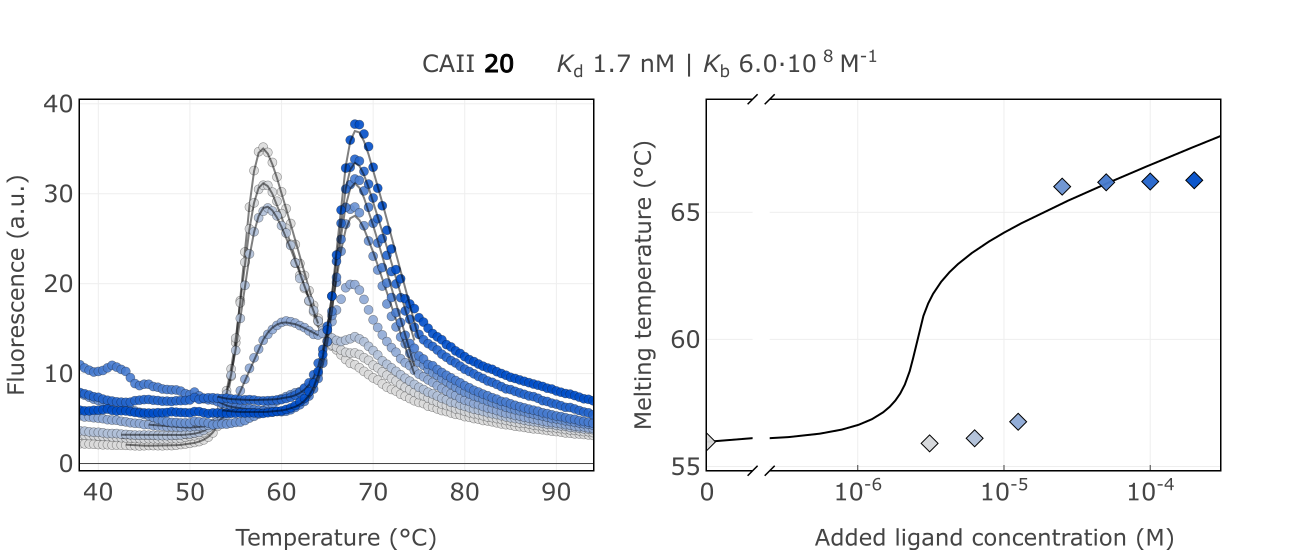

**B**

**A**

Figure S26. A) Raw FTSA data of compound **20** binding to CA II. B) Enzyme melting temperature dependence on sulfonamide **20** concentration.

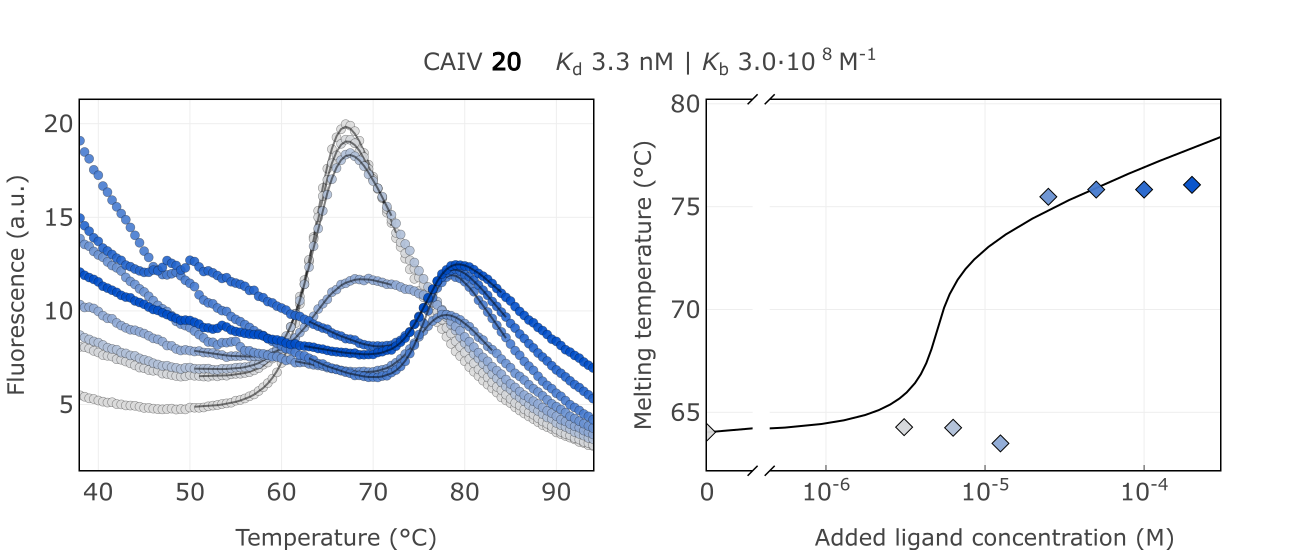

**B**

**A**

Figure S27. A) Raw FTSA data of compound **20** binding to CA IV. B) Enzyme melting temperature dependence on sulfonamide **20** concentration.

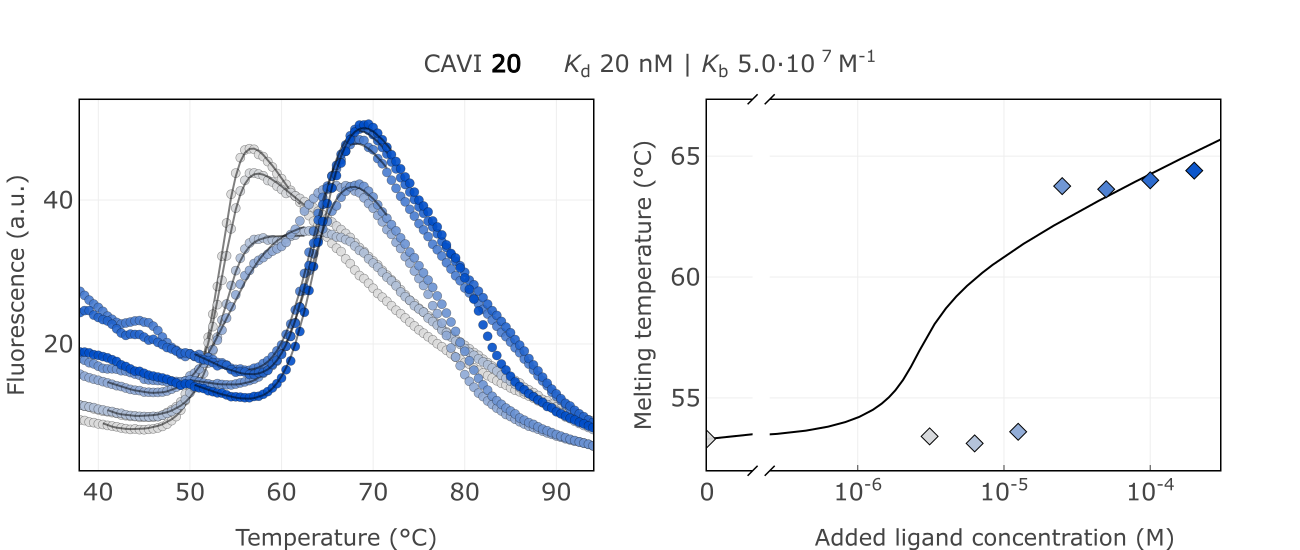

**B**

**A**

Figure S28. A) Raw FTSA data of compound **20** binding to CA VI. B) Enzyme melting temperature dependence on sulfonamide **20** concentration.

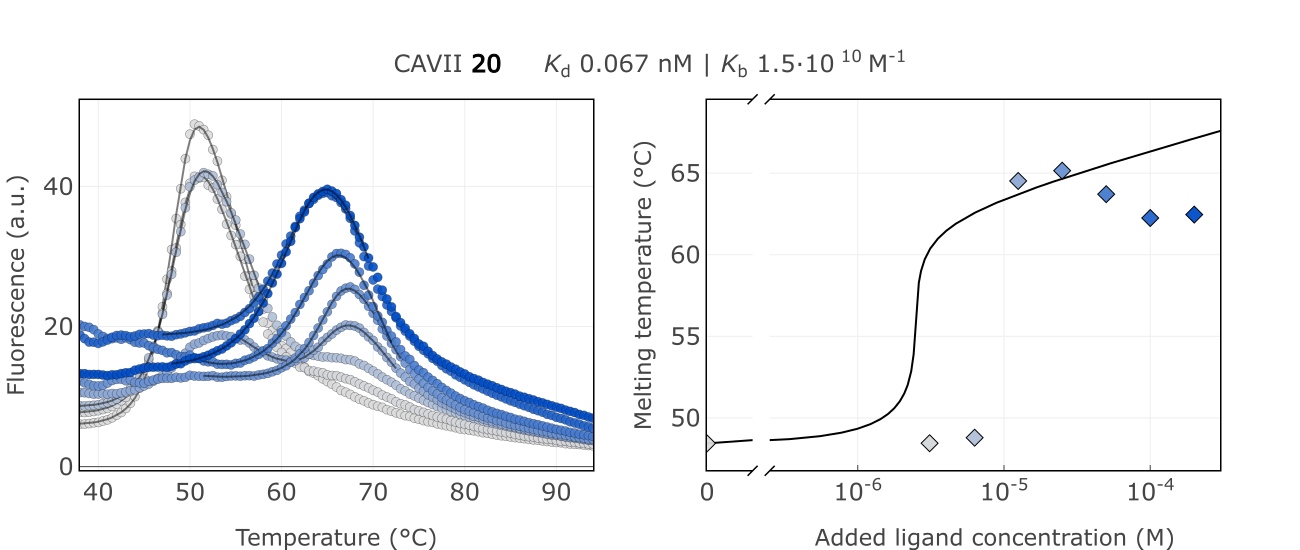

**B**

**A**

Figure S29. A) Raw FTSA data of compound **20** binding to CA VII. B) Enzyme melting temperature dependence on sulfonamide **20** concentration.

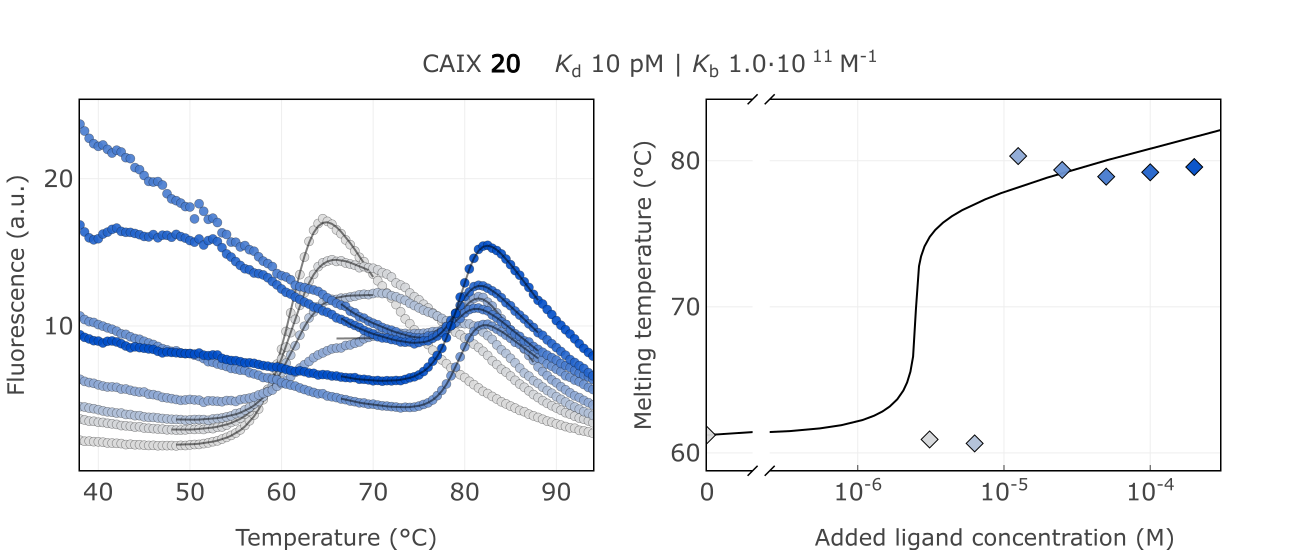

**B**

**A**

Figure S30. A) Raw FTSA data of compound **20** binding to CA IX. B) Enzyme melting temperature dependence on sulfonamide **20** concentration.

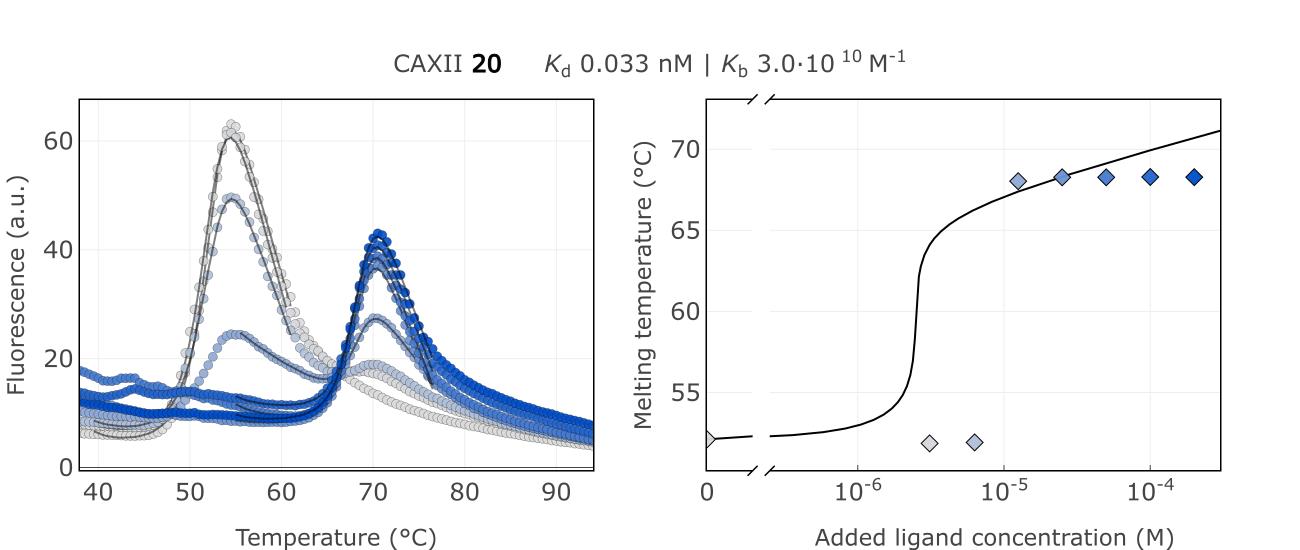

**B**

**A**

Figure S31. A) Raw FTSA data of compound **20** binding to CA XII. B) Enzyme melting temperature dependence on sulfonamide **20** concentration.

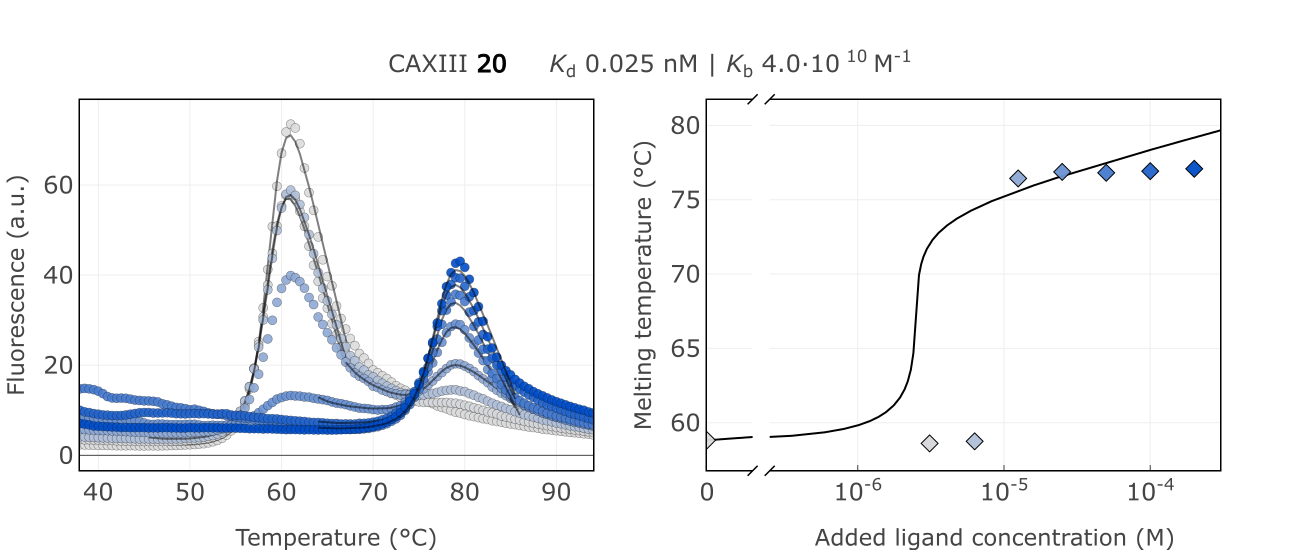

**B**

**A**

Figure S32. A) Raw FTSA data of compound **20** binding to CA XIII. B) Enzyme melting temperature dependence on sulfonamide **20** concentration.

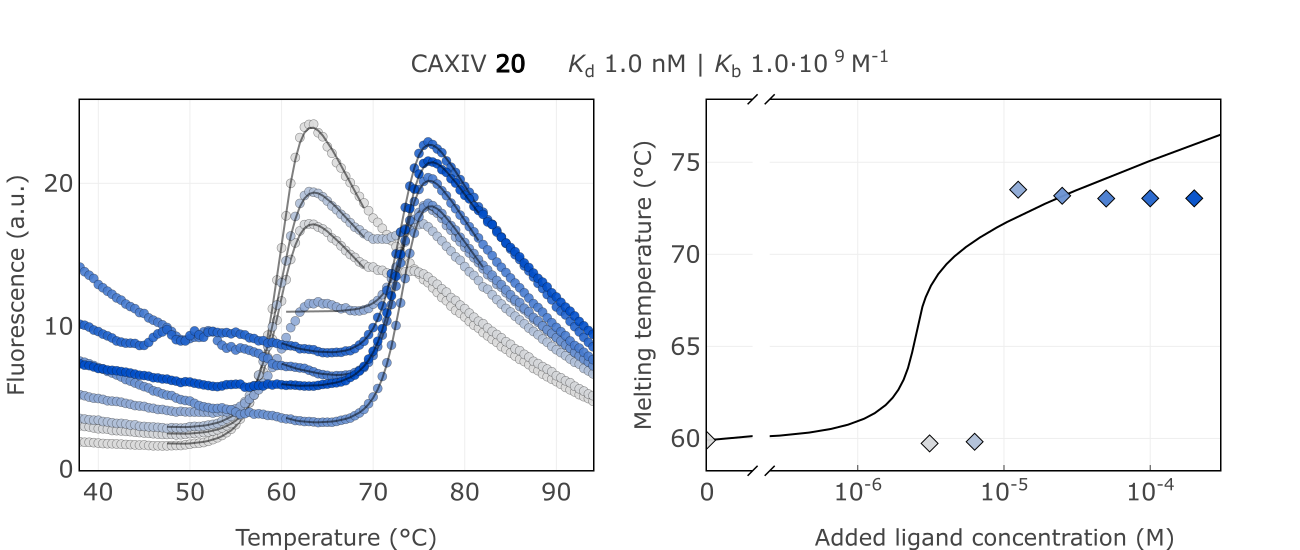

**B**

**A**

Figure S33. A) Raw FTSA data of compound **20** binding to CA XIV. B) Enzyme melting temperature dependence on sulfonamide **20** concentration.

**FTSA data, dependence on time**

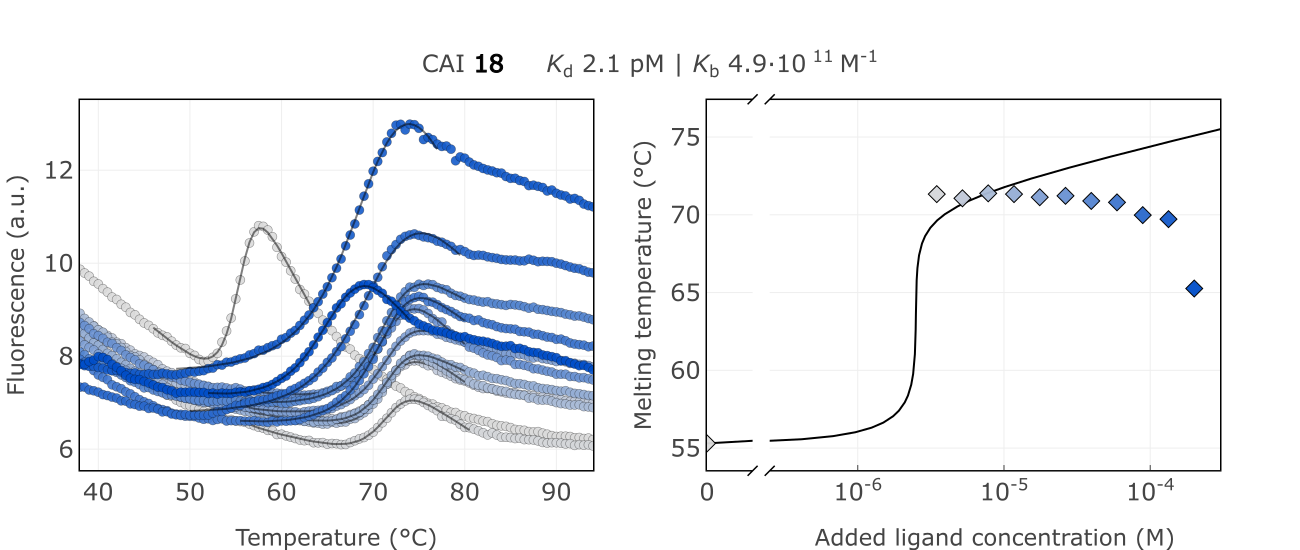

**B**

**A**

Figure S34. A) Raw FTSA data of compound **18** binding to CA I after 2 hour incubation period. B) Enzyme melting temperature dependence on sulfonamide **18** concentration.

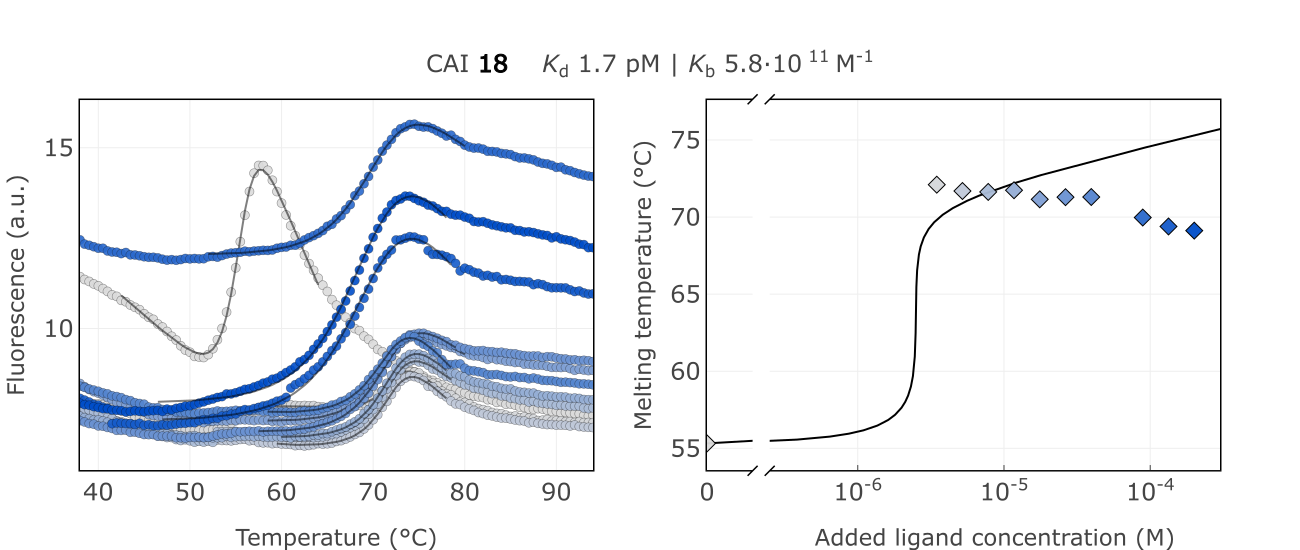

**B**

**A**

Figure S35. A) Raw FTSA data of compound **18** binding to CA I without incubating period. B) Enzyme melting temperature dependence on sulfonamide **18** concentration.

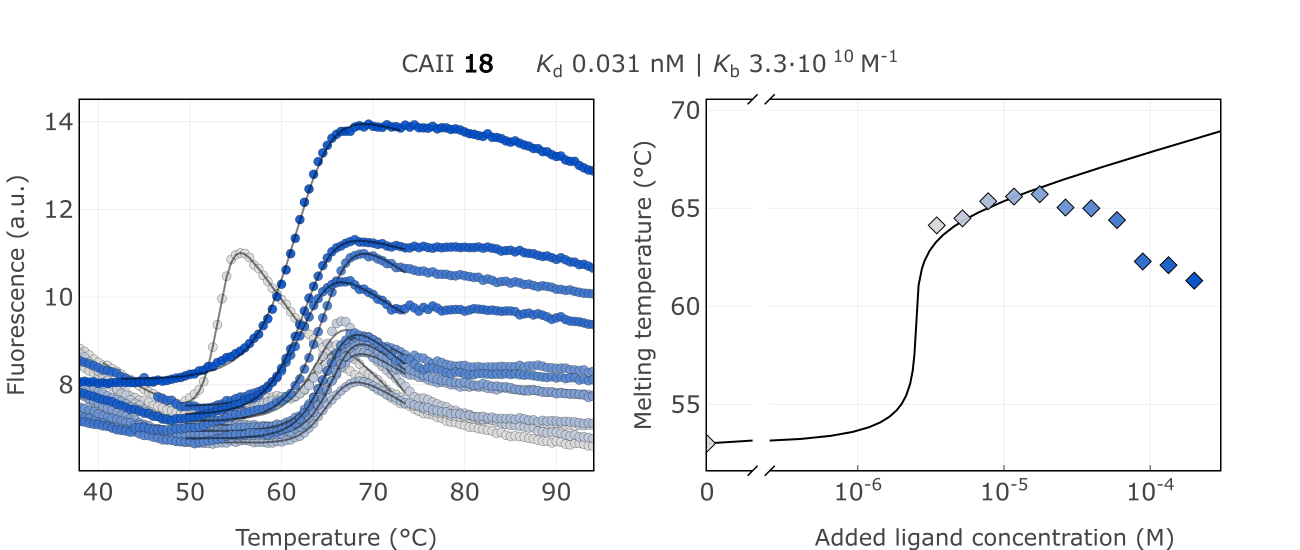

**B**

**A**

Figure S36. A) Raw FTSA data of compound **18** binding to CA II after 2 hour incubation period. B) Enzyme melting temperature dependence on sulfonamide **18** concentration.

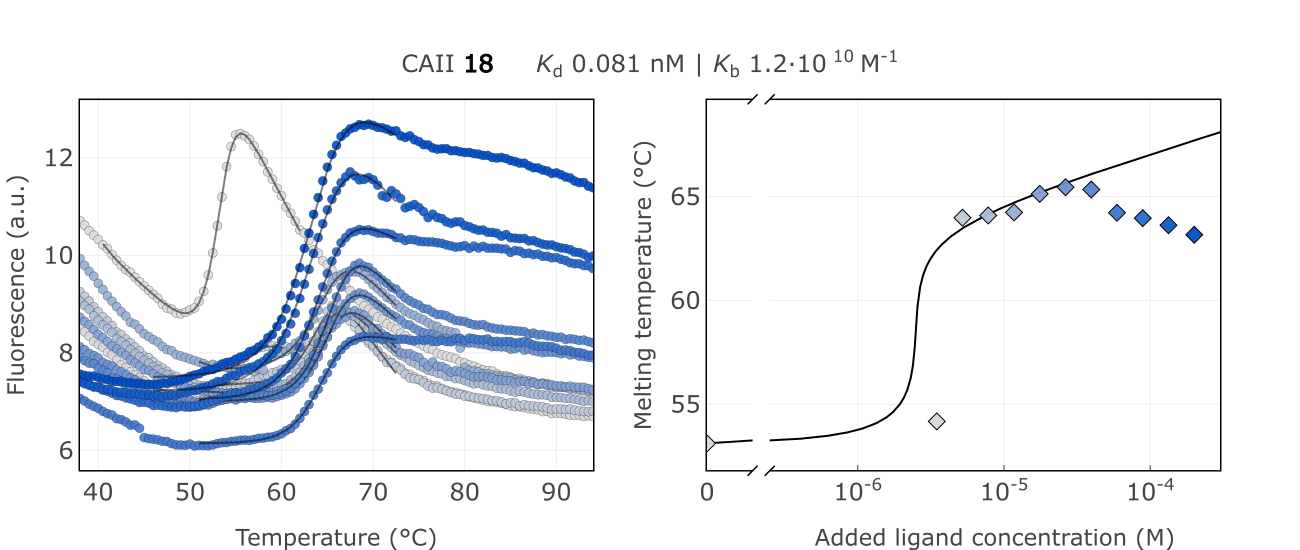

**B**

**A**

Figure S37. A) Raw FTSA data of compound **18** binding to CA II without incubating period. B) Enzyme melting temperature dependence on sulfonamide **18** concentration.

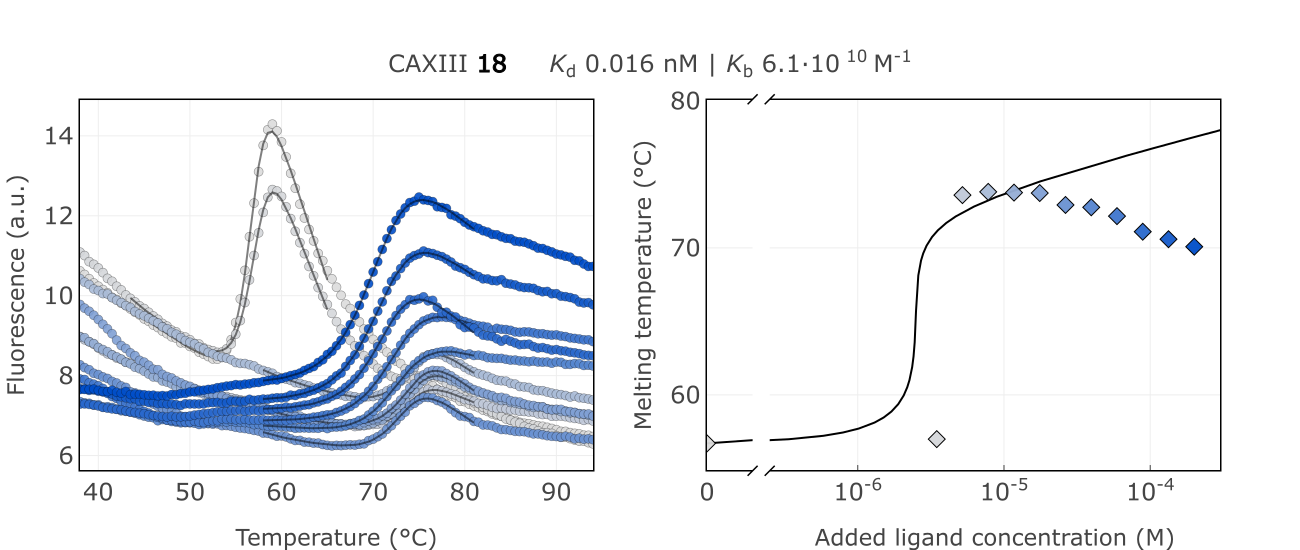

**B**

**A**

Figure S38. A) Raw FTSA data of compound **18** binding to CA XIII after 2 hour incubation period. B) Enzyme melting temperature dependence on sulfonamide **18** concentration.

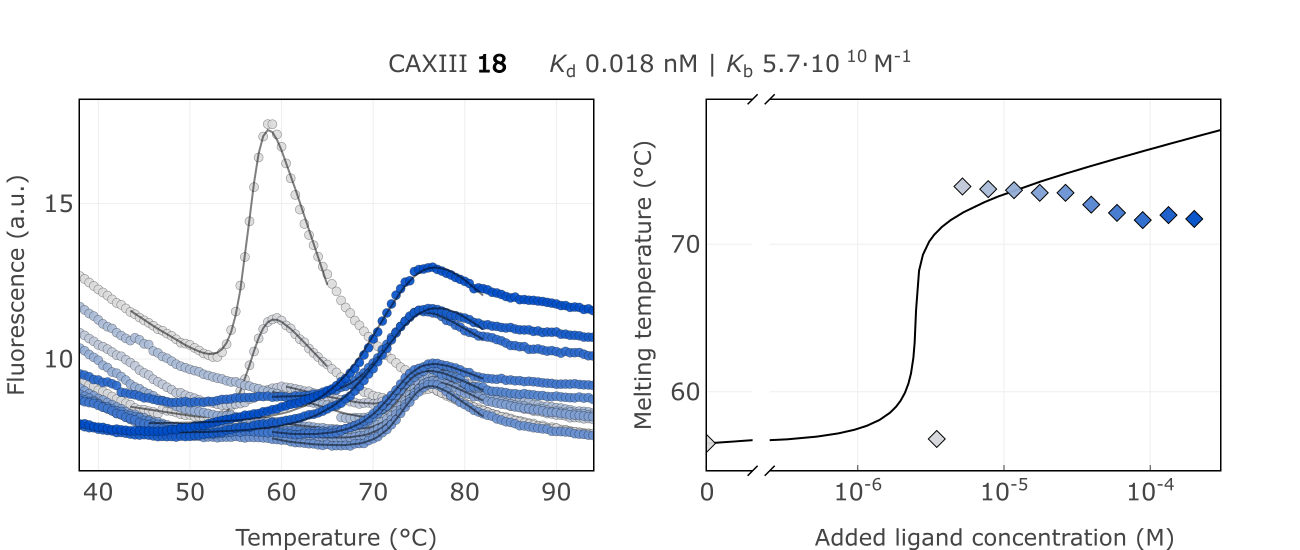

**B**

**A**

Figure S39. A) Raw FTSA data of compound **18** binding to CA XIII without incubating period. B) Enzyme melting temperature dependence on sulfonamide **18** concentration.

**Chemistry**

Compound numbering in the supplementary includes the intermediate compounds, thus numbers differ from the main manuscript. Compounds in the supplementary are marked with x. Compounds appearing in the main manuscript scheme 1 has the corresponding number in the brackets.

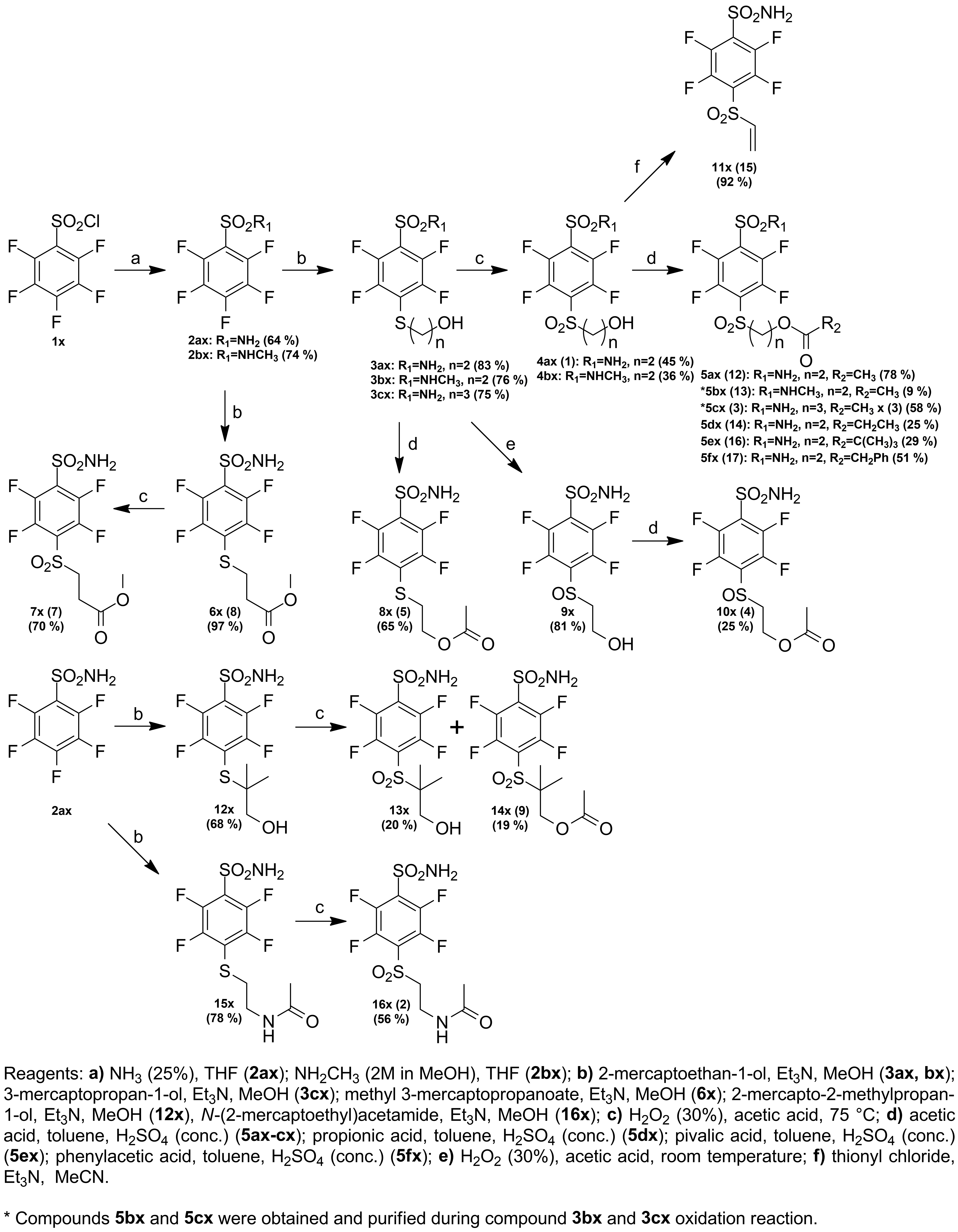

Scheme S1. Compound **5ax-fx**, **7x**, **8x**, **10x**, **11x, 14x** and **16x** synthesis.

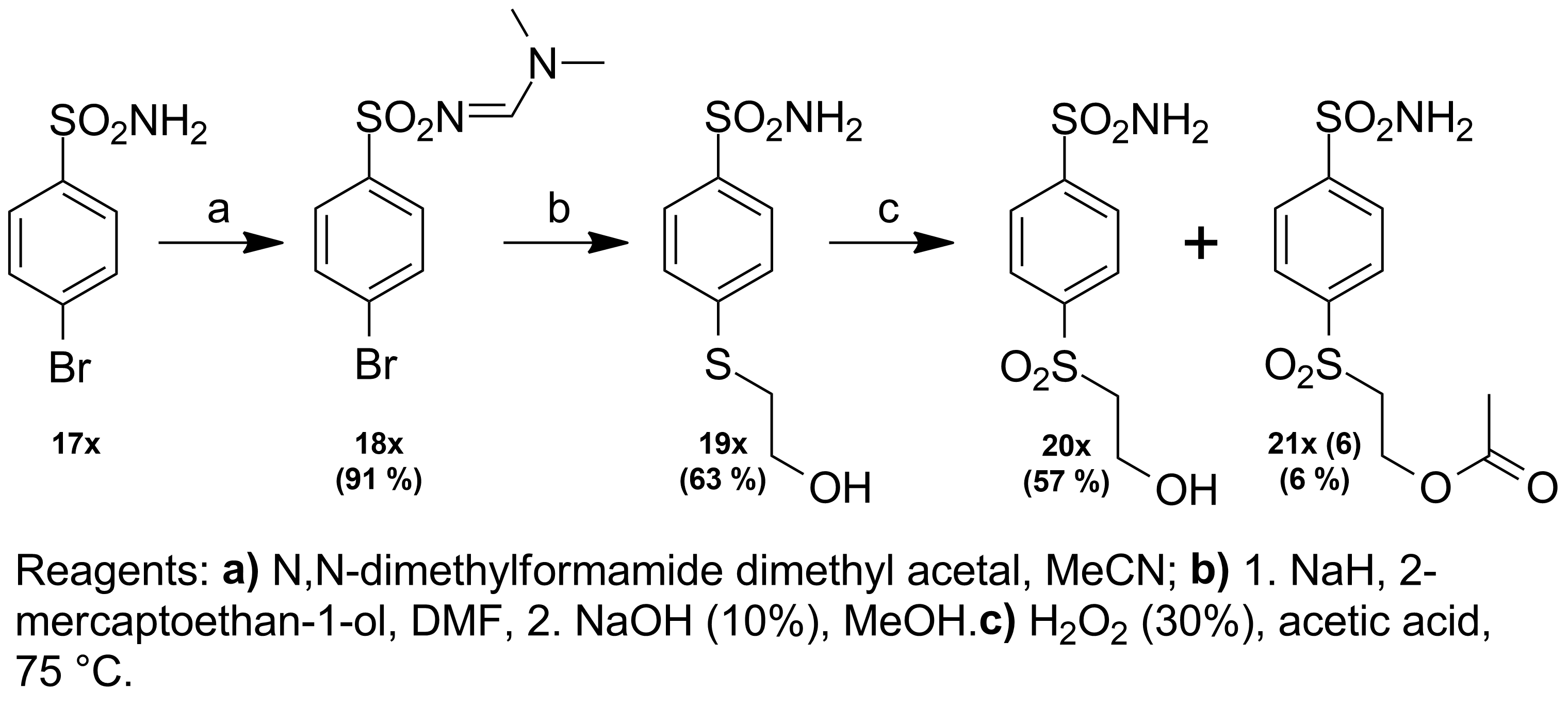

Scheme S2. Compound **21x** synthesis.

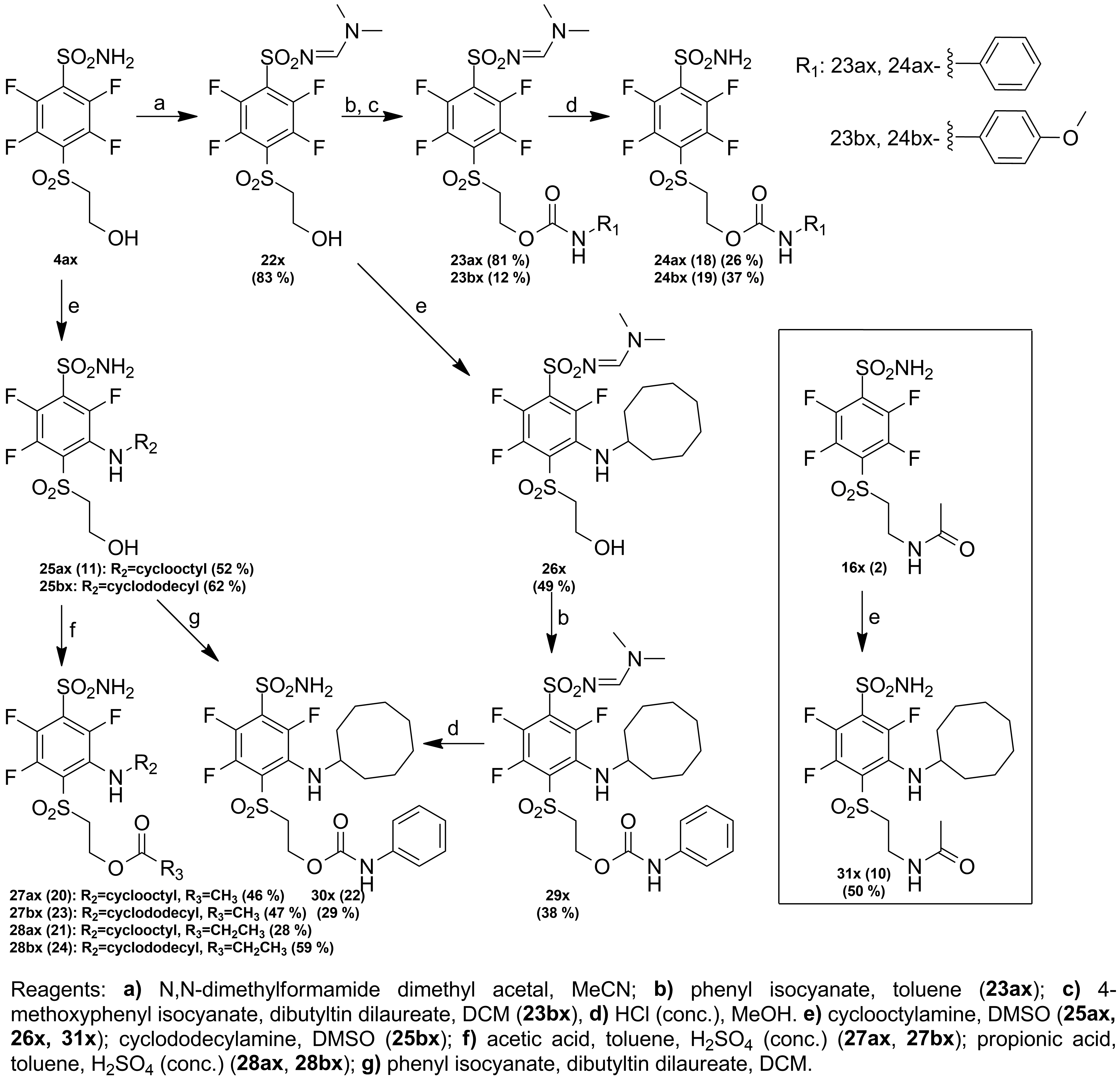

Scheme S3. Compound **24ax**, **24bx**, **27ax**, **27bx**, **28ax**, **28bx**, **30x** and **31x** synthesis.

All starting materials and reagents were commercial products or those which could be prepared according to known procedures. Melting points of the compounds were determined in open capillaries on a Thermo Scientific 9100 Series and are uncorrected. ^1^H and ^13^C NMR spectra were recorded on a Bruker spectrometer (400 and 100 MHz, respectively) in DMSO-D_6_ or CDCl_3_ using residual DMSO, CDCl_3_ signals (2.50 ppm, 7.26 ppm and 39.52 ppm, 77.16 ppm for ^1^H and ^13^C NMR spectra, respectively) as the internal standard. ^19^F NMR spectra were recorded on a Bruker spectrometer (376 MHz) with CFCl_3_ as an internal standard. TLC was performed with silica gel 60 F254 aluminum plates (Merck) and visualized with UV light. Column chromatography was performed using silica gel 60 (0.040-0.063 mm, Merck). High-resolution mass spectra (HRMS) were recorded on a Dual-ESI Q-TOF 6520 mass spectrometer (Agilent Technologies). Compound IUPAC names were generated with Chemdraw ultra 12.0.

2,3,4,5,6-pentafluorobenzenesulfonamide **2ax**

2,3,4,5,6-pentafluorobenzenesulfonamide (**2ax**) was prepared according to known procedure in literature (Dudutienė, V. et al. Bioorg. Med. Chem. 21, 2093–2106 (2013)). To a 100 ml cooled to -10 °C temperature THF, 2,3,4,5,6-pentafluorobenzenesulfonyl chloride (**1x**) (5 ml; 33.70 mmol; 1 eq.) was dissolved and NH_3_ (10 %) (10 ml) was slowly added dropwise while mixing as well as keeping reaction temperature under -10 °C. After reaction completion mixture was basified to pH 8-9 and left stirring for additional hour at room temperature. The product was recrystallized from H_2_O and white crystals were obtained. Yield: 5.33 g; (64%). Mp: 154-155 °C (close to the value in the literature (Dudutienė, V. *et al.* *Bioorg. Med. Chem.* 21, 2093–2106 (2013)), mp: 155-156 °C).

2,3,4,5,6-pentafluoro-N-methylbenzenesulfonamide **2bx**

Synthesis was described in (Baronas, D. et al. European Biophysics Journal 50, 993-1011 (2021)). To a 100 ml cooled to -10 °C temperature THF, 2,3,4,5,6-pentafluorobenzenesulfonyl chloride (**1x**) (2 ml; 13.47 mmol; 1 eq.) was dissolved and 2M methylamine in methanol (0.837 g; 26.9 mmol; 2 eq.) was slowly added dropwise while mixing as well as keeping reaction temperature under -10 °C. The reaction process was monitored using TLC and after reaction completion THF was evaporated under reduced pressure. The product was recrystallized from MeOH:H_2_O (1:4) and white crystals were obtained. Yield: 2.6 g; (74%). Mp: 95-97 °C (close to the value in the literature (Baronas, D. et al. European Biophysics Journal 50, 993-1011 (2021), mp: 96-97 °C).

2,3,5,6-tetrafluoro-4-((2-hydroxyethyl)thio)benzenesulfonamide **3ax**

2,3,5,6-tetrafluoro-4-((2-hydroxyethyl)thio)benzensulfonamide (**3ax**) was prepared according to known procedure in literature (Dudutienė, V. et al. Bioorg. Med. Chem. 21, 2093–2106 (2013)). 2,3,4,5,6-pentafluorobenzensulfonamide (**2ax**) (1.000 g; 4.05 mmol; 1 eq.), 2-mercaptoethanol (0.341 ml; 4.86 mmol; 1.2 eq.), Et_3_N (0.677 ml; 4.86 mmol; 1.2 eq.) were dissolved in MeOH (20 ml) and left to stir at room temperature over night. Next morning additional 2-mercaptoethanol (0.085 ml; 1.21 mmol; 0.3 eq.) and Et_3_N (0.169 ml; 1,21 mmol; 0.3 eq.) portions were added and reaction mixture was left to stir for 2 hours. After reaction completion the solvent was evaporated under reduced pressure and resultant preciptate was washed with H_2_O. Recrystalization was accomplished from H_2_O. Yield: 1.029 g; (83%). Mp: 111-112 °C (close to the value in the literature (Dudutienė, V. *et al.* *Bioorg. Med. Chem.* 21, 2093–2106 (2013)), mp: 111-112 °C).

2,3,5,6-tetrafluoro-4-((2-hydroxyethyl)thio)-N-methylbenzenesulfonamide **3bx**

2,3,5,6-tetrafluoro-4-((2-hydroxyethyl)thio)-N-methylbenzenesulfonamide (**3bx**) was prepared according to known procedure in the literature (2). 2,3,4,5,6-pentafluoro-N-methylbenzenesulfonamide (**2bx**) (0.500 g; 1.92 mmol; 1eq.), 2-mercaptoethanol (0.175 ml; 2.50 mmol; 1.3 eq.), Et_3_N (0.349 ml, 2.50 mmol, 1.3 eq.) were dissolved in MeOH (20 ml) and left to stir at room temperature for 20 hours. After reaction completion the solvent was evaporated under reduced pressure and resultant preciptate was washed with H_2_O. Recrystalization was accomplished from H_2_O. Yield: 0.464 g; (76%). Mp: 99 °C. ^1^H NMR (400 MHz, DMSO-d_6_, δ): 2.62 (3H, s, SO_2_NHCH_3_) 3.15 (2H, t, *J =* 6.1 Hz, SCH_2_), 3.59 (2H, q, *J =* 5.9 Hz, CH_2_O), 4.94 (1H, t, *J =* 5.4 Hz, OH), 8.42 (1H, s, NH). ^13^C NMR (100 MHz, DMSO-d_6_, δ): 28.28 (SO_2_NHCH_3_), 36.52 (SCH_2_, t, *J* (^19^F – ^13^C) = 3.0 Hz), 60.66 (CH_2_O), 118.23 (C1, t, *J* (^19^F – ^13^C) = 15.8 Hz), 119.99 (C4, t, *J* (^19^F – ^13^C) = 20.2 Hz), 143.01 (C2 and C6, ddt, ^1^*J* (^19^F – ^13^C) = 253.7 Hz, ^2^*J* (^19^F – ^13^C) = 16.9 Hz, ^3^*J* (^19^F – ^13^C) = 4.5 Hz), 146.41 (C3 and C5, ddt, ^1^*J* (^19^F – ^13^C) = 243.8 Hz, ^2^*J* (^19^F – ^13^C) = 15.4 Hz, ^3^*J* (^19^F – ^13^C) = 3.9 Hz). ^19^F NMR (376 MHz, DMSO-d_6_, δ): -132.79 – -132.97 (2F, m), -139.08 – -139.25 (2F, m).

2,3,5,6-tetrafluoro-4-((3-hydroxypropyl)thio)benzenesulfonamide **3cx**

2,3,4,5,6-pentafluorbenzensulfonamide (**2ax**) (0.400 g; 1.62 mmol), 3-mercaptopropan-1-ol (0.167 ml; 1.94 mmol; 1.2 eq.), Et_3_N (0.270 ml; 1.94 mmol; 1,2 eq.) were dissolved in MeOH (15 ml) and left to stir at room temperature over night. Next morning additional 3-mercaptopropan-1-ol (0.027 ml; 0.32 mmol; 0.2 eq.) and Et_3_N (0.044 ml; 0.32 mmol; 0.2 eq.) portions were added and reation mixture was left to stir for 3 hours. After reaction completion the solvent was evaporated under reduced pressure and product was purified by column chromatography silica gel, EtOAc/CHCl_3_ (1:1), Rf= 0.39). Yield 0.386 (75%). Mp: 148-149 °C. ^1^H NMR (400 MHz, DMSO-d_6_, δ): 1.65 (2H, p, *J* = 6.3 Hz, CH_2_CH_2_CH_2_), 3.09 (2H, t, *J* = 7.3 Hz, CH_2_S), 3.46 (2H, t, *J =* 6.0 Hz, CH_2_O), 8.39 (2H, s, SO_2_NH_2_). ^13^C NMR (100 MHz, DMSO-d_6_, δ): 30.77 (SCH_2_, t, *J (*^19^F-^13^C) = 3.1 Hz), 32.68 (CH_2_CH_2_CH_2_), 58.70 (CH_2_O), 118.67 (C1, t, *J (*^19^F – ^13^C) = 20.4 Hz), 122.46 (C4, t, *J (*^19^F – ^13^C) = 15.7 Hz), 142.54 (C2 and C6, ddt, ^1^*J (*^19^F – ^13^C) = 253.6 Hz, ^2^*J (*^19^F – ^13^C) = 16.9 Hz, ^3^*J (*^19^F – ^13^C) = 4.3 Hz), 146.48 (C3 and C5, ddt, ^1^*J (*^19^F – ^13^C) = 245.3 Hz, ^2^*J (*^19^F – ^13^C) = 13.9 Hz, ^3^*J (*^19^F – ^13^C) = 3.8 Hz). ^19^F NMR (376 MHz, DMSO-d_6_, δ): -133.23 – -133.38 (2F, m), -139.04 – -139.19 (2F, m). HRMS for C_9_H_9_F_4_NO_3_S_2_ [(M+H)^+^]: calc. 320.0033, found 320.0028.

2,3,5,6-tetrafluoro-4-((2-hydroxyethyl)sulfonyl)benzenesulfonamide **4ax** (**1**)

2,3,5,6-tetrafluoro-4-((2-hydroxyethyl)sulfonyl)benzenesulfonamide (**4ax** (**1**)) was prepared according to known procedure in literature (3). 2,3,5,6-tetrafluoro-4-((2-hydroxyethyl)thio)benzensulfonamide (**3ax**) (1.542 g; 5.06 mmol; 1 eq.) was dissolved in acetic acid (30 ml) and heated at 75 °C temperature for 18 hours. H_2_O_2_ (30%) was added by portions (0.1 ml) every 30 minutes (overall 3.6 ml) until complete starting material conversion. Afterwards, the solvent was evaporated under reduced pressure and the product was purified by column chromatography (silica gel, EtOAc/CHCl_3_ (1:1), Rf= 0.30). Yield: 0.768 g; (45%). Mp: 138-139 °C (close to the value in the literature (Dudutienė, V. *et al.* *Bioorg. Med. Chem.* 21, 2093–2106 (2013)), mp: 139-140 °C).

**Compounds 5bx (13) and 4bx were synthesized during compound 3bx oxidation.**

2,3,5,6-tetrafluoro-4-((2-hydroxyethyl)thio)-N-methylbenzenesulfonamide (**3bx**) (0.300 g, 0.94 mmol) was dissolved in acetic acid (10 ml) and heated at 75 °C temperature for 21 hours. H_2_O_2_ (30%) was added by portions (0.1 ml) every hour (overall 2.1 ml) until complete starting material conversion. Afterwards, the solvent was evaporated under reduced pressure and products were purified by column chromatography (silica gel, EtOAc/CHCl_3_ (1:1), Rf**_4bx_**= 0.53, Rf**_5bx_**= 0.79)

2,3,5,6-tetrafluoro-4-((2-hydroxyethyl)sulfonyl)-N-methylbenzenesulfonamide **4bx**

Yield: 0.119 g; (36%). Mp: 201 °C. ^1^H NMR (400 MHz, DMSO-d_6_, δ): 2.65 (3H, s, SO_2_NHCH_3_), 3.76 (2H, t, *J =* 5.6 Hz, SO_2_CH_2_), 3.86 (2H, q, *J =* 5.2 Hz, CH_2_O), 4.97 (1H, t, *J =* 5.3 Hz, OH), 8.68 (1H, s, NH). ^13^C NMR (100 MHz, DMSO-d_6_, δ): 28.31 (SO_2_NHCH_3_), 55.07 (SO_2_CH_2_), 59.54 (CH_2_O), 122.97 (C1, t, *J* (^19^F – ^13^C) = 15.0 Hz), 123.86 (C4, t, *J* (^19^F – ^13^C) = 15.7 Hz), 143.35 (C2 and C6, dd, ^1^*J* (^19^F – ^13^C) = 255.2 Hz, ^2^*J* (^19^F – ^13^C) = 15.7 Hz), 144.34 (C3 and C5, dd, ^1^*J* (^19^F – ^13^C) = 244.6 Hz, ^2^*J* (^19^F – ^13^C) = 13.3 Hz). ^19^F NMR (376 MHz, DMSO-d_6_, δ): -136.00 (2F, dd, ^1^*J =* 26.2 Hz, ^2^*J* = 12.7 Hz), -136.61 – -136.77 (2F, m). HRMS for C_9_H_9_F_4_NO_5_S_2_ [(M-H)^-^]: calc. 349.9786, found 349.9785.

2-((2,3,5,6-tetrafluoro-4-(N-methylsulfamoyl)phenyl)sulfonyl)ethyl acetate **5bx** (**13**)

Yield: 0.035 g; (9%). Mp: 143 °C. ^1^H NMR (400 MHz, DMSO-d_6_, δ): 1.85 (3H, s, COCH_3_), 2.65 (3H, d, *J =* 4.6 Hz, SO_2_NHCH_3_), 4.01 (2H, t, *J =* 5.4 Hz, SO_2_CH_2_), 4.41 (2H, t, *J =* 5.7 Hz, CH_2_O), 8.73 (1H, q, *J =* 4.6 Hz, NH). ^13^C NMR (100 MHz, DMSO-d_6_, δ): 20.11 (COCH_3_), 28.32 (SO_2_NHCH_3_), 55.95 (SO_2_CH_2_), 57.21 (CH_2_O), 121.74 (C1, t, *J* (^19^F – ^13^C) = 14.9 Hz), 124.46 (C4, t, *J* (^19^F – ^13^C) = 16.0 Hz), 143.64 (C2 and C6, d, ^1^*J* (^19^F – ^13^C) = 255.6 Hz), 144.57 (C3 and C5, d, ^1^*J* (^19^F – ^13^C) = 264.6 Hz), 169.56 (OC(O)). ^19^F NMR (376 MHz, DMSO-d_6_, δ): -135.37 – -135.56 (2F, m), -136.09 – -136.30 (2F, m). HRMS for C_11_H_11_F_4_NO_6_S_2_ [(M-H)^-^]: calc. 391.9891, found 391.9891.

2-((2,3,5,6-tetrafluoro-4-sulfamoylphenyl)sulfonyl)ethyl acetate **5ax** (**12**)

2,3,5,6-tetrafluoro-4-((2-hydroxyethyl)sulfonyl)benzensulfonamide (**4ax** (**1**)) (0.153 g; 0.45 mmol), acetic acid (0.260 ml; 4.54 mmol; 10 eq.) and one drop of H_2_SO_4_ (conc.) were dissolved in toluene (25 ml) and refluxed for 3 hours. The resulting mixture was cooled to 5 °C and white crystals were filtered. The product was purified by column chromatography (silica gel, EtOAc/CHCl_3_ (1:1), Rf= 0.40). Yield: 0.134 g; (78%). Mp: 153-154 °C. ^1^H NMR (400 MHz, DMSO-d_6_, δ): 1.83 (3H, s, CH_3_), 4.02 (2H, t, *J =* 5.3 Hz, SO_2_CH_2_), 4.40 (2H, t, *J =* 5.4 Hz, CH_2_O), 8.69 (2H, s, SO_2_NH_2_). ^13^C NMR (100 MHz, DMSO-d_6_, δ): 20.07 (CH_3_), 55.91 (SO_2_CH_2_), 57.28 (CH_2_O), 121.33 (C1, t, *J* (^19^F – ^13^C) = 15.3 Hz), 127.86 (C4, t, *J* (^19^F – ^13^C) = 15.7 Hz), 142.23 (C2 and C6, dd, ^1^*J* (^19^F – ^13^C) = 252.0 Hz, ^2^*J* (^19^F – ^13^C) = 20.2 Hz), 144.33 (C3 and C5, dd, ^1^*J* (^19^F – ^13^C) = 259.5 Hz, ^2^*J* (^19^F – ^13^C) = 18.6 Hz), 169.52 (OC(O)). ^19^F NMR (376 MHz, DMSO-d_6_, δ): -135.60 – -135.78 (2F, m), -136.60 – -136.78 (2F, m). HRMS for C_10_H_9_F_4_NO_6_S_2_ [(M-H)^-^]: calc. 377.9735, found 377.9735.

2,3,5,6-tetrafluor-4-[2-(acetil)propilsulfonil]benzensulfonamidas **5cx** (**3**)

2,3,5,6-tetrafluoro-4-((3-hydroxypropyl)thio)benzenesulfonamide (**3cx**) (0.378 g; 0.95 mmol) was dissolved in acetic acid (10 ml) and heated at 75 °C temperature for 18 hours. H_2_O_2_ (30%) was added by portions (0.1 ml) every 30 minutes (overall 2.1 ml) until complete starting material conversion. Afterwards, the solvent was evaporated under reduced pressure and product was purified by column chromatography (silica gel, EtOAc/CHCl_3_ (1:1), Rf= 0.64). Yield: 0.216 g; (58%). Mp: 188 °C. ^1^H NMR (400 MHz, DMSO-d_6_, δ): 2.00 (3H, s, CH_3_), 2.03 (2H, p, *J* = 7.8 Hz, CH_2_CH_2_CH_2_) 3.66 (2H, t, *J* = 7.8 Hz CH_2_SO_2_), 4.08 (2H, t, *J* = 6.4 Hz, CH_2_O), 8.64 (2H, s, SO_2_NH_2_). ^13^C NMR (100 MHz, DMSO-d_6_, δ): 20.66 (CH_3_), 21.39 (CH_2_CH_2_CH_2_), 53.71 (SO_2_CH_2_), 61.58 (CH_2_O), 120.30 (C1, t, *J* (^19^F – ^13^C) = 15.0 Hz), 127.74 (C4, t, *J* (^19^F – ^13^C) = 15.4 Hz), 142.99 (C2 and C6, d, *J* (^19^F – ^13^C) = 254.1 Hz), 144.46 (C3 and C5, d, *J* (^19^F – ^13^C) = 261.0 Hz), 170.39 (OC(O)). ^19^F NMR (376 MHz, DMSO-d_6_, δ): -135,62 – -135,82 (2F, m), -136,35 – -136,58 (2F, m). HRMS for C_11_H_11_F_4_NO_6_S_2_ [(M-H)^-^]: calc. 391.9891, found 391.891.

2-((2,3,5,6-tetrafluoro-4-sulfamoylphenyl)sulfonyl)ethyl propionate **5dx** (**14**)

2,3,5,6-tetrafluoro-4-((2-hydroxyethyl)sulfonyl)benzensulfonamide (**4ax** (**1**)) (0.152 g; 0.451 mmol), propionic acid (2ml) and three drops of H_2_SO_4_ (conc.) were dissolved in toluene (25 ml) and refluxed for 1 hour. The resulting mixture was washed with brine (3x10 ml). The organic phase was dried using anhydrous Na_2_SO_4_ and evaporated under reduced pressure. The product was purified by column chromotography (silica gel, EtOAc/CHCl_3_ (1:1), Rf= 0.74). Yield: 0.0448 g; (25%). Mp: 109-111 °C. ^1^H NMR (400 MHz, DMSO-d_6_, δ): 0.93 (3H, t, *J =* 7.5 Hz, CH_2_CH_3_), 2.08 (2H, q, *J =* 7.5 Hz, CH_2_CH_3_), 4.03 (2H, t, *J =* 5.6 Hz, SO_2_CH_2_), 4.42 (2H, t, *J =* 5.6 Hz, CH_2_O), 8.69 (2H, s, SO_2_NH_2_). ^13^C NMR (100 MHz, DMSO-d_6_, δ): 8.63 (CH_3_), 26.36 (CH_2_CH_3_), 56.00 (SO_2_CH_2_), 57.22 (CH_2_O), 121.32 (C1, t, *J* (^19^F – ^13^C) = 14.6 Hz), 127.85 (C4, t, *J* (^19^F – ^13^C) = 15.2 Hz), 142.90 (C2 and C6, dd, ^1^*J* (^19^F – ^13^C) = 256.1 Hz, ^2^*J* (^19^F – ^13^C) = 18.8 Hz), 144.36 (C2 and C5, dd, ^1^*J* (^19^F – ^13^C) = 259.1 Hz, ^2^*J* (^19^F – ^13^C) = 20.4 Hz), 172.82 (OC(O)). ^19^F NMR (376 MHz, DMSO-d_6_, δ): -135.61 – -135.83 (2F, m), -136.58 – -136.79 (2F, m). HRMS for C_11_H_11_F_4_NO_6_S_2_ [(M-H)^-^]: calc. 391.9891, found 391.9892.

2-((2,3,5,6-tetrafluoro-4-sulfamoylphenyl)sulfonyl)ethyl pivalate **5ex** (**16**)

2,3,5,6-tetrafluoro-4-((2-hydroxyethyl)sulfonyl)benzensulfonamide (**4ax** (**1**)) (0.050 g; 0.015 mmol), pivalic acid (0.076 g; 0.074 mmol; 5 eq.) and three drops of H_2_SO_4_ (conc.) were dissolved in toluene (10 ml) and refluxed for 30 minutes. The resulting mixture was washed with brine (3x10 ml). The organic phase was dried using anhydrous Na_2_SO_4_ and evaporated under reduced pressure. The product was purified by column chromotography (silica gel, EtOAc/CHCl_3_ (1:1), Rf= 0.62). Yield: 0.018 g; (29%). Mp: 67-68 °C. ^1^H NMR (400 MHz, DMSO-d_6_, δ): 1.02 (9H, s, C(CH_3_)_3_), 4.04 (2H, t, *J =* 5.4 Hz, SO_2_CH_2_), 4.42 (2H, t, *J =* 5.5 Hz, CH_2_O), 8.70 (2H, s, SO_2_NH_2_). ^13^C NMR (100 MHz, DMSO-d_6_, δ): 26.46 (C(CH_3_)_3_), 38.00 (C(CH_3_)_3_), 56.32 (SO_2_CH_2_), 57.35 (CH_2_O), 121.12 (C1, t, *J* (^19^F – ^13^C) = 14.9 Hz), 127.94 (C4, t, *J* (^19^F – ^13^C) = 15.4 Hz), 143.01 (C2 and C6, dd, ^1^*J* (^19^F – ^13^C) = 257.3 Hz, ^2^*J* (^19^F – ^13^C) = 13.4 Hz), 144.35 (C3 and C5, dd, ^1^*J* (^19^F – ^13^C) = 256.9 Hz, ^2^*J* (^19^F – ^13^C) = 16.7 Hz), 176.84 (OC(O)). ^19^F NMR (376 MHz, DMSO-d_6_, δ): -135.72 – -135.90 (2F, m), -136.46 – -136.63 (2F, m). HRMS for C_13_H_15_F_4_NO_6_S_2_ [(M-H)^-^]: calc. 420.0204, found 420.0202.

2-((2,3,5,6-tetrafluoro-4-sulfamoylphenyl)sulfonyl)ethyl 2-phenylacetate **5fx** (**17**)

2,3,5,6-tetrafluoro-4-((2-hydroxyethyl)sulfonyl)benzensulfonamide (**4ax** (**1**)) (0.041 g; 0.122 mmol; 1 eq.), phenylacetic acid (0.075 g; 0.551 mmol; 5 eq.) and three drops of H_2_SO_4_ (conc.) were dissolved in toluene (15 ml) and refluxed for 1 hour. The resulting mixture was washed with brine (3x10 ml). The organic phase was dried using anhydrous Na_2_SO_4_ and evaporated under reduced pressure. The product was purified by column chromatography (silica gel, EtOAc/CHCl_3_ (1:1), Rf= 0.71). Yield: 0.0282 g; (51%). Mp: 77-79 °C. ^1^H NMR (400 MHz, DMSO-d_6_, δ): 3.49 (2H, s, OC(O)CH_2_), 4.05 (2H, t, *J =* 5.5 Hz, SO_2_CH_2_), 4.46 (2H, t, *J =* 5.5 Hz, CH_2_O), 7.19 (2H, d, *J =* 6.9 Hz, phenyl), 7.26 (1H, t, *J =* 7.2 Hz, phenyl), 7,31 (2H, t, *J =* 7.1 Hz, phenyl), 8.69 (2H, s, SO_2_NH_2_). ^13^C NMR (100 MHz, DMSO-d_6_, δ): 40.68 (COCH_2_), 55.96 (SO_2_CH_2_), 57.55 (CH_2_O), 121.24 (C1, t, *J* (^19^F – ^13^C) = 14.8 Hz), 126.95 (C4 of phenyl), 127.90 (C4, t, *J* (^19^F – ^13^C) = 15.7 Hz), 128.36 (phenyl), 142.96 (C2 and C6, dd, ^1^*J* (^19^F – ^13^C) = 259.0 Hz, ^2^*J* (^19^F – ^13^C) = 17.7 Hz), 144.33 (C3 and C5, dd, ^1^*J* (^19^F – ^13^C) = 254.2 Hz, ^2^*J* (^19^F – ^13^C) = 18,6 Hz), 170.50 (OC(O)). ^19^F NMR (376 MHz, DMSO-d_6_, δ): -135.53 – -135.70 (2F, m), -136.43 – -136.60 (2F,m). HRMS for C_16_H_13_F_4_NO_6_S_2_ [(M-H)^-^]: calc. 454.0048, found 454.0044.

Methyl 3-((2,3,5,6-tetrafluoro-4-sulfamoylphenyl)thio)propanoate **6x** (**8**)

2,3,4,5,6-pentafluorbenzensulfonamide (**2ax**) (0.675 g; 2.73 mmol), methyl 3-mercaptopropanoate (0.394 ml, 3.55 mmol, 1.3 eq.), Et_3_N (0.456 ml, 3.28 mmol, 1.2 eq.) were dissolved in MeOH (15 ml) and left to stir at room temperature for 0.5 hour. After reaction completion the solvent was evaporated under reduced pressure and resultant preciptate was washed with H_2_O. Recrystalization was accomplished from H_2_O. Yield: 0.921 g; (97%). Mp: 124-125 °C. ^1^H NMR (400 MHz, DMSO-d_6_, δ): 2.66 (2H, t, *J =* 6.8 Hz, CH_2_C(O)), 3.22 (2H, t, *J =* 6.8 Hz, SCH_2_), 3.56 (3H, s, CH_3_O), 8.43 (2H, s, SO_2_NH_2_). ^13^C NMR (100 MHz, DMSO-d_6_, δ): 29.22 (SCH_2_, t, *J* (^19^F – ^13^C) = 2.8 Hz), 34.42 (CH_2_C(O)O), 51.54 (CH_3_O), 117.71 (C1, t, *J* (^19^F – ^13^C) = 20.5 Hz), 122.86 (C4, t, *J* (^19^F – ^13^C) = 15.6 Hz), 142.52 (C2 and C6, ddt, ^1^*J* (^19^F – ^13^C) = 253.7 Hz, ^2^*J* (^19^F – ^13^C) = 16.8 Hz, ^3^*J* (^19^F – ^13^C) = 4.4 Hz), 146.70 (C3 and C5, ddt, ^1^*J* (^19^F – ^13^C) = 243.6 Hz, ^2^*J* (^19^F – ^13^C) = 15.8 Hz, ^3^*J* (^19^F – ^13^C) = 3.4 Hz), 171.35 (C(O)O). ^19^F NMR (376 MHz, DMSO-d_6_, δ): -132.67 – -132.83 (2F, m), -139.06 – -139.23 (2F, m). HRMS for C_10_H_9_F_4_NO_4_S_2_ [(M-H)^-^]: calc. 345.9836, found 345.9836.

Methyl 3-((2,3,5,6-tetrafluoro-4-sulfamoylphenyl)sulfonyl)propanoate **7x** (**7**)

Methyl 3-((2,3,5,6-tetrafluoro-4-sulfamoylphenyl)thio)propanoate (**6x** (**8**)) (0.043 g; 0.12 mmol) ) was dissolved in acetic acid (5 ml) and heated at 75 °C temperature for 4.5 hours. H_2_O_2_ (30%) was added by portions (0.1 ml) every 1.5 hour (overall 0.3 ml) until complete starting material conversion. Afterwards, the solvent was evaporated under reduced pressure and product was recrystallized from H_2_O. Yield: 0.032 g; (70%). Mp: 164 °C. ^1^H NMR (400 MHz, DMSO-d_6_, δ): 2.84 (2H, t, *J =* 7.2 Hz, CH_2_C(O)), 3.59 (3H, s, CH_3_O), 3.85 (2H, t, *J =* 7.2 Hz, SO_2_CH_2_), 8.67 (2H, s, SO_2_NH_2_). ^13^C NMR (100 MHz, DMSO-d_6_, δ): 26.93 (CH_2_C(O)), 51.98 (CH_3_O), 52.51 (SO_2_CH_2_), 120.14 (C1, t, *J (*^19^F – ^13^C) = 15.0 Hz), 127.89 (C4, t, *J (*^19^F – ^13^C) = 15.6 Hz), 143.01 (C2 and C6, dd, ^1^*J (*^19^F – ^13^C) = 259.7 Hz, ^2^*J (*^19^F – ^13^C) = 16.7 Hz), 144.56 (C3 and C5, dd, ^1^*J (*^19^F – ^13^C) = 254.3 Hz, ^2^*J (*^19^F – ^13^C) = 14.9 Hz), 170.12 (C(O)O). ^19^F NMR (376 MHz, DMSO-d_6_, δ): -135,68 -: -135,88 (2F, m), -136,56 -: -136,76 (2F, m). HRMS for C_10_H_9_F_4_NO_6_S_2_ [(M-H)^-^]: calc. 377.9735, found 377.9735.

2-((2,3,5,6-tetrafluoro-4-sulfamoylphenyl)thio)ethyl acetate **8x** (**5**)

2,3,5,6-tetrafluoro-4-((2-hydroxyethyl)thio)benzenesulfonamide (**3ax**) (0.104 g; 0.30 mmol; 1eq.), acetic acid (10 ml) and two drops of H_2_SO_4_ (conc.) were dissolved in toluene (35 ml) and refluxed for 2 hours. The resulting mixture was washed with brine (3x10 ml). The organic phase was dried using anhydrous Na_2_SO_4_ and evaporated under reduced pressure. The product was purified by column chromotography (silica gel, EtOAc/CHCl_3_ (1:1), Rf= 0.76). Yield: 0.078 g; (65%). Mp: 108-110 °C. ^1^H NMR (400 MHz, DMSO-d_6_, δ): 1.90 (3H, s, CH_3_), 3.29 (2H, t, *J =* 5.9 Hz, SCH_2_), 4.16 (2H, t, *J =* 5.3 Hz, CH_2_O), 8.44 (2H, s, SO_2_NH_2_). ^13^C NMR (100 MHz, DMSO-d_6_, δ): 20.27 (CH_3_), 32.56 (SCH_2_, t, *J* (^19^F – ^13^C) = 2.8 Hz), 63.15 (CH_2_O), 117.89 (C4, t, *J* (^19^F – ^13^C) = 20.5 Hz), 122.94 (C1, t, *J* (^19^F – ^13^C) = 15.6 Hz), 142.48 (C3 and C5, ddt, ^1^*J* (^19^F – ^13^C) = 253.7 Hz, ^2^*J* (^19^F – ^13^C) = 17.2 Hz, ^3^*J* (^19^F – ^13^C) = 4.4 Hz), 146.72 (C2 and C6, dd, ^1^*J* (^19^F – ^13^C) = 240.8 Hz, ^2^*J* (^19^F – ^13^C) = 13.9 Hz), 169.95 (OC(O)). ^19^F NMR (376 MHz, DMSO-d_6_, δ): -132.44 – -132.69 (2F, m), -139.10 – -139.30 (2F, m). HRMS for C_10_H_9_F_4_NO_4_S_2_ [(M-H)^-^]: calc. 345.9836, found 345.9838.

2,3,5,6-tetrafluoro-4-((2-hydroxyethyl)sulfinyl)benzenesulfonamide **9x**

2,3,5,6-tetrafluoro-4-((2-hydroxyethyl)thio)benzenesulfonamide (**3ax**) (0.095 g; 0.31 mmol; 1eq.), H_2_O_2_ (0.2 ml; 30%) were dissolved in acetic acid (4 ml) and left to stir at room temperature for 20 hours (after 2 hours additional portion of H_2_O_2_ (0.2 ml; 30%) was added). Afterwards, the solvent was evaporated under reduced pressure and the product was purified by column chromatography (silica gel, EtOAc/CHCl_3_ (1:1), Rf= 0.13).Yield: 0.081 g; (81%). Mp: 160-161 °C. ^1^H NMR (400 MHz, DMSO-d_6_, δ): 3.39 – 3.49 (1H, m, SOCH_2_), 3.59 (1H, dt, ^1^*J* = 13.2 Hz, ^2^*J* = 4.0 Hz, SOCH_2_), 3.84 (2H, q, *J* = 4.8 Hz, CH_2_OH), 5.18 (1H, t, *J* = 5.0 Hz), 8.55 (2H, s, SO_2_NH_2_). ^13^C NMR (100 MHz, DMSO-d_6_, δ): 54.20 (SO_2_CH_2_), 56.90 (CH_2_O), 125.50 (C1, t, *J* = 15.4 Hz), 126.09 (C4, t, *J* = 17.6 Hz), 142.42 (C2 and C6, dd, ^1^*J* (^19^F – ^13^C) = 247.8 Hz, ^2^*J* (^19^F – ^13^C) = 16.4 Hz), 144.34 (C3 and C5, ddt, ^1^*J* (^19^F – ^13^C) = 252.3 Hz, ^2^*J* (^19^F – ^13^C) = 15.7 Hz, ^3^*J* (^19^F – ^13^C) = 5.9 Hz). ^19^F NMR (376 MHz, DMSO-d_6_, δ): -137.61 – -137.80 (2F, m), -139.07 – -139.26 (2F, m). HRMS for C_8_H_7_F_4_NO_4_S_2_ [(M+H)^+^]: calc. 321.9825, found 321.9824.

2-((2,3,5,6-tetrafluoro-4-sulfamoylphenyl)sulfinyl)ethyl acetate **10x** (**4**)

2,3,5,6-tetrafluoro-4-((2-hydroxyethyl)sulfinyl)benzenesulfonamide (**9x**) (0.081 g; 0.22 mmol), acetic acid (2 ml) and 3 drops of H_2_SO_4_ (conc.) were dissolved in toluene (30 ml) and refluxed for 1 hour. The resulting mixture was washed with brine (3x10 ml). The organic phase was dried using anhydrous Na_2_SO_4_ and evaporated under reduced pressure. The product was purified by column chromotography (silica gel, EtOAc/CHCl_3_ (1:1), Rf= 0.36). Yield: 0.023 g; (25%). Mp: 132-133 °C. ^1^H NMR (400 MHz, DMSO-d_6_, δ): 1.96 (3H, s, CH_3_), 3.68 – 3.81 (2H, m, SO_2_CH_2_), 4.36 – 4.49 (2H, m, CH_2_O), 8.58 (2H, s, SO_2_NH_2_). ^13^C NMR (100 MHz, DMSO-d_6_, δ): 20.30 (CH_3_), 52.52 (SO_2_CH_2_), 57.06 (CH_2_O), 125.33 (C1, t, *J* (^19^F – ^13^C) = 17.1 Hz), 125.80 (C4, t, *J* (^19^F – ^13^C) = 16.0 Hz), 142.47 (C2 and C6, dd, ^1^*J* (^19^F – ^13^C) = 251.9 Hz, ^2^*J* (^19^F – ^13^C) = 15.3 Hz), 144.35 (C3 and C5, ^1^*J* (^19^F – ^13^C) = 252.7 Hz, ^2^*J* (^19^F – ^13^C) = 14.9 Hz), 169.92 (OC(O)). ^19^F NMR (376 MHz, DMSO-d_6_, δ): -137.51 – -137.68 (2F, m), -138.94 – -139.10 (2F, m). HRMS for C_10_H_9_F_4_NO_5_S_2_ [(M-H)^-^]: calc. 361.9786, found 361.9794.

2,3,5,6-tetrafluoro-4-(vinylsulfonyl)benzenesulfonamide **11x** (**15**)

2,3,5,6-tetrafluoro-4-((2-hydroxyethyl)sulfonyl)benzensulfonamide (**3ax**) (0.200 g; 0.592 mmol; 1 eq.), thionyl chloride (0.064 ml; 0.85 mmol; 1.5 eq.), Et_3_N (0.006 ml; 0.43 mmol; 0.75 eq.) were dissolved in MeCN (2 ml) and left to stir at room temperature for 52 hours. The resulting mixture was washed with brine (3x10 ml). The organic phase was dried using anhydrous Na_2_SO_4_ and evaporated under reduced pressure. The product was purified by column chromotography (silica gel, EtOAc/CHCl_3_ (1:1), Rf= 0.64). Yield: 0.173 g; (92%). Mp: 174-176 °C. ^1^H NMR (400 MHz, DMSO-d_6_, δ): 6.54 (1H, d, *J =* 17.0 Hz, CHCH_2_), 6.57 (1H, d, *J =* 23.6 Hz, CHCH_2_), 7.36 (1H, dd, ^1^*J =* 16.3 Hz, ^2^*J =* 9.8 Hz, CHCH_2_), 8.63 (2H, s, SO_2_NH_2_). ^13^C NMR (100 MHz, DMSO-d_6_, δ): 121.79 (C1, t, *J* (^19^F – ^13^C) = 14.2 Hz), 128.17 (C4, t, *J* (^19^F – ^13^C) = 15.7 Hz), 133,34 (CH_2_), 138,12 (CH), 143.45 (C3 and C5, dd, ^1^*J* (^19^F – ^13^C) = 259.0 Hz, ^2^*J* (^19^F – ^13^C) = 11.1 Hz) 144.49 (C2 and C6, dd, ^1^*J* (^19^F – ^13^C) = 256.0 Hz, ^2^*J* (^19^F – ^13^C) = 14.8 Hz). ^19^F NMR (376 MHz, DMSO-d_6_, δ): -135.89 – -136.11 (2F, m), -136.43 – -136.72 (2F, m). HRMS for C_8_H_5_F_4_NO_4_S_2_ [(M-H)^-^]: calc. 317.9523, found 317.9524.

2,3,5,6-tetrafluoro-4-((1-hydroxy-2-methylpropan-2-yl)thio)benzenesulfonamide **12x**

2,3,4,5,6-pentafluorobenzensulfonamide (**2ax**) (0.753 g; 3.04 mmol; 1 eq.), 2-mercapto-2-methylpropan-1-ol (0.388 g; 3.66 mmol; 1.2 eq.), Et_3_N (0.510 ml; 3.66 mmol; 1.2 eq.) were dissolved in MeOH (10 ml) and left to stir at room temperature over night. Afterwards the solvent was evaporated under reduced pressure and resultant preciptate was washed with H_2_O. The product was purified by column chromatography (silica gel, EtOAc/CHCl_3_ (1:1), Rf= 0.61). Yield: 0.690 g; (68%). Mp: 146-147 °C. ^1^H NMR (400 MHz, DMSO-d_6_, δ): 1.22 (6H, s, CH_3_), 3.39 (2H, d, *J* = 5.5 Hz, CH_2_OH), 5.12 (1H, s, OH), 8.44 (2H, s, SO_2_NH_2_). ^13^C NMR (100 MHz, DMSO-d_6_, δ): 25.29 (CH_3_), 55.27 (CH_2_OH), 69.96 (SC(CH_3_)_2_), 115.08 (C1, t, *J* (^19^F – ^13^C) = 22.2 Hz), 124.57 (C4, t, *J* (^19^F – ^13^C) = 15.6 Hz), 142.53 (C2 and C6, ddt, ^1^*J* (^19^F – ^13^C) = 254.5 Hz, ^2^*J* (^19^F – ^13^C) = 17.5 Hz, ^3^*J* (^19^F – ^13^C) = 4.1 Hz), 148.32 (C3 and C5, dd, ^1^*J* (^19^F – ^13^C) = 244.0 Hz, ^2^*J* (^19^F – ^13^C) = 14.6 Hz). ^19^F NMR (376 MHz, DMSO-d_6_, δ): -128.45 – -128.60 (2F, m), -138.67 – -138.84 (2F, m). HRMS for C_10_H_11_F_4_NO_3_S_2_ [(M+H)^+^]: calc. 356.0009, found 355.9999.

**Compounds 13x and 14x (9) were synthesized during 12x oxidation.**

2,3,5,6-tetrafluoro-4-((1-hydroxy-2-methylpropan-2-yl)thio)benzenesulfonamide (**12x**) (0.676 g, 2.03 mmol) was dissolved in acetic acid (10 ml) and heated at 75 °C temperature for 2.5 hours. H_2_O_2_ (30%) was added by portions (0.2 ml) every ~0.3H (overall 0.8 ml) until complete starting material conversion. Afterwards, the solvent was evaporated under reduced pressure and product was purified by column chromatography (silica gel, EtOAc/CHCl_3_ (1:1), Rf**_14x (9)_**= 0.66, Rf**_13x_**= 0.55)

2,3,5,6-tetrafluoro-4-((1-hydroxy-2-methylpropan-2-yl)sulfonyl)benzenesulfonamide **13x**

Yield: 0.145 g; (20%). Mp: 210-211 °C. ^1^H NMR (400 MHz, DMSO-d_6_, δ): 1.33 (6H, s, (CH_3_)_2_), 3.69 (2H, d, *J* = 5.3 Hz, CCH_2_), 5.22 (1H, t, *J* = 4.2 Hz, OH), 8.61 (2H, s, SO_2_NH_2_). ^13^C NMR (100 MHz, DMSO-d_6_, δ): 17.35 ((CH_3_)_2_), 63.92 (CH_2_O), 67.74 (SO_2_C(CH_3_)_2_), 120.08 (C1, t, *J* (^19^F – ^13^C) = 14.6 Hz), 127.79 (C4, t, *J* (^19^F – ^13^C) = 15.7 Hz), 142.83 (C2 and C6, dd, ^1^*J* (^19^F – ^13^C) = 252.0 Hz, ^2^*J* (^19^F – ^13^C) = 15.7 Hz), 144.98 (C3 and C5, dd, ^1^*J* (^19^F – ^13^C) = 257.4 Hz, ^2^*J* (^19^F – ^13^C) = 17.7 Hz). ^19^F NMR (376 MHz, DMSO-d_6_, δ): -132.27 – -132.44 (2F, m), -137.02 – -137.18 (2F, m). HRMS for C_10_H_11_F_4_NO_5_S_2_ [(M-H)^+^]: calc. 366.0088, found 366.0088.

2-methyl-2-((2,3,5,6-tetrafluoro-4-sulfamoylphenyl)sulfonyl)propyl acetate **14x** (**9**)

Yield: 0.153 g; (19%). Mp: 177-178 °C. ^1^H NMR (400 MHz, DMSO-d_6_, δ): 1.41 (6H, s, (CH_3_)_2_), 1.87 (3H, s, C(O)CH_3_), 4.29 (2H, s, CCH_2_), 8.64 (2H, s, SO_2_NH_2_). ^13^C NMR (100 MHz, DMSO-d_6_, δ): 17.36 ((CH_3_)_2_), 19.92 (C(O)CH_3_), 65.31 (CH_2_O), 65.76 (SO_2_C(CH_3_)_2_), 118.79 (C1, t, *J* (^19^F – ^13^C) = 14.7 Hz), 128.33 (C4, t, *J* (^19^F – ^13^C) = 15.8 Hz), 143.12 (C2 and C6, dd, ^1^*J* (^19^F – ^13^C) = 258.8 Hz, ^2^*J* (^19^F – ^13^C) = 13.9 Hz), 145.06 (C3 and C5, dd, ^1^*J* (^19^F – ^13^C) = 255.9 Hz, ^2^*J* (^19^F – ^13^C) = 13.9 Hz), 169.41 (OC(O)). ^19^F NMR (376 MHz, DMSO-d_6_, δ): -132.52 – -132.70 (2F, m), -136.40 – -136.57 (2F, m). HRMS for C_12_H_13_F_4_NO_6_S_2_ [(M-H)^-^]: calc. 406.0048, found 406.0052.

N-(2-((2,3,5,6-tetrafluoro-4-sulfamoylphenyl)thio)ethyl)acetamide **15x**

N-(2-((2,3,5,6-tetrafluoro-4-sulfamoylphenyl)thio)ethyl)acetamide (**15x**) was prepared according to known procedure in the literature (2). 2,3,4,5,6-pentafluorbenzensulfonamide (**2ax**) (2.320 g; 9.39 mmol), N-(2-mercaptoethyl)acetamide (1.300 ml; 13.1 mmol; 1.4 eq.), Et_3_N (1.960 ml; 14.1 mmol; 1,5 eq.) were dissolved in MeOH (25 ml) and left to stir at room temperature for 2 hours. Afterwards additional portions of N-(2-mercaptoethyl)acetamide (0.150 ml; 1.51 mmol, 0.16 eq.) and Et_3_N (0.150 ml; 1.08 mmol; 0.11 eq.) were added. The reation mixture was left to stir further for additional 1 hour. After reaction completion the solvent was evaporated under reduced pressure and product was purified by recrystallization from MeOH/H_2_O 1:6 mixture obtaining white crystals. Yield 2.562 (78%). Mp: 169-170 °C. (2), mp: 169-171 °C.

N-(2-((2,3,5,6-tetrafluoro-4-sulfamoylphenyl)sulfonyl)ethyl)acetamide **16x** (**2**)

N-(2-((2,3,5,6-tetrafluoro-4-sulfamoylphenyl)sulfonyl)ethyl)acetamide (**16x** (**2**)) was prepared according to known procedure in literature (4). N-(2-((2,3,5,6-tetrafluoro-4-sulfamoylphenyl)thio)ethyl)acetamide (**15x**) (2.592 g; 7.48 mmol; 1 eq.) was dissolved in acetic acid (70 ml) and heated at 75 °C temperature for 10 hours. H_2_O_2_ (30%) was added by portions (0.1 ml) every 30 minutes (overall 5 ml) until complete starting material conversion. Afterwards, the solvent was evaporated under reduced pressure and the product was purified by recrystallization in MeOH/H_2_O 1:4 mixture obtaining white crystals. Yield: 1.576 g; (56%). Mp: 224-225 °C (4), mp: 224-225 °C).

N'-((4-bromophenyl)sulfonyl)-N,N-dimethylformamidine **18x**

N'-((4-bromophenyl)sulfonyl)-N,N-dimethylformimidamide (**18x**) was prepared according to known procedure in literature (Dudutienė, V. et al. Bioorg. Med. Chem. 21, 2093–2106 (2013). N,N-Dimethylformamide dimetyl acetal (0.303 g; 2.54 mmol; 1.2 eq.) was dissolved in MeCN (7 ml) and added dropwise to a mixture of 4-bromobenzenesulfonamide (**17x**) (0.500 g; 2.12 mmol; 1 eq.) in MeCN (3 ml). Reaction mixture was left to stir at room temperature for 1 hour and afterwards, solvent was evaporated under reduced pressure and resultant precipitate were washed with H_2_O. Yield: 0.562 g; (91%). Mp: 142-143 °C (close to the value in the literature (Dudutienė, V. et al. Bioorg. Med. Chem. 21, 2093–2106 (2013), mp: 141-143 °C).

4-((2-hydroxyethyl)thio)benzenesulfonamide **19x**

4-((2-hydroxyethyl)thio)benzenesulfonamide (**19x**) was prepared according to known procedure in literature (Dudutienė, V. et al. Bioorg. Med. Chem. 21, 2093–2106 (2013). 2-mercaptoethanol (0.161 g; 2.04 mmol; 2 eq.) was added dropwise to a suspension of NaH (0.089 g; 55 % oil dispersion; 2.04 mmol; 2 eq.) in DMF (1 ml). After gas emission was complete, N'-((4-bromophenyl)sulfonyl)-N,N-dimethylformamidine (**18x**) (0.300 g; 1.02 mmol; 1eq.) was added and reaction mixture was heated for 1 hour at 95 °C temperature. Subsequently DMF was removed under reduced pressure, resultant precipitate was dissolve in a mixture of MeOH (1 ml) /NaOH solution (10 %; 1 ml) and refluxed for another hour. MeOH was evaporated under reduced pressure and resultant suspension was diluted with H_2_O, washed with petrol ether and acidified with HCl (10 %). Afterwards reaction mixture was extracted with EtOAc (3x10 ml), collected organic phase was dried using anhydrous Na_2_SO_4_ and evaporated under reduced pressure. The product was purified by column chromotography (silica gel, EtOAc/CHCl_3_ (1:1), Rf= 0.27). Yield 0.149 g; (63%). Mp: 110-111 °C (close to the value in the literature (Dudutienė, V. et al. Bioorg. Med. Chem. 21, 2093–2106 (2013), mp: 111-112 °C).

**Compounds 20x and 21x (6) were synthesized during 19x oxidation.**

4-((2-hydroxyethyl)thio)benzenesulfonamide (**19x**) (0.050 g, 0.21 mmol) was dissolved in acetic acid (2 ml) and heated at 75 °C temperature for 5 hours. H_2_O_2_ (30%) was added by portions (0.1 ml) every hour (overall 0.5 ml) until complete starting material conversion. Afterwards, the solvent was evaporated under reduced pressure and products were purified by column chromatography (silica gel, EtOAc/CHCl_3_ (1:1), Rf**_20x_**= 0.09, Rf**_21x (6_**_)_= 0.34)

4-((2-hydroxyethyl)sulfonyl)benzenesulfonamide **20x**

Yield: 0.032 g; (57%). Mp: 153-154 °C. ^1^H NMR (400 MHz, DMSO-d_6_, δ): 3.54 (2H, t, *J =* 6.2 Hz SO_2_CH_2_), 3.70 (2H, t, *J =* 6.0 Hz, CH_2_O), 7.68 (2H, s, SO_2_NH_2_), 8.04 (2H, d, *J =* 8.7 Hz, ArH), 8.10 (2H, d, *J =* 8.8 Hz, ArH). ^13^C NMR (100 MHz, DMSO-d_6_, δ): 54.98 (SO_2_CH_2_), 57.50 (CH_2_O), 126.47 (C1), 128.65 (C4), 142.97 (C2 and C6), 148.40 (C3 and C5). HRMS for C_8_H_11_NO_5_S_2_ [(M-H)^-^]: calc. 264.0006, found 264.0009.

2-((4-sulfamoylphenyl)sulfonyl)ethyl acetate **21x** (**6**)

Yield: 0.004 g; (6%). Mp; 129-130 °C. ^1^H NMR (400 MHz, DMSO-d_6_, δ): 1.68 (3H, s, COCH_3_), 3.82 (2H, t, *J =* 5.6 Hz, SO_2_CH_2_), 4.27 (2H, t, *J =* 5.6 Hz, CH_2_O), 7.68 (2H, s, SO_2_NH_2_), 8.07 (2H, d, *J =* 8.7 Hz, ArH), 8.12 (2H, d, *J =* 8.7 Hz, ArH). ^13^C NMR (100 MHz, DMSO-d_6_, δ): 20.10 (CH_3_CO), 53.73 (SO_2_CH_2_), 57.49 (CH_2_O), 126.57 (C1), 128.78 (C4), 142.49 (C2 and C6), 148.64 (C3 and C5), 169.59 (OC(O)). HRMS for C_15_H_12_F_4_N_2_O_6_S_2_ [(M-H)^-^]: calc. 306.0112, found 306.0116.

N,N-dimethyl-N'-((2,3,5,6-tetrafluoro-4-((2-hydroxyethyl)sulfonyl)phenyl)sulfonyl)formamidine **22x**

2,3,5,6-tetrafluoro-4-((2-hydroxyethyl)sulfonyl)benzensulfonamide (**4ax** (**1**)) (0.160 g; 0.047 mmol) and N,N-dimethyl-formamide dimethyl acetal (0.076 g; 0.074 mmol; 5 eq.) were dissolved in acetonitrile (10 ml) and left to stir at room temperature for 1 hour. The solvent was evaporated under reduced pressure and the product was purified by column chromotography (silica gel, EtOAc, Rf= 0.65). Yield: 0.155 g; (83%). Mp: 188-189 °C. ^1^H NMR (400 MHz, DMSO-d_6_, δ): 2.99 (3H, s, NCH_3_), 3.23 (3H, s, NCH_3_), 3.71 (2H, t, *J =* 5.3 Hz, SO_2_CH_2_), 3.85 (2H, q, *J =* 5.3 Hz, CH_2_O), 4.98 (1H, t, *J =* 5.2 Hz, OH), 8.30 (1H, s, NCH). ^13^C NMR (100 MHz, DMSO-d_6_, δ): 35.64 (NCH_3_), 41.36 (NCH_3_), 55.07 (SO_2_CH_2_), 59.53 (CH_2_OH), 122.46 (C1, t, *J* (^19^F – ^13^C) = 15.1 Hz), 126.45 (C4, t, *J* (^19^F – ^13^C) = 15.2 Hz), 143.00 (C2 and C6, dd, ^1^*J* (^19^F – ^13^C) = 253.7 Hz, ^2^*J* (^19^F – ^13^C) = 17.8 Hz), 144.23 (C3 and C5, dd, ^1^*J* (^19^F – ^13^C) = 252.2 Hz, ^2^*J* (^19^F – ^13^C) = 16.3 Hz), 160.93 (NCHN). ^19^F NMR (376 MHz, DMSO-d_6_, δ): -136.20 – -136.38 (2F, m), -136.72 – -136.91 (2F, m). HRMS for C_11_H_12_F_4_N_2_O_5_S_2_ [(M+H)^+^]: calc. 393.0197, found 393.0199.

2-((4-(N-((dimethylamino)methylene)sulfamoyl)-2,3,5,6-tetrafluorophenyl)sulfonyl)ethyl phenylcarbamate **23ax**

N,N-dimethyl-N'-((2,3,5,6-tetrafluoro-4-((2-hydroxyethyl)sulfonyl)phenyl)sulfonyl)formamidine (**22x**) (0.068 g; 0.17 mmol; 1 eq.) and phenyl isocyanate (0.028 ml; 0.26 mmol; 1.5 eq.) were dissolved in toluene (10 ml) and refluxed at boiling point for 14 hours. Additional phenyl isocyanate portions (0.057 ml; 0.052 mmol; 3 eq.) were added after 2 hours and 7 hours respectively. The solvent was evaporated under reduced pressure and the product was purified by column chromatography (silica gel, EtOAc/CHCl_3_ (1:1), Rf= 0.43). Yield: 0.072 g; (81%). Mp: 177-178 °C. ^1^H NMR (400 MHz, DMSO-d_6_, δ): 2.96 (3H, s, NCH_3_), 3.22 (3H, s, NCH_3_), 4.04 (2H, t, *J =* 5.3 Hz, SO_2_CH_2_), 4.51 (2H, t, *J =* 5.4 Hz, CH_2_O), 7.00 (1H, t, *J =* 7.3 Hz, CH (C4) of phenyl), 7.27 (2H, t, *J =* 7.8 Hz, CH (C3 and C5) of phenyl), 7.39 (2H, d, *J =* 7.6 Hz, CH (C2 and C6) of phenyl), 8.27 (1H, s, NCHN), 9.62 (1H, s, OC(O)NH). ^13^C NMR (100 MHz, DMSO-d_6_, δ): 35.61 (NCH_3_), 41.34 (NCH_3_), 56.36 (SO_2_CH_2_), 57.38 (CH_2_O), 118.54 (C4 of phenyl), 121.10 (C1, t, *J* (^19^F – ^13^C) = 14.9 Hz), 122.75 (C3 and C5 of phenyl), 126.95 (C4, t, *J* (^19^F – ^13^C) = 15.1 Hz), 128.70 (C2 and C6 of phenyl), 138.54 (C1 of phenyl), 143.29 (C2 and C6, dd, ^1^*J* (^19^F – ^13^C) = 262.1 Hz, ^2^*J* (^19^F – ^13^C) = 22.0 Hz), 144.25 (C5 and C3, dd, ^1^*J* (^19^F – ^13^C) = 247.2 Hz, ^2^*J* (^19^F – ^13^C) = 13.0 Hz), 152.64 (OC(O)NH), 160.89 (NCHN). ^19^F NMR (376 MHz, DMSO-d_6_, δ): -135.99 – -136.16 (2F, m), -136.28 – -136.47 (2F, m). HRMS for C_18_H_17_F_4_N_3_O_6_S_2_ [(M+H)^+^]: calc. 512.0568, found 512.0571.

2-((4-(N-((dimethylamino)methylene)sulfamoyl)-2,3,5,6-tetrafluorophenyl)sulfonyl)ethyl (4-methoxyphenyl)carbamate **23bx**

N,N-dimethyl-N'-((2,3,5,6-tetrafluoro-4-((2-hydroxyethyl)sulfonyl)phenyl)sulfonyl)formamidine (**22x**) (0.050 g; 0.13 mmol; 1 eq.), 4-methoxyphenyl isocyanate (0.025 ml; 0.19 mmol; 1.5 eq.), dibutyltin dilaurate (0.015 ml; 0.026 mmol; 0.2 eq.) were dissolved in acetonitrile (3 ml) and left to stir at room temperature for 5 days. Additional 4-methoxyphenyl isocyanate portion (0.025 ml; 0.19 mmol; 1.5 eq.) was added after 2 days. The solvent was evaporated under reduced pressure and the product was purified by column chromatography (silica gel, EtOAc/CHCl_3_ (1:1), Rf= 0.41). Yield: 0.008 g; (12%). Mp: 167-168 °C. ^1^H NMR (400 MHz, DMSO-d_6_, δ): 2.96 (3H, s, NCH_3_), 3.21 (3H, s, NCH_3_), 3.70 (3H, s, OCH_3_), 4.03 (2H, t, *J* = 5.4 Hz, SO_2_CH_2_), 4.49 (2H, t, *J* = 5.3 Hz, CH_2_O), 6.85 (2H, d, *J* = 8.8 Hz, CH (C2 and C6) of phenyl), 7.28 (2H, d, *J* = 7.2 Hz, CH (C3 and C6) of phenyl), 8.27 (1H, s, NCHN), 9.41 (1H, s, OC(O)NH). ^13^C NMR (100 MHz, DMSO-d_6_, δ): 35.58 (NCH_3_), 41.34 (NCH_3_), 55.14 (OCH_3_), 56.40 (SO_2_CH_2_), 57.31 (CH_2_O), 113.91 (C3 and C5 of phenyl), 120.25 (C2 and C6 of phenyl), 121.13 (C1, t, *J* (^19^F – ^13^C) = 14.7 Hz), 126.91 (C4), 128.63 (C1 of phenyl), 143.34 (C2 and C6, dd, ^1^*J* (^19^F – ^13^C) = 250.0 Hz, ^2^*J* (^19^F – ^13^C) = 11.2 Hz), 144.16 (C3 and C5, dd, ^1^*J* (^19^F – ^13^C) = 242.7 Hz, ^2^*J* (^19^F – ^13^C) = 10.3 Hz), 152.79 (OC(O)NH), 155.03 (C4 of phenyl), 160.90 (NCHN). ^19^F NMR (376 MHz, DMSO-d_6_, δ): -135.95 – -136.16 (2F, m), -136.30 – -136.55 (2F,m). HRMS for C_19_H_19_F_4_N_3_O_7_S_2_ [(M+Na)^+^]: calc. 564.0493, found 564.0501.

2-((2,3,5,6-tetrafluoro-4-sulfamoylphenyl)sulfonyl)ethyl phenylcarbamate **24ax** (**18**)

2-((4-(N-((dimethylamino)methylene)sulfamoyl)-2,3,5,6-tetrafluorophenyl)sulfonyl)ethyl phenylcarbamate (**23ax**) (0.030 g; 0.059 mmol; 1eq.) and three drops of HCl (conc.) were dissolved in toluene (3 ml) and refluxed at boiling point for 18 hours. After 5 hours additional 10 drops of HCl (conc.) were added. The solvent was evaporated under reduced pressure and the product was purified by column chromatography (silica gel, EtOAc/CHCl_3_ (1:1), Rf= 0.59). Yield: 0.007 g; (26%). Mp: 204-205 °C. ^1^H NMR (400 MHz, DMSO-d_6_, δ): 4.06 (2H, t, *J =* 5.4 Hz, SO_2_CH_2_), 4.52 (2H, t, *J =* 5.5 Hz, CH_2_O), 7.00 (1H, t, *J =* 7.3 Hz, CH (C4) of phenyl), 7.26 (2H, t, *J =* 7.9 Hz, (C3 and C5) of phenyl), 7.41 (2H, d, *J =* 7.9 Hz, CH (C2 and C6) of phenyl), 8.60 (2H, s, SO_2_NH_2_), 9.65 (1H, s, OC(O)NH). ^13^C NMR (100 MHz, DMSO-d_6_, δ): 56.39 (SO_2_CH_2_), 57.26 (CH_2_O), 118.51 (C4 of phenyl), 121.18 (C1, t, *J* (^19^F – ^13^C) = 13.1 Hz), 122.72 (C3 and C5 of phenyl), 127.86 (C4, t, *J* (^19^F – ^13^C) = 15.0 Hz), 128.71 (C2 and C6 of phenyl), 138.57 (C1 of phenyl), 143.12 (C2 and C6, dd, ^1^*J* (^19^F – ^13^C) = 257.4 Hz, ^2^*J* (^19^F – ^13^C) = 17.7 Hz), 144.26 (C3 and C5, dd, ^1^*J* (^19^F – ^13^C) = 259.2 Hz, ^2^*J* (^19^F – ^13^C) = 19.5 Hz), 152.65 (OC(O)NH). ^19^F NMR (376 MHz, DMSO-d_6_, δ): -136.18 – -136.49 (4F, m). HRMS for C_15_H_12_F_4_N_2_O_6_S_2_ [(M-H)^-^]: calc. 455.0000, found 455.0001.

2-((2,3,5,6-tetrafluoro-4-sulfamoylphenyl)sulfonyl)ethyl (4-methoxyphenyl)carbamate **24bx** (**19**)

2-((4-(N-((dimethylamino)methylene)sulfamoyl)-2,3,5,6-tetrafluorophenyl)sulfonyl)ethyl (4-methoxyphenyl)carbamate (**23bx**) (0.006 g; 0.011 mmol; 1 eq.) and three drops of HCl (conc.) were dissolved in MeOH (3 ml) and refluxed for 31 hours. The solvent was evaporated under reduced pressure and the product was purified by column chromatography (silica gel, EtOAc/CHCl_3_, (1:1), Rf= 0.57). Yield: 0.002 g; (37%). Mp: 188-189 °C. ^1^H NMR (400 MHz, DMSO-d_6_, δ): 3.70 (3H, s, OCH_3_), 4.05 (2H, t, *J* = 5.4 Hz, SO_2_CH_2_), 4.49 (2H, t, *J* = 5.4 Hz, CH_2_O), 6.85 (2H, d, *J* = 8.9 Hz, phenyl), 7.30 (2H, d, *J* = 7.5 Hz, phenyl), 8.61 (2H, s, SO_2_NH_2_), 9.45 (1H, s, OC(O)NH). ^13^C NMR (100 MHz, DMSO-d_6_, δ): 55.16 (OCH_3_), 56.44 (SO_2_CH_2_), 57.23 (CH_2_O), 113.94 (C3 and C5 of phenyl), 120.14 (C2 and C6 of phenyl), 121.23 (C1), 127.85 (C4, t, *J* (^19^F – ^13^C) = 12.2 Hz), 143.06 (C2 and C6, d, *J* (^19^F – ^13^C) = 255.9 Hz), 144.26 (C3 and C5, dd, ^1^*J* (^19^F – ^13^C) = 259.6 Hz, ^2^*J* (^19^F – ^13^C) = 19.2 Hz), 152.82 (OC(O)NH), 155.01 (C4 of phenyl). ^19^F NMR (376 MHz, DMSO-d_6_, δ): -136.22 – -136.49 (4F, m). HRMS for C_16_H_14_F_4_N_2_O_7_S_2_ [(M-H)^-^]: calc. 485.0106, found 485.0105.

3-(cyclooctylamino)-2,5,6-trifluoro-4-((2-hydroxyethyl)sulfonyl)benzenesulfonamide **25ax** (**11**)

3-(cyclooctylamino)-2,5,6-trifluoro-4-((2-hydroxyethyl)sulfonyl)benzenesulfonamide (**25ax** (**11**) was prepared according to known procedure in literature (Dudutienė, V. et al. J. Med. Chem. 57 (2014) 9435–9446). 2,3,5,6-tetrafluoro-4-((2-hydroxyethyl)sulfonyl)benzensulfonamide (**4ax** (**1**)) (0.549 g; 1.63 mmol; 1 eq.) and cyclooctylamine (0.446 ml; 3.26 mmol; 2 eq.) were dissolved in DMSO (2 ml) and left to stir at room temperature overnight. After full starting material conversion reaction mixture was washed with brine (10 ml) and extracted with EtOAc (3x15 ml). The organic phase was dried using anhydrous Na_2_SO_4_ and evaporated under reduced pressure. The product was purified by column chromotography (silica gel, EtOAc/CHCl_3_ (1:1), Rf= 0.52). Yield 0.380 g; (52%). Mp: 90-91 °C (close to the value in the literature (Dudutienė, V. et al. J. Med. Chem. 57 (2014) 9435–9446), mp: 89-90 °C).

3-(cyclododecylamino)-2,5,6-trifluoro-4-((2-hydroxyethyl)sulfonyl)benzenesulfonamide **25bx**

3-(cyclododecylamino)-2,5,6-trifluoro-4-((2-hydroxyethyl)sulfonyl)benzenesulfonamide (**25bx**) was prepared according to known procedure in literature (Dudutienė, V. et al. ChemMedChem 2015, 10, 662 – 687). 2,3,5,6-tetrafluoro-4-((2-hydroxyethyl)sulfonyl)benzensulfonamide (**4ax** (**1**)) (0.300 g 0.89 mmol; 1 eq.) and cyclododecylamine (0.326 g; 1.78 mmol; 2 eq.) were dissolved in DMSO (2 ml) and left to stir at room temperature for 2 hours. After full starting material conversion reaction mixture was washed with brine (10 ml) and extracted with EtOAc (3x15 ml). The organic phase was dried using anhydrous Na_2_SO_4_ and evaporated under reduced pressure. The product was purified by column chromotography (silica gel, EtOAc/CHCl_3_ (1:1), Rf= 0.60). Yield: 0.278 g; (62%). Mp: 143-145 °C (close to the value in the literature (Dudutienė, V. et al. ChemMedChem 2015, 10, 662 – 687), mp: 143-144 °C).

N'-((3-(cyclooctylamino)-2,5,6-trifluoro-4-((2-hydroxyethyl)sulfonyl)phenyl)sulfonyl)-N,N-dimethylformamidine **26x**

N,N-dimethyl-N'-((2,3,5,6-tetrafluoro-4-((2-hydroxyethyl)sulfonyl)phenyl)sulfonyl)formamidine (**22x**) (0.100 g; 0.25 mmol; 1 eq.) and cyclooctylamine (0.070 ml; 0.51 mmol; 2 eq.) were dissolved in DMSO (1 ml) and left to stir at room temperature for 1.5 hours. The reaction mixture was washed with brine (5 ml) and extracted using EtOAc (3x10 ml). The collected organic phase was dried using anhydrous Na_2_SO_4_ and evaporated under reduced pressure. The product was purified by column chromatography (silica gel, EtOAc, Rf= 0.64) and yellow oil was obtained. Yield: 0.062 g; (49%). ^1^H NMR (400 MHz, DMSO-d_6_, δ): 1.40 – 1.70 (12H, m, cyclooctane), 1.76 – 1.87 (2H, m, cyclooctane), 2.97 (3H, s, NCH_3_), 3.21 (3H, s, NCH_3_), 3.64 (2H, t, *J =* 5.4 Hz, SO_2_CH_2_), 3.73 (1H, br.s, NHCH(CH_2_)_2_), 3.81 (2H, q, *J =* 5,4 Hz, CH_2_O), 4.98 (1H, t, *J =* 5.2 Hz, OH), 6.60 (1H, d, *J =* 8.2 Hz, NHCH(CH_2_)_2_), 8.28 (1H, s, NCHN). ^13^C NMR (100 MHz, DMSO-d_6_, δ): 22.89 (cyclooctane), 25.11 (cyclooctane), 26.65 (cyclooctane), 32.27 (cyclooctane), 35.48 (NCH_3_), 41.20 (NCH_3_), 55.02 (CH_2_O), 55.31 (CH of cyclooctane, d, *J* (^19^F – ^13^C) = 11.1 Hz), 59.52 (SO_2_CH_2_, d, *J* (^19^F – ^13^C) = 2.5 Hz), 116.70 (C1, dd, ^1^*J* (^19^F – ^13^C) = 12.8 Hz, ^2^*J* (^19^F – ^13^C) = 5.5 Hz), 126.05 (C4, dd, ^1^*J* (^19^F – ^13^C) = 18.2 Hz, ^2^*J* (^19^F – ^13^C) = 13.9 Hz), 134.49 (C3, d, *J* (^19^F – ^13^C) = 16.1 Hz), 136.97 (C6, d, *J* (^19^F – ^13^C) = 246.7 Hz), 144.26 (C2, d, ^1^*J* (^19^F – ^13^C) = 252.5 Hz), 145.62 (C5, dd, ^1^*J* (^19^F – ^13^C) = 246.0 Hz, ^1^*J* (^19^F – ^13^C) = 12.4 Hz), 160.75 (NCHN). ^19^F NMR (376 MHz, DMSO-d_6_, δ): -124.67 (1F, br.s), -134.26 (1F, dd, *J*^1^= 27.3 Hz, *J*^2^= 12.2 Hz), -150.64 (1F, dd, *J*^1^= 27.3 Hz, *J*^2^= 6.7 Hz). HRMS for C_19_H_28_F_3_N_3_O_5_S_2_ [(M+H)^+^]: calc. 500.1495, found 500.1493.

2-((2-(cyclooctylamino)-3,5,6-trifluoro-4-sulfamoylphenyl)sulfonyl)ethyl acetate **27ax** (**20**)

3-(cyclooctylamino)-2,5,6-trifluoro-4-((2-hydroxyethyl)sulfonyl)benzenesulfonamide (**25ax** (**11**) (0.040 g; 0.090 mmol), acetic acid (1.2 ml) and two drops of H_2_SO_4_ (conc.) were dissolved in toluene (10 ml) and refluxed for 1 hour. The resulting mixture was washed with brine (3x10 ml). The organic phase was dried using anhydrous Na_2_SO_4_ and evaporated under reduced pressure. The product was purified by column chromotography (silica gel, EtOAc/CHCl_3_ (1:1), Rf= 0.78). Yield: 0.0203 g; (46%). Mp: 96 °C. ^1^H NMR (400 MHz, CDCl_3_, δ): 1.45 – 1.73 (12H, m, cyclooctane), 1.83 – 1.91 (2H, m, cyclooctane), 1.93 (3H, s, CH_3_), 3.66 (2H, t, *J =* 5.7 Hz, SO_2_CH_2_), 3.88 (1H, s, cyclooctane), 4.48 (2H, t, *J =* 5.7 Hz, CH_2_O), 5.65 (2H, s, SO_2_NH_2_), 6.84 (2H, d, *J =* 6.7 Hz, NH). ^13^C NMR (100 MHz, CDCl_3_, δ): 20.46 (CH_3_), 23.38 (cyclooctane), 25.54 (cyclooctane), 27.31 (cyclooctane), 33.04 (cyclooctane), 56.24 (cyclooctane, d, *J* (^19^F-^13^C) = 11.7 Hz), 56.43 (SO_2_CH_2_, d, *J (*^19^F-^13^C) = 4.0 Hz), 57.39 (CH_2_O), 115.10 (C1, dd, ^1^*J (*^19^F-^13^C) = 12.7 Hz, ^2^*J (*^19^F-^13^C) = 5.9 Hz), 126.63 (C4, dd, ^1^*J (*^19^F-^13^C) = 17.1 Hz, ^2^*J (*^19^F-^13^C) = 13.3 Hz), 135.85 (C3, d, *J (*^19^F-^13^C) = 13.6 Hz), 136.57 (C6, ddd, ^1^*J (*^19^F-^13^C) = 248.33 Hz, ^2^*J (*^19^F-^13^C) = 17.8 Hz, ^3^*J (*^19^F-^13^C) = 3.9 Hz), 144.36 (C2, d, ^1^*J (*^19^F-^13^C) = 252.7 Hz), 146.28 (C5, dd, ^1^*J (*^19^F-^13^C) = 252.8 Hz, ^2^*J (*^19^F-^13^C) = 15.8 Hz, ^3^*J (*^19^F-^13^C) = 4.5 Hz), 170.29 (OC(O)). ^19^F NMR (376 MHz, CDCl_3_, δ): -125.63 (1F, dd, ^1^*J =* 12.4 Hz, ^2^*J* = 8.8 Hz), -132.86 (1F, dd, ^1^*J =* 25.7 Hz, ^2^*J* = 12.5 Hz), -151.63 (1F, dd, ^1^*J =* 25.8 Hz, ^2^*J* = 8.8 Hz). HRMS for C_18_H_25_F_3_N_2_O_6_S_2_ [(M+H)^+^]: calc. 487.1179, found 487.1179.

2-((2-(cyclododecylamino)-3,5,6-trifluoro-4-sulfamoylphenyl)sulfonyl)ethyl acetate **27bx** (**23**)

3-(cyclododecylamino)-2,5,6-trifluoro-4-((2-hydroxyethyl)sulfonyl)benzenesulfonamide (**25bx**) (0.102 g; 0.204 mmol), acetic acid (4 ml) and three drops of H_2_SO_4_ (conc.) were dissolved in toluene (25 ml) and refluxed for 4 hours. The resulting mixture was washed with brine (3x10 ml). The organic phase was dried using anhydrous Na_2_SO_4_ and evaporated under reduced pressure. The product was purified by column chromotography (silica gel, EtOAc/CHCl_3_ (1:1), Rf= 0.86). Yield: 0.052 g; (47%). Mp: 138-140 °C. ^1^H NMR (400 MHz, DMSO-d_6_, δ): 1.21 – 1.46 (20H, m, cyclododecane), 1.55 – 1.65 (2H, m, cyclododecane), 1.81 (3H, s, OC(O)CH_3_), 3.78 (1H, s, cyclododecane), 3.92 (2H, t, *J =* 4.9 Hz, SO_2_CH_2_), 4.36 (2H, t, *J =* 4.9 Hz, CH_2_O), 6.52 (1H, d, *J =* 8.6 Hz, NH), 8.38 (2H, s, SO_2_NH_2_). ^13^C NMR (100 MHz, DMSO-d_6_, δ): 20.04 (CH_3_), 20.45 (cyclododecane), 22.60 (cyclododecane), 22.71 (cyclododecane), 23.73 (cyclododecane), 23.89 (cyclododecane), 30.08 (cyclododecane), 52.84 (CH of cyclododecane, d, *J =* 11.6 Hz), 55.66 (SO_2_CH_2_, d, *J =* 2.3 Hz), 57.65 (CH_2_O), 115.34 (C1, dd, ^1^*J* (^19^F – ^13^C) = 13.2 Hz, ^2^*J* (^19^F – ^13^C) = 5.1 Hz), 127.71 (C4, dd, ^1^*J* (^19^F – ^13^C) = 18.1 Hz, ^2^*J* (^19^F – ^13^C) = 13.9 Hz), 135.13 (C3, d, ^1^*J* (^19^F – ^13^C) = 15.3 Hz), 136.67 (C6, d, ^1^*J* (^19^F – ^13^C) = 253.6 Hz), 143.95 (C2, d, ^1^*J* (^19^F – ^13^C) = 252.9 Hz), 145.58 (C5, dd, ^1^*J* (^19^F – ^13^C) = 246.4 Hz, ^2^*J* (^19^F – ^13^C) = 13.7 Hz), 169.54 (OC(O)). ^19^F NMR (376 MHz, DMSO-d_6_, δ): -124.88 (1F, s), -134.22 (1F, dd, ^1^*J =* 26.8 Hz, ^2^*J =* 12.5 Hz), -150.76 (1F, dd, ^1^*J =* 26.8 Hz, ^2^*J =* 6.7 Hz). HRMS for C_22_H_33_F_3_N_2_O_6_S_2_ [(M+H)^+^]: calc. 543.1805, found 543.1803.

2-((2-(cyclooctylamino)-3,5,6-trifluoro-4-sulfamoylphenyl)sulfonyl)ethyl propionate **28ax** (**21**)

3-(cyclooctylamino)-2,5,6-trifluoro-4-((2-hydroxyethyl)sulfonyl)benzenesulfonamide (**25ax** (**11**)) (0.100 g; 0.225 mmol), propionic acid (3 ml) and three drops of H_2_SO_4_ (conc.) were dissolved in toluene (25 ml) and refluxed for 3 hours. The resulting mixture was washed with brine (3x10 ml). The organic phase was dried using anhydrous Na_2_SO_4_ and evaporated under reduced pressure. The product was purified by column chromotography (silica gel, EtOAc/CHCl_3_ (1:1), Rf= 0.89) and yellow oil was obtained. Yield: 0.0313 g; (28%). ^1^H NMR (400 MHz, CDCl_3_, δ): 1.06 (3H, t, *J* = 7.6 Hz, CH_2_CH_3_), 1.45-1.74 (12H, m, cyclooctane), 1.83-1.92 (2H, m, cyclooctane), 2.17 (2H, q, *J* = 7.6 Hz, CH_2_CH_3_), 3.66 (2H, t, *J* = 5.8 Hz, SO_2_CH_2_), 3.88 (1H, s, CH of cyclooctane), 4.49 (2H, t, *J* = 5.8 Hz, CH_2_O), 5.55 (2H, s, SO_2_NH_2_), 6,85 (1H, d, *J =* 8.0 Hz, NH). ^13^C NMR (100 MHz, CDCl_3_, δ): ): 8.88 (CH_2_CH_3_), 23.40 (cyclooctane), 25.56 (cyclooctane), 27.19 (CH_2_CH_3_), 27.31 (cyclooctane), 33.06 (cyclooctane), 56.25 (CH of cyclooctane, d, *J (*^19^F-^13^C) = 11.7 Hz), 56.58 (SO_2_CH_2_, d, *J (*^19^F-^13^C) = 3.9 Hz), 57.28 (CH_2_O), 115.18 (C1, dd, ^1^*J* (^19^F-^13^C) = 13.3 Hz, ^2^*J* (^19^F-^13^C) = 6.5 Hz), 126.63 (C4, dd, ^1^*J* (^19^F-^13^C) = 16.8 Hz, ^2^*J* (^19^F-^13^C) = 13.4 Hz), 135.89 (C3, d, *J (*^19^F-^13^C) = 12.9 Hz), 136.59 (C6, ddd, ^1^*J (*^19^F-^13^C) = 248.3 Hz, ^2^*J (*^19^F-^13^C) = 17.4 Hz, ^3^*J (*^19^F-^13^C) = 4.3 Hz), 144.39 (C2, d, ^1^*J (*^19^F-^13^C) = 253.3 Hz, ^2^*J (*^19^F-^13^C) = 3.7 Hz), 146.30 (C5, ddd, ^1^*J (*^19^F-^13^C) = 253.2 Hz, ^2^*J (*^19^F-^13^C) = 16.5 Hz, ^3^*J (*^19^F-^13^C) = 5.0 Hz), 173.73 (OC(O)). ^19^F NMR (376 MHz, CDCl_3_, δ): -125.61 (1F, dd, ^1^*J =* 12.4 Hz, ^2^*J =* 8.9 Hz), -132.79 (1F, dd, ^1^*J =* 25.8 Hz, ^2^*J =* 12.5 Hz), -151.60 (1F, dd, ^1^*J =* 25.8 Hz, ^2^*J =* 8.8 Hz). HRMS for C_19_H_27_F_3_N_2_O_6_S_2_ [(M+H)^+^]: calc. 501.1335, found 501.1339.

2-((2-(cyclododecylamino)-3,5,6-trifluoro-4-sulfamoylphenyl)sulfonyl)ethyl propionate **28bx** (**24**)

3-(cyclododecylamino)-2,5,6-trifluoro-4-((2-hydroxyethyl)sulfonyl)benzenesulfonamide (**25bx**) (0.0377 g; 0.075 mmol), propionic acid (2 ml) and three drops of H_2_SO_4_ (conc.) were dissolved in toluene (10 ml) and refluxed for 1 hour. The resulting mixture was washed with brine (2x5 ml). The organic phase was dried using anhydrous Na_2_SO_4_ and evaporated under reduced pressure. The product was purified by column chromotography (silica gel, CHCl_3_, Rf= 0.20). Yield: 0.0248 g; (59%). Mp: 110-112 °C. ^1^H NMR (400 MHz, DMSO-d_6_, δ): 0.91 (3H, t, *J =* 7.5 Hz, CH_2_CH_3_), 1.23 – 1.44 (20H, m, cyclododecane), 1.54 – 1.64 (2H, m, cyclododecane), 2.05 (2H, q, *J =* 7.6 Hz, CH_2_CH_3_), 3.78 (1H, s, cyclododecane), 3.94 (2H, t, *J =* 5.2 Hz, SO_2_CH_2_), 4.37 (2H, t, *J =* 5.0 Hz, CH_2_O), 6.52 (1H, d, *J =* 8.2 Hz, NH), 8.40 (2H, s, SO_2_NH_2_). ^13^C NMR (100 MHz, DMSO-d_6_, δ): 8.56 (CH_2_CH_3_), 20.44 (cyclododecane), 22.58 (cyclododecane), 22.70 (cyclododecane), 23.71 (cyclododecane), 23.87 (cyclododecane), 26.36 (CH_2_CH_3_), 30.08 (cyclododecane), 52.82 (CH of cyclododecane, d, *J =* 11.6 Hz), 55.76 (SO_2_CH_2_), 57.59 (CH_2_O), 115.32 (C1, t, *J* (^19^F – ^13^C) = 7.0 Hz), 127.67 (C4, t, *J* (^19^F – ^13^C) = 14.2 Hz), 135.11 (dd, ^1^*J* (^19^F – ^13^C) = 13.9 Hz, ^2^*J* (^19^F – ^13^C) = 2.4 Hz), 136.65 (C6, d, ^1^*J* (^19^F – ^13^C) = 252.8 Hz), 143.98 (C2, d, ^1^*J* (^19^F – ^13^C) 253.0 Hz, ), 145.56 (C5, dd, ^1^*J* (^19^F – ^13^C) = 249.8 Hz, ^2^*J* (^19^F – ^13^C) = 15.4 Hz), 172.82 (OC(O)). ^19^F NMR (376 MHz, DMSO-d_6_, δ): -124.86 (1F, s), -134.21 (1F, dd, ^1^*J =* 26.9 Hz, ^2^*J =* 12.4 Hz), -150.79 (1F, dd, ^1^*J =* 26.9 Hz, ^2^*J =* 6.6 Hz). HRMS for C_23_H_35_F_3_N_2_O_6_S_2_ [(M+H)^+^]: calc. 557.1961, found 557.1966.

2-((2-(cyclooctylamino)-4-(N-((dimethylamino)methylene)sulfamoyl)-3,5,6-trifluorophenyl)sulfonyl)ethyl phenylcarbamate **29x**

N'-((3-(cyclooctylamino)-2,5,6-trifluoro-4-((2-hydroxyethyl)sulfonyl)phenyl)sulfonyl)-N,N-dimethylformamidine (**26x**) (0.038 g; 0.08 mmol; 1 eq.) and phenyl isocyanate (0.025 ml; 0.23 mmol; 3 eq.) were dissolved in toluene (7 ml) and refluxed at boiling point for 28 hours. Additional phenyl isocyanate portions (0.025 ml; 0.023 mmol; 3 eq.) were added after 10 hours and 24 hours respectively. The solvent was evaporated under reduced pressure and the product was purified by column chromatography (silica gel, EtOAc/CHCl_3_ (1:1), Rf= 0.65). Yield: 0.018 g; (38%). Mp: 177-178 °C. ^1^H NMR (400 MHz, DMSO-d_6_, δ): 1.33 – 1.63 (12H, m, cyclooctane), 1.71 – 1.82 (2H, m, cyclooctane), 2.95 (3H, s, NCH_3_), 3.20 (3H, s, NCH_3_), 3.69 (1H, br.s, CH of cyclooctane), 3.95 (2H, t, *J* = 5.0 Hz, SO_2_CH_2_), 4.46 (2H, t, *J* = 5.2 Hz, CH_2_O), 6.58 (1H, d, *J* = 8.2 Hz, NHCH(CH_2_)_2_), 6.99 (1H, t, *J* = 7.3 Hz, CH (C4) of phenyl), 7.26 (2H, t, *J* = 7.8 Hz, CH (C3 and C5) of phenyl), 7.42 (2H, d, *J* = 6.6 Hz, CH (C2 and C6) of phenyl), 8.27 (1H, s, NCHN), 9.66 (1H, s, C(O)NH). ^13^C NMR (100 MHz, DMSO-d_6_, δ): 22.83 (cyclooctane), 25.03 (cyclooctane), 26.54 (cyclooctane), 32.31 (cyclooctane), 35.46 (NCH_3_), 41.20 (NCH_3_), 55.27 (CH of cyclooctane, d, *J* = 11.3 Hz), 56.14 (SO_2_CH_2_, d, *J* = 2.4 Hz), 57.50 (CH_2_O), 115.03 (C1, dd, ^1^*J* (^19^F – ^13^C) = 13.2 Hz, ^2^*J* (^19^F – ^13^C) =5.9 Hz), 118.48 (C4 of phenyl), 122.64 (C3 and C5 of phenyl), 126.57 (C4, dd, ^1^*J* (^19^F – ^13^C) = 18.2 Hz, ^2^*J* (^19^F – ^13^C) = 14.1 Hz), 128.62 (C2 and C6 of phenyl), 134.55 (C3, d, *J* (^19^F – ^13^C) = 14.5 Hz), 137.09 (C6, ddd, ^1^*J* (^19^F – ^13^C) = 241.5 Hz, ^2^*J* (^19^F – ^13^C) =17.6 Hz, ^3^*J* (^19^F – ^13^C) = 5.0 Hz), 138.68 (C1 of phenyl), 144.29 (C2, dd, ^1^*J* (^19^F – ^13^C) = 253.2 Hz, ^2^*J* (^19^F – ^13^C) = 21.3 Hz), 145.66 (C5, ddd, ^1^*J* (^19^F – ^13^C) = 250.5 Hz, ^2^*J* (^19^F – ^13^C) = 17.2 Hz, ^3^*J* (^19^F – ^13^C) = 4.0 Hz), 152.75 (OC(O)NH), 160.76 (NCHN). ^19^F NMR (376 MHz, DMSO-d_6_, δ): -124.61 (1F, s), -134.25 (1F, dd, ^1^*J* = 27.4 Hz, ^2^*J* = 12.3 Hz), 150.25 (1F, dd, ^1^*J* = 27.7 Hz, ^2^*J* = 6.7 Hz). HRMS for C_26_H_33_F_3_N_4_O_6_S_2_ [(M-H)^-^]: calc. 617.1721, found 617.1723.

2-((2-(cyclooctylamino)-3,5,6-trifluoro-4-sulfamoylphenyl)sulfonyl)ethyl phenylcarbamate **30x** (**22**)

2-((2-(cyclooctylamino)-4-(N-((dimethylamino)methylene)sulfamoyl)-3,5,6-trifluorophenyl)sulfonyl)ethyl phenylcarbamate (**29x**) (0.015 g; 0.024; 1 eq.) and three drops of HCl (conc.) were dissolved in MeOH (3 ml) and refluxed at boiling point for 29 hours. The solvent was evaporated under reduced pressure and the product was purified by column chromatography (silica gel, EtOAc/CHCl_3_ (1:1), Rf= 0.88). Yield: 0.004 g; (29%). Mp: 174-175 °C. ^1^H NMR (400 MHz, DMSO-d_6_, δ): 1.36 – 1.63 (12H, m, cyclooctane), 1.75 – 1.85 (2H, m, cyclooctane), 3.72 (1H, br.s, CH of cyclooctane), 3.97 (2H, t, *J* = 5.3 Hz, SO_2_CH_2_), 4.46 (2H, t, *J* = 5.4 Hz, CH_2_O), 6.62 (1H, d, *J* = 8.3 Hz, NHCH(CH_2_)_2_), 6.99 (1H, t, *J* = 7.3 Hz, C4 of phenyl), 7.26 (2H, t, *J* = 7.8 Hz, C3 and C5 of phenyl), 7.43 (2H, d, *J* = 8.0 Hz, C2 and C6 of phenyl), 8.32 (2H, s, SO_2_NH_2_), 9.67 (1H, s, OC(O)NH). ^13^C NMR (100 MHz, DMSO-d_6_, δ): 22.74 (cyclooctane), 24.89 (cyclooctane), 26.66 (cyclooctane), 32.15 (cyclooctane), 55.39 (CH of cyclooctane, d, *J* = 11.2 Hz), 56.14 (SO_2_CH_2_, d, *J* = 2.8 Hz), 57.45 (CH_2_O), 114.93 (C1, d, *J* (^19^F – ^13^C) = 12.6 Hz), 118.45 (C4 of phenyl), 122.64 (C3 and C5 of phenyl), 127.61 (C4, dd, ^1^*J* (^19^F – ^13^C) = 18.6 Hz, ^2^*J* (^19^F – ^13^C) = 14.2 Hz), 128.65 (C2 and C6 of phenyl), 134.66 (C3, dd, ^1^*J* (^19^F – ^13^C) = 14.2 Hz, ^2^*J* (^19^F – ^13^C) = 2.7 Hz), 138.68 (C1 of phenyl), 144.10 (C2, d, *J* (^19^F – ^13^C) = 18.2 Hz), 145.59 (C5, ddd, ^1^*J* (^19^F – ^13^C) = 250.0 Hz, ^2^*J* (^19^F – ^13^C) = 19.3 Hz, ^3^*J* (^19^F – ^13^C) = 3.0 Hz), 152.75 (OC(O)NH). ^19^F NMR (376 MHz, DMSO-d_6_, δ): -124.89 (1F, s), -134.36 (1F, dd, ^1^*J* = 27.0 Hz, ^2^*J* = 12.2 Hz), -150.47 (1F, dd, ^1^*J* = 27.0 Hz, ^2^*J* = 6.4 Hz). HRMS for C_23_H_28_F_3_N_3_O_6_S_2_ [(M+H)^+^]: calc. 564.1444, found 564.1444.

N-(2-((2-(cyclooctylamino)-3,5,6-trifluoro-4-sulfamoylphenyl)sulfonyl)ethyl)acetamide **31x** (**10**)

N-(2-((2-(cyclooctylamino)-3,5,6-trifluoro-4-sulfamoylphenyl)sulfonyl)ethyl)acetamide (**31x** (**10**)) was prepared according to known procedure in literature (4). N-(2-((2,3,5,6-tetrafluoro-4-sulfamoylphenyl)sulfonyl)ethyl)acetamide (**16x** (**2**) (1.39 g; 3.67 mmol; 1 eq.) and cyclooctylamine (1.01 ml; 7.35 mmol; 2 eq.) were dissolved in DMSO (3 ml) and left stirring at room temperature overnight. After full starting material conversion reaction mixture was washed with brine (10 ml) and extracted with EtOAc (3x15 ml). The organic phase was dried using anhydrous Na_2_SO_4_ and evaporated under reduced pressure. The product was purified by column chromotography (silica gel, EtOAc Rf= 0.53). Yield 0.900 g; (50%). Mp: 161-162 °C (close to the value in the literature (4), mp: 162-163 °C).

**Compound purity by HPLC**

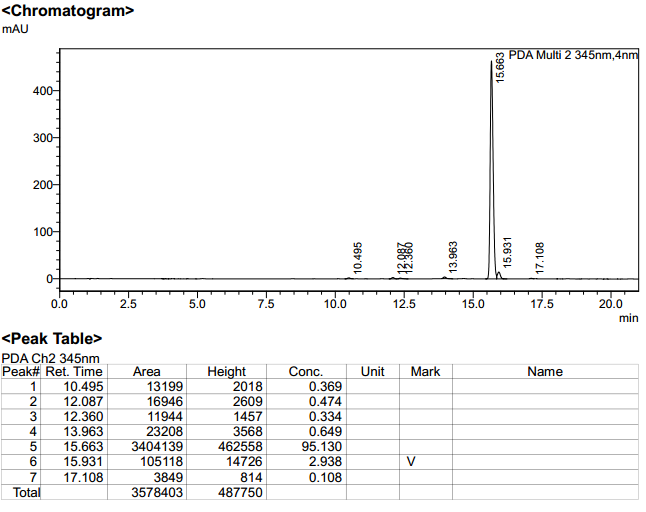

Figure S40. Integrated HPLC profile of compound **27bx** (**23**) with UV detection at 345nm.

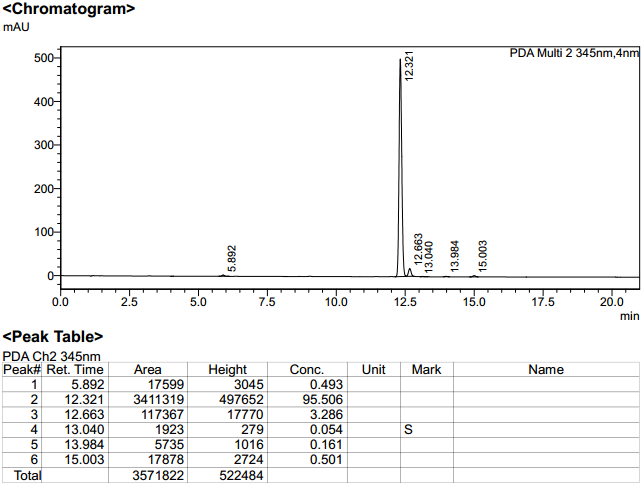

Figure S41. Integrated HPLC profile of compound **30x** (**22**) with UV detection at 345nm.

4.	Petrosiute, A., A. Zaksauskas, and A. Luciunaite. Combination of carbonic anhydrase IX and CCR2 inhibition as a path to treat neuroblastoma. *Submitted*.
